# Fatty acid metabolism controls plasma membrane cholesterol accessibility via ATGL-dependent phospholipid remodeling

**DOI:** 10.1101/2025.08.21.671640

**Authors:** Kalani M. Wijesinghe, Chai-wan Kim, Summer Chen, Emily O. Schad, Erin Takeshima, Shuo Li, Chandni B. Khandwala, Dylan Calhoon, Megan Danielewicz, Ryan D. Leib, Javier Garcia-Bermudez, Desiree Tillo, Andres M. Lebensohn, James A. Olzmann, Rajat Rohatgi, Maia Kinnebrew

## Abstract

Accessible cholesterol, the pool of membrane cholesterol with high chemical activity, regulates vital processes including vertebrate development and pathogen evasion. The mechanisms that govern plasma membrane (PM) cholesterol accessibility are incompletely understood. Using a genome-wide screen we find that acetyl-CoA carboxylase alpha (ACC1) loss causes a ∼10-fold increase in PM accessible cholesterol in cells and a male mouse model. We demonstrate that reduced fatty acyl-CoA levels, achieved by targeting metabolic enzymes including ACC1, fatty acid synthase (FASN) or acyl-CoA synthetases long chain (ACSL1, 3 and 4), results in the activation of adipose triacylglycerol lipase (ATGL). ATGL activation elevates polyunsaturated diacylglycerols even in triacylglycerol-deficient backgrounds, suggesting that ATGL acts independently of triacylglycerol hydrolysis. These diacylglycerol species are utilized to generate polyunsaturated phosphatidylcholine and phosphatidylethanolamine, which raises PM fluidity and cholesterol accessibility. Conversely, ATGL inhibition rigidifies the PM, reducing PM accessible cholesterol. Increased PM fluidity impairs cholesterol transport, triggering SREBP2 activation. This study reveals a surprising link between fatty acid metabolism and cholesterol homeostasis, demonstrating how ATGL-dependent lipid remodeling tunes membrane fluidity to regulate intracellular cholesterol signaling.

## Introduction

Cholesterol constitutes approximately 40% of all lipids in the plasma membrane (PM) through the coordinated effort of nearly 100 proteins that regulate its synthesis, transport and metabolism^1,2^. While being vital for cell and organismal health, cholesterol is also linked to numerous human diseases, including cardiovascular disease, metabolic syndrome, cancer and neurodegeneration. Despite a rich history of investigation dating to the 1700s, we lack a complete understanding of how cholesterol is regulated. For example, only in the last 15 years has a pivotal concept come to light: while the majority of cholesterol plays structural roles in the PM, ∼2% of PM cholesterol has high chemical activity^3–7^. This pool of cholesterol, often referred to as *active* or *accessible* cholesterol, can dynamically bind to proteins and drive cell signaling events. The complete importance of accessible cholesterol in vivo is unknown, but thus far it has been shown to control de novo cholesterol synthesis, viral and bacterial pathogenesis, and the Hedgehog signaling pathway^8–13^.

The discovery and characterization of accessible cholesterol was made possible through the identification of a family of bacterial cholesterol-dependent cytolysin proteins such as Perfringolysin O (PFO) and Anthrolysin O (ALO), which specifically bind to accessible cholesterol^14–16^. The non-lytic cholesterol-binding domain 4 of PFO or ALO (PFOD4 or ALOD4) is an invaluable tool to interrogate the localization, function, and regulation of accessible cholesterol. For example, pioneering studies revealed that accessible cholesterol is continually transported from the PM to the endoplasmic reticulum (ER) where its levels are sensed to control de novo cholesterol synthesis^17–19^. When ER cholesterol is high, the membrane-tethered transcription factor Sterol Regulatory Element Binding Protein 2 (SREBP2) is retained in the ER and cholesterol synthesis is blocked. When ER cholesterol is low, SREBP2 is transported to the Golgi where it is proteolytically cleaved into a transcriptionally active fragment that drives the expression of low density lipoprotein receptor (LDLR), which facilitates low density lipoprotein (LDL) uptake, as well as genes involved in cholesterol biosynthesis (**Supplementary Fig. 1a**)^12,20–23^. A long-standing mystery is how changes in PM accessible cholesterol are conveyed to the cholesterol-sensing machinery in the ER. Additionally, are there proteins that sense and control the amount of PM accessible cholesterol directly?

The most well-characterized proteins that regulate PM accessible cholesterol levels are the Gram domain-containing proteins, GRAMD1A, GRAMD1B and GRAMD1C. These proteins localize in the ER and move cholesterol down its concentration gradient at PM-ER contact sites^24–27^. A triple knockout of GRAMD1A/GRAMD1B/GRAMD1C does not affect cell viability, demonstrating that additional pathways for cholesterol transport out of the PM must exist^28^. Other pathways such as cytoplasmically localized START-domain containing proteins or oxysterol binding proteins are highly redundant and thus, their relative contribution to PM cholesterol homeostasis has not been systematically investigated^29,30^. Finally, cholesterol can be internalized from the PM through vesicular transport, although there is no known endocytic pathway dedicated specifically to maintaining cholesterol homeostasis (**Supplementary Fig. 1b**)^31–34^.

The high accessible cholesterol content of vertebrate PMs has been exploited by both bacterial and viral pathogens for their entry into cells^10,13,35,36^. To counteract this, the immune system has evolved a mechanism to evade pathogens where PM accessible cholesterol levels are rapidly depleted. Interferon signaling induces the expression of the ER-localized enzyme cholesterol 25-hydroxylase (CH25H), which converts cholesterol to 25-hydroxycholesterol (25HC). 25HC diffuses across the PM to surrounding cells, triggering rapid PM cholesterol depletion by allosterically activating acyl-coenzyme A:cholesterol acyltransferase 1 and 2 (ACAT1 and ACAT2; genes *SOAT1* and *SOAT2*) at the ER. The ACAT1/2 proteins esterify cholesterol with a fatty acid, which is then stored in lipid droplets (LDs)^37–39^. How cholesterol is transported out of the PM in response to 25HC is unknown.

In this study we used a genome-wide screen to find regulators of PM accessible cholesterol in 25HC-treated cells. We find that ACC1, the rate-limiting enzyme in fatty acid biosynthesis, helps to maintain low PM accessible cholesterol; loss of ACC1 dramatically raises PM cholesterol accessibility both at baseline and in response to 25HC exposure. Mechanistically, we show that ACC1 loss depletes fatty acyl-CoA pools, which leads to the activation of ATGL, and ATGL is the critical downstream effector of the PM accessible cholesterol changes observed. ATGL activation leads to triacylglycerol (TAG) hydrolysis and LD catabolism, though we show that PM cholesterol changes occur independently of TAGs by creating a triple knockout cell line lacking the core TAG-synthesizing enzymes. Mass spectrometry instead reveals that, in the absence of TAGs, ATGL activation elevates polyunsaturated diacylglycerol (DAGs) species. Because DAGs are precursors used to make phosphatidylcholine (PC) and phosphatidylethanolamine (PE), activated ATGL elevates polyunsaturated PC and PE. Finally, polyunsaturated PC and PE raises PM fluidity and cholesterol accessibility; surprisingly, this elevated fluidity appears to restrict cholesterol movement out of the PM, leading to SREBP2 activation and elevated rates of cholesterol biosynthesis. Altogether, we find an unexpected link between fatty acyl-CoA availability and cholesterol metabolism, with broad implications for the field of lipid homeostasis.

## Results

### Random-insertion mutagenesis screens identify cholesterol regulatory genes

To test the known pathway of PM to ER cholesterol transport, we knocked out GRAMD1A, GRAMD1B and GRAMD1C (*GRAMD1-*TKO) in human haploid HAP1 cells (**Supplementary Fig. 1c**)^24–27^. Using the accessible cholesterol binding protein ALOD4 fused to mNeonGreen (mNG-ALOD4), we found that *GRAMD1*-TKO cells showed 1.2-fold elevated PM accessible cholesterol compared to wild-type (WT) cells, consistent with a defect in accessible cholesterol trafficking out of the PM (**Fig. 1a**). However, when treated with 25HC, accessible cholesterol was depleted from the PM with similar kinetics to WT cells (**Fig. 1b**). These results were indistinguishable to GRAMD1B single knockout cells (**Supplementary Fig. 1d**-**f**). 25HC can also reduce PM accessible cholesterol levels by activating liver X receptors (LXRs). LXR is a sterol-responsive transcription factor that increases the expression of ABC transporters that export cholesterol from the PM of the cell (**Supplementary Fig. 1b**)^40^. Overnight treatment with the LXR inhibitor GSK2033 strongly reduced the transcript levels of the ABC transporter ABCA1, and slightly elevated PM accessible cholesterol levels; however, it did not significantly block PM cholesterol depletion induced by 25HC (**Fig. 1c**, **d** and **Supplementary Fig. 1g**-**i**). These data demonstrate that when cells are stimulated with 25HC, accessible cholesterol can exit the PM in a GRAMD1A/B/C- and LXR-independent manner. This raises the question: how does cholesterol exit the PM in response to 25HC (**Fig. 1e**)?

**Figure 1.**
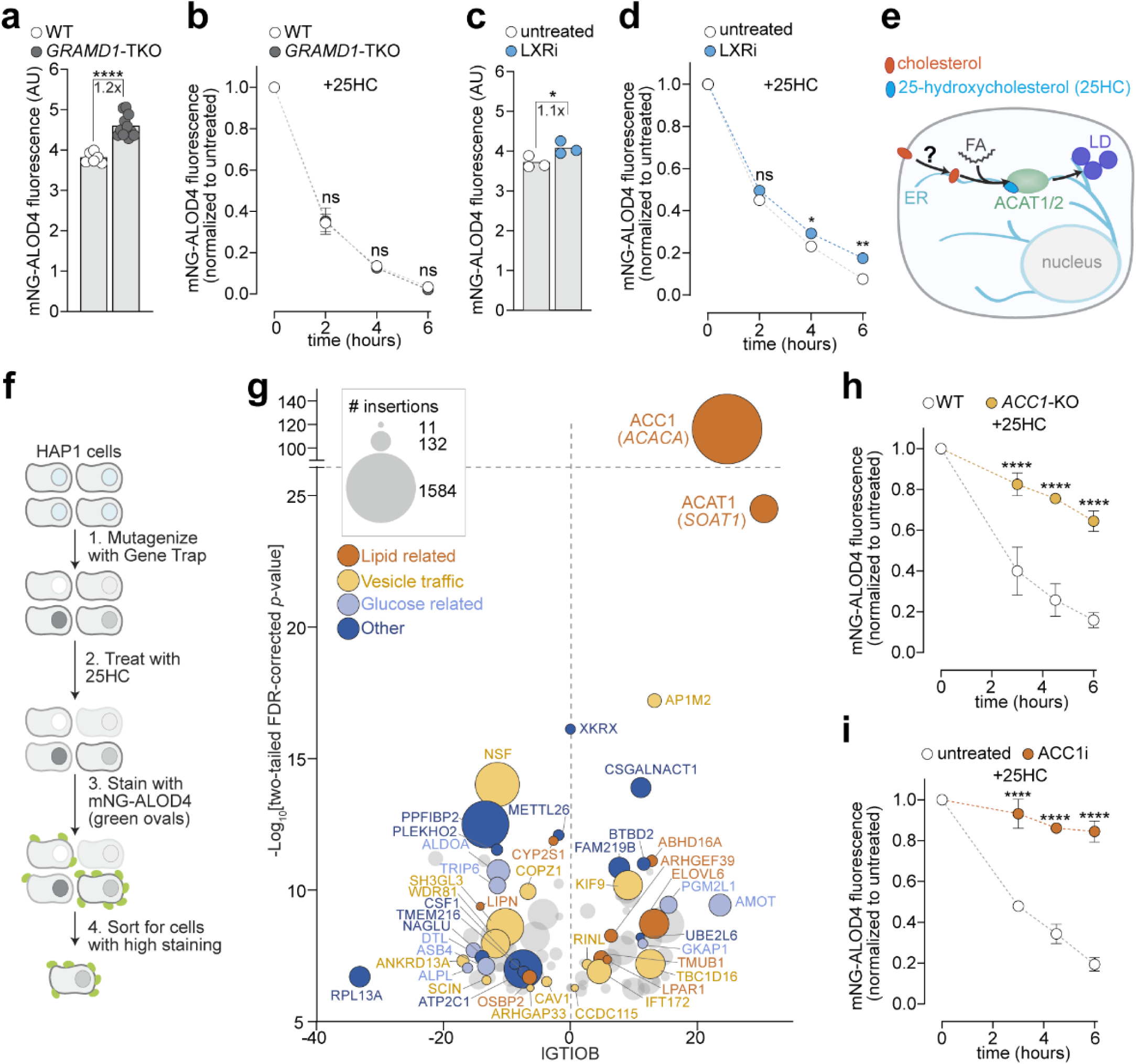
Random-insertion mutagenesis screens identify cholesterol regulatory genes, **a-d** PM cholesterol was measured by mNG-ALOD4 staining followed by flow cytometry. WT HAP1 cells are compared to *GRAMD1-TKO* either left untreated (a) or treated with 4 pM 25HC for the times indicated **(b).** WT cells were treated with or without LXR inhibitor (LXRi; 1 pM GSK2033) for 16 hrs and either left untreated **(c)** or treated with 25HC for the times indicated **(d).** Each circle is the median value for a biological replicate, with n=5,000 cells analyzed per replicate (a and c). e Schematic showing cholesterol trafficking from the PM to the ER membrane where it is esterified with a fatty acid (FA) by ACAT1/2; esterified cholesterol is stored in LDs. 25HC allosterically activates ACAT1/2, promoting this process, **f** Schematic of genetic screening pipeline: the mutant cell library was treated with 4 pM 25HC for 6 hrs and then cells were slained with mNG-ALOD4. The top 10% of fluorescent cells were isolated for sequencing, **g** Screen results showing enriched genes with an two-tailed FDR-corrected p-value of less than 2E-4. IGTIOB scores the inactivating potential of the mapped insertions, and circle size shows the number of insertions for each gene. Genes are colored based on their proposed functions. **h,i** mNG-ALOD4 flow cytometry analysis of PM cholesterol in cells left untreated **(h)** or treated for 16 hrs with an ACC1 inhibitor (ACC1i; 30 pM Firsocostat) **(i)** and then treated with 4 pM 25HC for the times indicated. Data is normalized to cells not treated with 25HC for each condition. Each circle is the mean of three biological replicates, with n=5,000 cells analyzed per replicate; error bars are the standard deviation **(b,d,** h and i), Statistical tests compare 1wo conditions at the same time point of 25HC treatment **(b,d, h** and **i).** Statistical significance was determined by two-tailed Welch’s t-test (a and **c)** or two-way ANOVA followed by Sidaks multiple comparisons test **(b,d,h** and i). Exact p-values are reported in the Methods. AU is an arbitrary unit.

To find genes that regulate PM cholesterol, we devised a genome-wide screening strategy. We reasoned that strong regulators of cellular cholesterol are likely to be required for cell viability, and therefore avoided CRISPR knockout screening. Instead, we used random-insertion mutagenesis to generate a library of ∼100 million cells in the HAP1 cell line. This mutational strategy can generate hypomorphic alleles by truncating proteins or by reducing the transcriptional output of genes. To induce accessible cholesterol exit from the PM we treated cells with 25HC for 6 hours, and then stained live cells at the outer leaflet of the PM with mNG-ALOD4. Finally, we sorted for cells with the top 10% of mNG-ALOD4 fluorescence to identify mutants in which cholesterol transport in response to 25HC was blocked (**Fig. 1f**)^17,19^.

Surprisingly, we identified acetyl-CoA carboxylase alpha (ACC1, gene named *ACACA*) as the top hit by a large margin (**Fig. 1g**). ACC1 catalyzes the rate limiting step in fatty acid biosynthesis, converting acetyl-CoA into malonyl-CoA, which is then elongated by fatty acid synthase (FASN) to produce long-chain saturated fatty acids^41^. Interestingly, the majority of ACC1 insertions enriched in the screen mapped to intron 52, which truncates ACC1 at its carboxyltransferase domain (**Supplementary Fig. 1j**). Using CRISPR to generate a series of ACC1 truncations, we found that an exon 52 truncation behaves identically to an exon 11 truncation lacking both catalytic domains. Furthermore, the exon 52 truncation was not detectable by Western blot and it was not responsive to the ACC1 inhibitor Firsocostat, a drug tested in human clinical trials for the treatment of metabolic dysfunction-associated steatotic liver disease (MASLD), demonstrating that it lacks protein function (**Supplementary Fig. 1k**-**m**). Throughout this manuscript, we utilize cell lines with ACC1 truncated at exon 52 for all assays, and we refer to these as *ACC1*-KO cells. Interestingly, we found that ACC1 truncated at exon 3 was lethal to cells; in support of this, previous work found that exons 2-4 of ACC1 form a circular RNA which is required for cell viability (**Supplementary Fig. 1n**)^42^.

In validation of the screen results, treatment of *ACC1*-KO cells with 25HC revealed a significant defect in cholesterol depletion compared to WT cells, with >60% of cholesterol remaining at the PM after 6 hours (**Fig. 1h**). The ACC1 inhibitor Firsocostat also prevented PM cholesterol depletion induced by 25HC in HAP1, 293T and 3T3 cells, demonstrating that this response is conserved across cell types from different lineages (**Fig.1i** and **Supplementary Fig. 1o**)^43^. ACC1 has been the subject of extensive research due to its role in human diseases, but it has not been linked to cholesterol homeostasis in cells, further motivating us to decipher this mystery.

Niemann-Pick type C1 protein (NPC1) is required to transport LDL-derived cholesterol out of the lysosome; mutations in NPC1 are the most common cause of Niemann-Pick disease type C, a fatal syndrome. We and others have shown that NPC1 inhibition or knockout causes reduced PM accessible cholesterol (**Supplementary Fig. 2a**-**d**)^44^. To identify a strategy to restore PM accessible cholesterol, we performed a second random insertion mutagenesis screen where cells were treated with the NPC1 inhibitor U18666A for 20 hours, then stained with mNG-ALOD4. Enrichment of the top 10% of mNG-ALOD4 fluorescent cells identified ACC1 as the top hit (**Supplementary Fig. 2e**). In validation of the screen, *ACC1*-KO cells treated with U18666A had PM cholesterol levels indistinguishable from WT untreated cells (**Supplementary Fig. 2a**). To more closely mimic a disease state, we treated either HAP1 cells or mouse embryonic fibroblasts lacking NPC1 (*NPC1*-KO) with the ACC1 inhibitor Firsocostat. In both cell types, ACC1 inhibition elevated PM accessible cholesterol (**Supplementary Fig. 2b, d**). These findings show that ACC1 inhibition does not raise PM accessible cholesterol levels through a mechanism involving NPC1, since it effectively raised PM accessible cholesterol in *NPC1*-KO cell lines.

### ACC1 functions independently of ACAT1/2 to maintain PM accessible cholesterol levels

Using both microscopy and flow cytometry, ACC1 loss caused a striking increase in PM accessible cholesterol in untreated cells (**Fig. 2a**, **b** and **Supplementary Fig. 1k**, **l**). This effect was observed in multiple cell types and with structurally distinct ACC1 inhibitors, including the dimerization inhibitor Firsocostat and the carboxyltransferase domain active site inhibitor CP-640186 (**Fig. 2c**, **d**)^45,46^. ACC1 loss also increased cell death caused by the cholesterol-dependent cytolysin PFO* (**Fig. 2e**). PFO* binds to accessible cholesterol and above 4°C it oligomerizes to generate pores in the PM^11,16^. We hypothesized that ACC1 could raise PM cholesterol accessibility by reducing sphingomyelin levels since sphingomyelin binds and sequesters cholesterol in the PM. Using mass spectrometry (MS), a sphingomyelin-binding probe, and a sphingomyelinase (SMase) assay, we showed that ACC1 loss does not reduce sphingomyelin levels (**Fig. 2f** and **Supplementary Fig. 3a**-**c**). Notably, ACC1 loss was able to raise PM accessible cholesterol to a similar degree as SMase treatment; this demonstrates that PM cholesterol accessibility can be dramatically altered through sphingomyelin-independent mechanisms.

**Figure 2.**
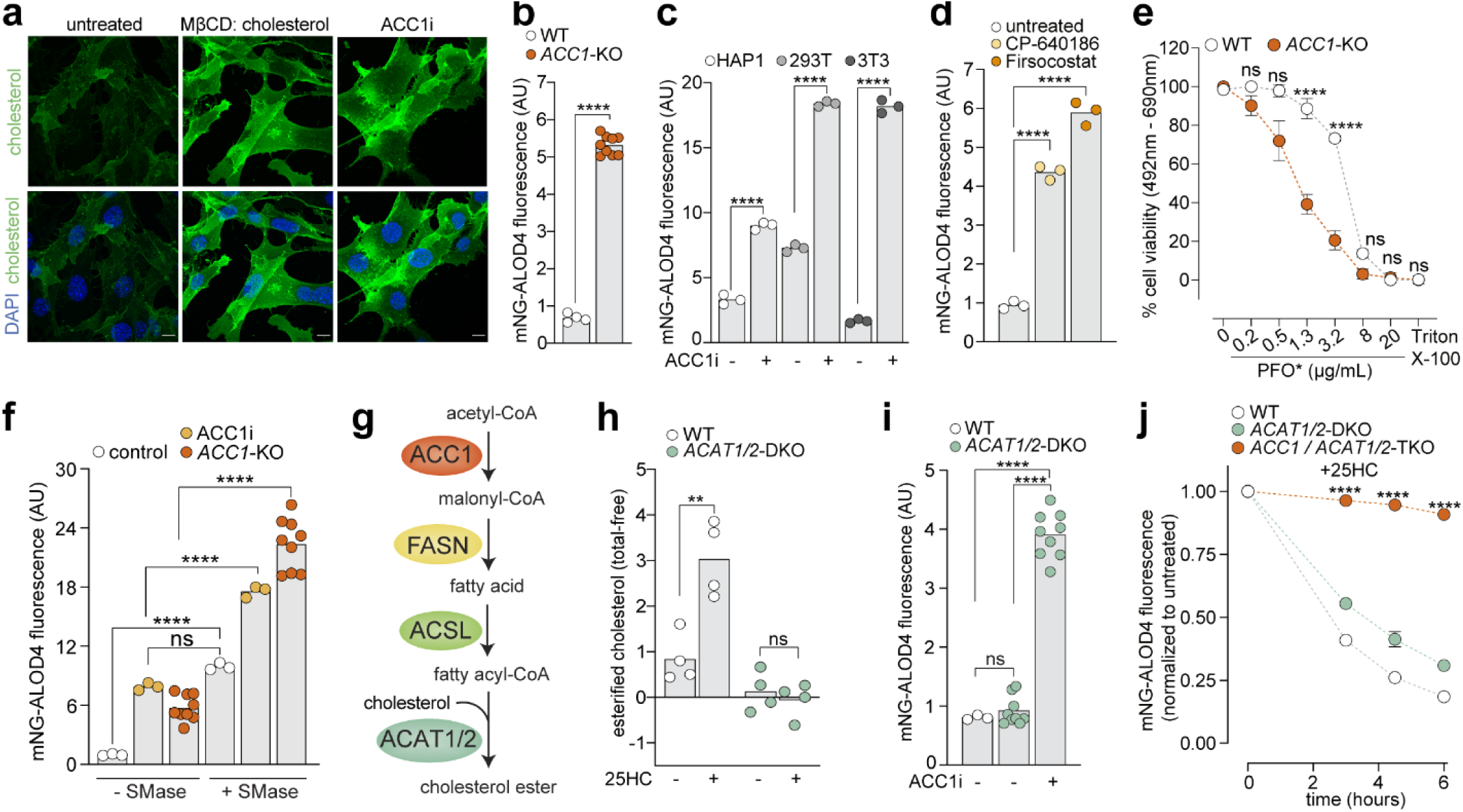
ACC1 functions independently of ACAT1/2 to maintain PM accessible cholesterol levels, **a** Fluorescence microscopy images showing WT 3T3 cells left untreated or treated with 0.3 mM M|3CD:cholesterol or ACC1i (30 pM Firsocostat) and stained with mNG-ALOD4 and DAPI; the scale bar is 10 microns, **b-d** Cell lines were left untreated **(b)** or treated with 30 pM Firsocostat (c) or 20 pM CP-640186 for 16 hrs **(d)** and then stained with mNG-ALOD4 and analyzed by flow cytometry, **e** Cells were treated with 4 pM 25HC for 4 hrs and then media was replaced with media containing PFO* for 30 minutes at 37°C before cell viability was measured with an XTT assay, **f** ACC1-KO cells or WT HAP1 cells were left untreated or treated for 16 hrs with ACC1i. Cells were then treated wllh or without 150 mU/mg SMase for 30 minutes and stained with mNG-ALOD4 and analyzed by flow cytomelry. **g** Schematic of cholesterol ester synthesis. ACAT1/2 transfer a fatty acid from acyl-CoAto cholesterol, forming cholesterol esters, h WT and ACATJ/2-DKO cell lines were treated with or wilhout 4 pM 25HC for 6 hrs and the esterified cholesterol was quantified using an Amplex Red Assay (n=4 biological replicates per sample, with each sample measured in triplicate), **i** mNG-ALOD4 flow cytometry analysis of PM cholesterol in WT and *ACAT1/2-DK.O* cells left untreated or treated with ACC1i for 16 hrs. Each circle is the median value for a biological replicate, with n=5,000 cells analyzed per replicate **(b-d,** f and **i). j** Cells were treated with 4 pM 25HC for the indicated times and then mNG-ALOD4 staining was measured by flow cytometry. Data is normalized to cells not treated with 25HC, and each data point represents the average of three biological replicates, with n=5.000 cells analyzed per replicate; error bars denole the standard deviation. Cells **(b, d,** and **i)** were grown in lipoprotein-depleted serum media for 16 hours prior to treatments. Statistical significance was determined by a two-tailed Welch’s 1-test **(b),** one-way ANOVA followed by Dunnett’s multiple comparisons test **(d),** two-way ANOVA followed by Sidak’s multiple comparisons test (c,e,f,h and **j),** or Brown-Forsythe and Welch ANOVA followed by Dunnett’s T3 multiple comparisons test **(i).** Exact p-values are reported in the Methods. AU is an arbitrary unit.

25HC-driven PM cholesterol depletion was previously shown to require ACAT1/2; these proteins esterify cholesterol with a fatty acid, promoting its storage in LDs^37^. We hypothesized that ACC1 loss reduces the pool of available fatty acids that are used by ACAT1/2 to make cholesterol esters (CEs) (**Fig. 2g**). If cholesterol cannot be esterified and stored, PM cholesterol may rise as observed in *ACC1*-KO cells. Indeed, *ACC1*-KO cells treated with 25HC did not show increased CE formation (**Supplementary Fig. 4a**,**b**). However, we speculated this could result from the inability of cholesterol to effectively exit the PM. To test if the inability to generate CEs results in elevated PM cholesterol we generated *ACAT1/2* double knockout (*ACAT1/2*-DKO) cells (**Supplementary Fig. 4c**). *ACAT1/2*-DKO cells showed undetectable levels of CEs at baseline and in response to 25HC (**Fig. 2h**). Importantly, and in clear opposition to the aforementioned model, PM accessible cholesterol levels were unaltered in *ACAT1/2*-DKO cells compared to WT cells^37^. Inhibition of ACC1 in *ACAT1/2*-DKO cells led to a dramatic increase in PM accessible cholesterol, demonstrating that ACC1 inhibition acts through an ACAT1/2-independent pathway (**Fig. 2i**).

If ACC1 and ACAT1/2 are acting in the same pathway to regulate 25HC-induced cholesterol depletion, then knocking out ACC1 and ACAT1/2 together should not have a combinatorial effect. We treated *ACAT1/2*-DKO cells with ACC1 inhibitor, which caused a dramatic increase in PM cholesterol in the presence of 25HC (**Supplementary Fig. 4d**). Additionally, ACAT1/2 inhibition with the inhibitor SZ58-035 further blunted the effect seen in *ACC1*-KO cells (**Supplementary Fig. 4e**-**g**)^47^. These findings were further supported by a triple knockout cell line lacking ACC1, ACAT1 and ACAT2 (**Fig. 2j** and **Supplementary Fig. 4h**). These epistasis experiments cleanly demonstrate that ACC1 acts through a distinct pathway from the ACAT1/2 proteins, both at baseline and in response to 25HC.

### ATGL activity controls PM cholesterol accessibility

ACC1 inhibition caused rapid changes to PM accessible cholesterol; this is consistent with changes in protein activity or trafficking, and it is inconsistent with the induction of a transcriptional program, which will show a lag as new genes are synthesized (**Fig. 3a**, **b**). We therefore tested whether PM accessible cholesterol changes were specific to ACC1 or more broadly caused by reduced fatty acid levels by knocking out FASN, the enzyme that functions downstream of ACC1 to generate fatty acids from malonyl-CoA. Similarly to ACC1 loss, FASN knockout (*FASN*-KO) cells showed a ∼2.6-fold increase in PM accessible cholesterol and a defect in cholesterol depletion from the PM in response to 25HC (**Supplementary Fig. 5a**-**c**). We noted that *FASN*-KO cells were notably sick and slow growing, even in the presence of 20% serum-containing media, which may contribute to their less dramatic phenotype. Nonetheless, the effects of *FASN*-KO are consistent with reduced fatty acids, or a downstream metabolite, leading to elevated PM accessible cholesterol.

**Figure 3.**
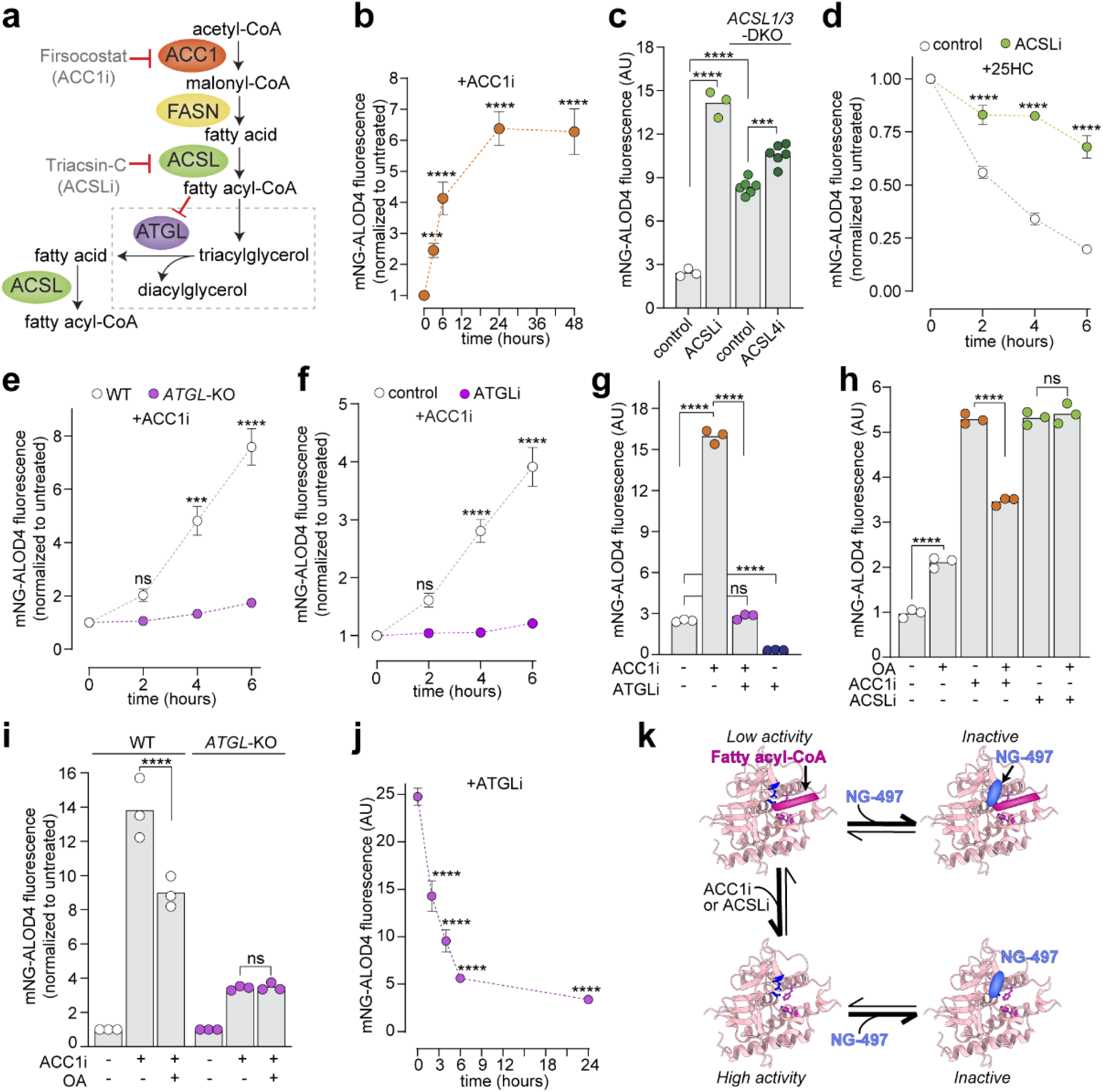
ATGL activity controls PM cholesterol accessibility, **a** Schematic showing the enzymes involved in the synthesis of fatty acyl-CoA. ATGL is allosterically inhibited by fatty acyl-CoA, which prevents it from hydrolyzing triacylglycerol (TAG) into diacylglycerol (DAG) and a fatty acid. Inhibitors targeting ACC1 and ACSL1,3,4 (ACSL) are indicated, **b-j** PM cholesterol was measured by mNG-ALOD4 staining followed by flow cytometry. All cells were grown in lipoprotein-depleted media for 16 hours prior to treatments, b WT HAP1 cells treated with ACC1i (30 pM Firsocostat) for the indicated times c WT and *ACSL* 7/3-DKO HAP1 cells were left untreated or treated with ACSLi (5 pM Triacsin-**C**) or ACSL4i (10 pM AS-252424) for 16 hrs. **d** WT HAP1 cells were treated with or without ACSLi for 16 hrs, followed by treatment with 4 pM 25HC for the indicated times, e WT or A7GL-KO HAP1 cells were treated with ACC1i for the indicated times, f WT cells were treated with or without ATGLi (50 pM NG-497) for 16 hours followed by treatment with ACC1i for the indicated times g WT HAP1 cells were treated with ACC1i. ATGLi, or both for 16 hrs h WT HAP1 cells were left untreated or treated with ACC1i or ACSLi for 16 hours, followed by treatment with oleic acid (OA, 200 pM) for 6 hours, **i** WT or ATGL-KO HAP1 cells were treated with or without ACC 1i for 16 hours followed by 6 hours of treatment with OA Each circle is the median value for a biological replicate, with n=5,000 cells analyzed per replicate (**c,g,h**, and **i**) j WT HAP1 cells were treated with ATGLi for the indicated limes, **k** Model schematic showing the activity of ATGL is determined by fatty acyl-CoAand NG-497 binding; light pink shows the Alphafold 3 structure ofATGL’s patatin domain, with catalytic residues (S47 and D166) shown in blue stick format. In b and d-f values were normalized to the untreated control at the corresponding time point; in b, **d-f** and **j** each data point represents the average of three biological replicates with n=5,000 cells analyzed per replicate; error bars denote the standard deviation. Statistical significance was determined by one-way ANOVAfollowed by Dunnett’s multiple comparisons test (b,j), two-way ANOVA followed by Sidak’s multiple comparisons test (c-f and i) or one-way ANOVA followed by Sidak’s multiple comparisons test (g,h). Exact p-values are reported in the Methods. AU is an arbitrary unit

Five acyl coenzyme A synthetase long-chain proteins (ACSL1 and ACSL3-6, henceforward called ACSL) function downstream of FASN to charge fatty acids with coenzyme A (CoA), generating fatty acyl-CoAs (**Fig. 3a**). To test the involvement of fatty acyl-CoA levels in the maintenance of PM cholesterol, we treated cells with Triacsin-C, an inhibitor of ACSL1, 3 and 4, and also made a CRISPR double knockout of ACSL1 and ACSL3 (*ACSL1/3*-DKO)^48,49^. Strikingly, both pharmacological and genetic ablation of ACSL caused elevated PM accessible cholesterol (**Fig. 3c** and **Supplementary Fig. 5d**). Additionally, ACSL inhibition also blocked PM accessible cholesterol depletion in response to 25HC (**Fig. 3d**).

Given that ACSL loss phenocopied ACC1 loss, we hypothesized that their common metabolic product, fatty acyl-CoA, modulates PM cholesterol accessibility. Adipose triacylglycerol lipase (ATGL, encoded by the gene *PNPLA2*) acts as a metabolic sensor for fatty acyl-CoAs. These molecules allosterically inhibit ATGL; their depletion thus elevates ATGL activity, resulting in the hydrolysis of triacylglycerol (TAG) into diacylglycerol (DAG) and a fatty acid, which restores fatty acyl-CoA pools (**Fig. 3a**)^50^. If the PM cholesterol phenotype was a direct consequence of fatty acyl-CoA depletion, an ATGL knockout (*ATGL*-KO) should exacerbate the effect by preventing fatty acid scavenging. Strikingly, we observed the opposite: both genetic deletion of ATGL and pharmacological inhibition with the human-specific ATGL inhibitor NG-497 blocked the elevation of PM accessible cholesterol induced by ACC1 or ACSL loss (**Fig. 3e**-**g** and **Supplementary Fig. 5e**,**f**)^51^. These results indicate that the phenotype is not caused by a reduction in fatty acyl-CoAs themselves, but rather by the subsequent activation of ATGL in response to low fatty acyl-CoA levels.

To confirm that ATGL is the critical effector downstream of ACC1, we attempted to rescue the ACC1-inhibited cells by restoring fatty acyl-CoA levels. Exogenously added palmitic or oleic acid successfully diminished the ACC1 phenotype. Additionally, as expected, fatty acids failed to rescue ACSL-inhibited cells because ACSLs are required to convert fatty acids into fatty acyl-CoAs (**Fig. 3h** and **Supplementary Fig. 5g**). Crucially, fatty acids failed to reverse the ACC1 phenotype in *ATGL*-KO cells, showing that in the absence of ATGL, fatty acyl-CoA levels are no longer sensed (**Fig. 3i**). Together, these data provide evidence that ACC1 inhibition lowers fatty acyl-CoA levels and activates ATGL to elevate PM accessible cholesterol.

While ATGL activation increased PM cholesterol accessibility, we found that ATGL inhibition caused the opposite effect. Treatment with the ATGL inhibitor NG-497 resulted in a rapid, ∼5-fold reduction in PM accessible cholesterol, a temporal profile that mirrored the kinetics of ACC1 inhibition but in the reverse direction (**Fig. 3g** and **3j**). This suggests that ATGL serves as a rheostat for PM accessible cholesterol: low fatty acyl-CoA levels allosterically activate ATGL to increase accessible cholesterol, and competitive inhibition of ATGL via NG-497 diminishes accessible cholesterol (**Fig. 3k**).

The dynamic nature of this process was further demonstrated through the observation that ATGL inhibition with NG-497 was able to reverse the elevated PM cholesterol levels in cells pre-treated with ACC1 inhibitor (**Supplementary Fig. 5h**). This also shows that ACC1 loss does not cause irreversible changes to the PM lipid composition; instead, it further argues that ATGL is the critical downstream determinant of the PM cholesterol changes we observe. Interestingly, we did not observe a reduction in accessible cholesterol in *ATGL*-KO cells, suggesting the presence of compensatory mechanisms that adapt to long-term ATGL deficiency to maintain cholesterol homeostasis. Collectively, these data indicate that ATGL activity is a key mediator of PM cholesterol accessibility; its activation status can be dynamically tuned to modulate the membrane landscape.

### ACC1 inhibition triggers ATGL activation and triacylglycerol hydrolysis

ATGL activation triggers the hydrolysis of TAG and thus the catabolism of LDs. ACC1 has primarily been viewed as a driver of LD biogenesis through its role in de novo lipogenesis rather than a regulator of LD breakdown via ATGL (**Fig. 4a**). We reasoned that if ACC1 inhibition indeed triggers ATGL activation, it should accelerate the catabolism of LDs. Supporting a localized role in this process, we found that ACC1 is enriched at the LD surface, as confirmed by both microscopy and density gradient ultracentrifugation (**Fig. 4b, c**)^52,53^. Consistent with ATGL activation, ACC1 inhibition caused the rapid loss of LDs, which inversely correlated with an increase in PM cholesterol accessibility (**Fig. 4d**)^54^. Overnight treatment with the ACC1 inhibitor Firsocostat decreased the median number of LDs per cell from ∼six to zero and *ACC1*-KO cells lacked LDs (**Supplementary Fig. 6a**, **b**). Interestingly, FASN inhibition with the small molecule C75 did not alter LD number or PM accessible cholesterol after 48 hours, and FASN also did not localize at LDs in these cells (**Fig. 4e** and **Supplementary Fig. 6c**). *FASN*-KO cells, in contrast, showed reduced LD numbers and elevated PM accessible cholesterol; this suggests that FASN inhibition results in LD loss on a slower timescale (**Supplementary Fig. 5b** and **6d**). Similarly to ACC1 inhibition, ACSL inhibition with Triacsin-C raised PM accessible cholesterol and triggered LD loss (**Fig. 4f**).

**Figure 4.**
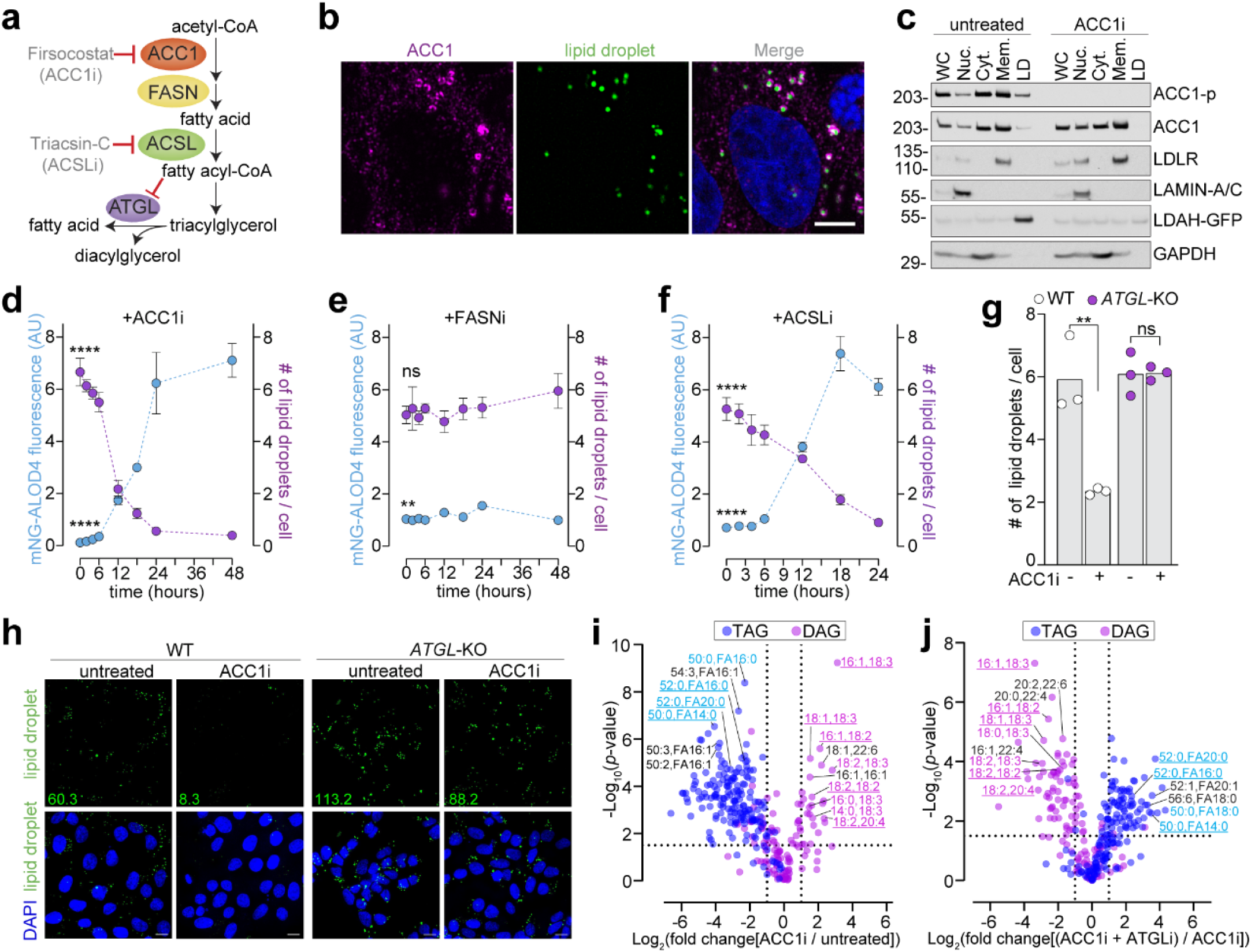
ACC1 inhibition triggers ATGL activation and triacylglycerol hydrolysis, **a** Schematic showing the synthesis of fatty acyl CoA from acetyl-CoA, inhibition of ACC1 or ACSL reduces fatty acyl-CoA levels, triggering ATGL activation, **b** Microscopy images of HAP1 cells stained with an ACC1 monoclonal antibody, BODIPY 493/503 to measure LDs, and DAPI; the scale bar is 5 microns, **c** Western blot analysis of fractionated HAP1 cells expressing the LD-localized protein LDAH fused to GFP (LDAH-GFP) with various cellular fractions including whole cell (WC), nucleus (Nuc.; detected by LaminA/C), cytoplasm (Cyt.; detected by GAPDH), cell membranes (Mem.; detected by LDLR), and LDs (detected by LDAH-GFP). Cells were either left untreated or treated with ACCIi (30 pM Firsocostat) for 24 hrs.The experiment was repeated independently three times with similar results **d-f** mNG-ALOD4 flow cytometry analysis of PM cholesterol (blue curves) and the corresponding fluorescence microscopy quantification of the number of LDs per cell (purple curves) in HAP1 cells grown in 5% fetal bovine serum and then treated with ACCIi (**d**), FASNi (10 pM C75) (e) or ACSLi (5 pM Triacsin-C) (**f**). Each data point represents the average of three biological replicates, with error bars denoting the standard deviation. g,h Quantification of LD numbers per cell (g) and corresponding microscopy images (**h**) in WT and ATGL-KO HAP1 cells treated with or without ACCIi for 6 hrs. Cells were stained with BODIPY 493/503 and DAPI; the scale bar is 5 microns. Green numbers in the LD field show the total LD area in μm^2^. For (**g**), n=75 cells examined per field of view across 3 fields of view. Each data point represents the total number of LDs per cell in a single field of view. **i.j** Volcano plot comparing lipid species in ACC1i-trea1ed versus untreated (**i**) or ACC1i-treated versus ACCIi and ATGLi (50 pM NG-497) combination treatment. Triacylglycerols (TAGs, blue circles) and diacylglycerols (DAGs, magenta circles) are shown with log_2_(fold-change) plotted against -log,_0_(p-value). The identity of select lipid species are shown, where underlined lipids are those found in both i and j, pink DAG names conlain either 18:2 or 18:3 acyl chains and blue TAG names are fully saturated. Dashed lines are arbitrary thresholds for fold-change and statistical significance, meant to highlight lipids that are altered across conditions. Statistical significance was determined by one-way ANOVA followed by Dunnett’s multiple comparisons test (**d-f**) or two-way ANOVA followed by Sidak’s multiple comparisons test (**g**). Exact p-values are reported in the Methods. AU is an arbitrary unit.

To determine whether LD depletion stemmed from impaired fatty acid synthesis or accelerated catabolism, we compared the kinetics of LD loss induced by ACC1 inhibition to known LD synthesis inhibitors. Blocking the activity of the TAG-synthesizing enzymes diacylglycerol O-acyltransferase 1 and 2 (DGAT1 and DGAT2) did not change LD numbers after 24 hours (**Supplementary Fig. 6e**). These drug treatments were robust since they blocked new LD formation induced by exogenous oleic acid addition. In contrast, ACC1 inhibition triggered a 50% reduction in LDs within only 10 hours (**Fig. 4d**). This depletion was further accelerated, reaching 50% loss in 6 hours, when cells were cultured in LDL-deficient media, likely because lipoprotein starvation further lowers fatty acyl-CoA levels to enhance ATGL activation (**Supplementary Fig. 6f**). These data provide evidence that ACC1 inhibition does not merely block LD formation but actively triggers ATGL-mediated LD catabolism.

Most compellingly, *ATGL*-KO cells were resistant to LD loss triggered by ACC1 inhibition; this demonstrates that in the absence of ATGL, ACC1 inhibition does not reduce LD numbers (**Fig. 4g**, **h**). To measure TAG hydrolysis through a more direct mechanism, we developed a targeted mass spectrometry pipeline to specifically measure TAGs and DAGs. ACC1 inhibition caused a dramatic reduction in TAG abundance, and a corresponding increase in DAG levels, and this was reversed by blocking ATGL (**Fig. 4i**, **j**). These data demonstrate that ACC1 inhibition triggers ATGL activation, leading to reduced TAG levels and LD loss.

To modulate ATGL activity in an orthogonal way, we targeted AMP-activated protein kinase (AMPK), a master regulator of cellular energy metabolism. AMPK regulates the activity of both ATGL and ACC1 through an activating and inactivating phosphorylation, respectively (**Supplementary Fig. 7a**). Consistent with our previous data, inhibition of AMPK with the drug Dorsomorphin (also called Compound C), which is expected to inhibit ATGL and activate ACC1, elevated LD numbers and lowered PM accessible cholesterol (**SupplementaryFig. 7b-g**). Testing additional structurally distinctive AMPK inhibitors lowered PM accessible cholesterol to various extents (**Supplementary Fig. h-l**). However, this effect was not recapitulated in cells where the AMPK isoforms AMPK1 and AMPK2 were genetically ablated with siRNA and CRISPR knockout, respectively. Given the variability of this assay (and the reliance on small molecule inhibitors), we did not further explore this line of investigation.

### ATGL modulates PM cholesterol accessibility in cells lacking LDs

Although LD depletion consistently correlated with increased PM cholesterol accessibility, it remained unclear whether LDs were regulatory determinants of membrane cholesterol. To decouple LD abundance from ATGL activity, we knocked out the three enzymes known to synthesize TAG: DGAT1, DGAT2, and DGAT1/2- Independent Enzyme Synthesizing Lipid Storage (DIESL) (here forward referred to as TAG-TKO) (**Fig. 5a** and **Supplementary Fig. 8a**). As expected, TAG-TKO cells lacked LDs, and no LDs were formed with either ATGL inhibition or exogenous fatty acid addition (**Fig. 5b**,**c** and **Supplementary Fig. 8b**).

**Figure 5.**
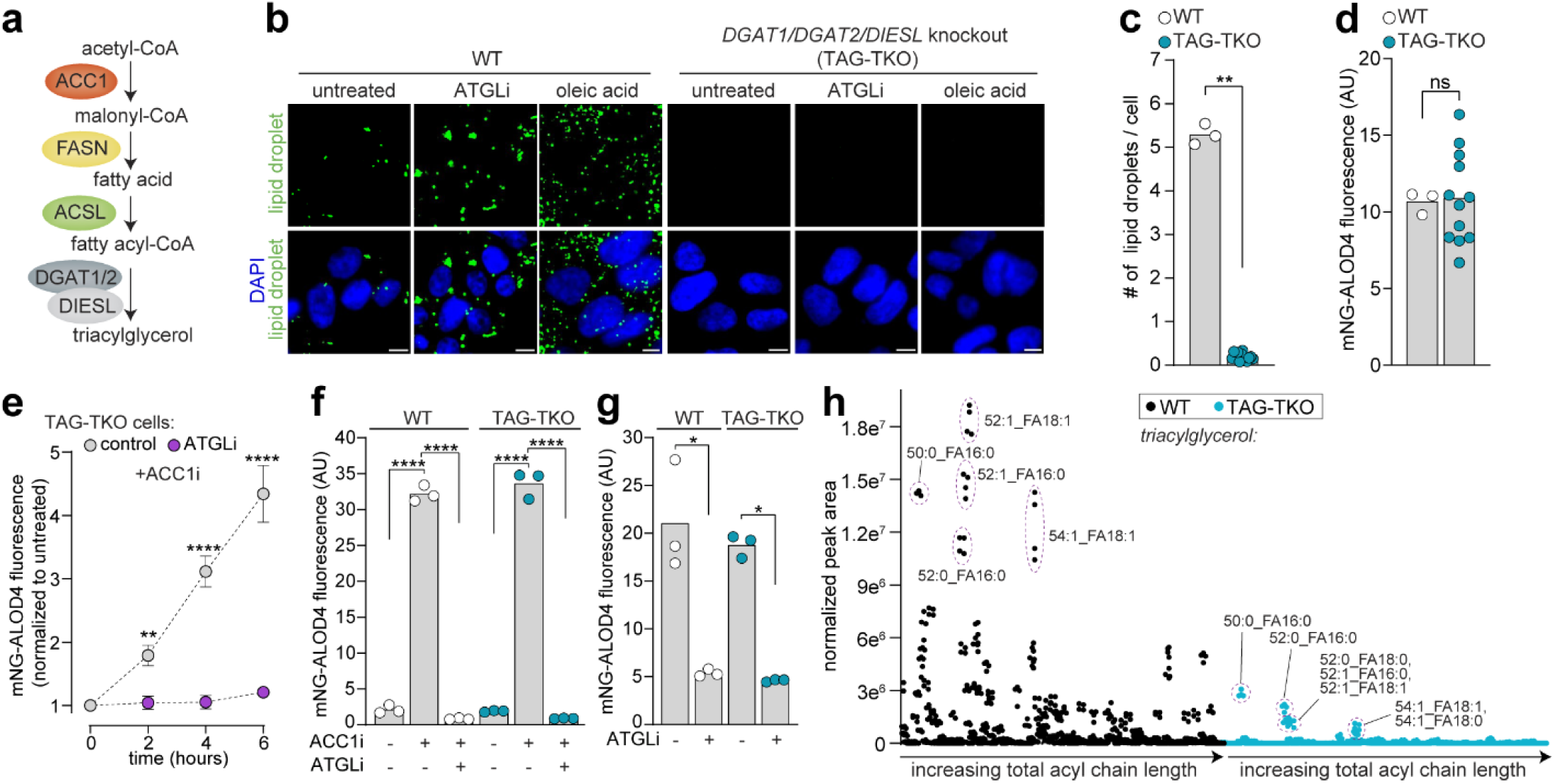
ATGL modulates PM cholesterol accessibility in cells lacking LDs. **a** Schematic of the TAG synthesis pathway highlighting TAG formation mediated by DGAT1, DGAT2 (DGAT1/2) and DIESL. **b,c** Microscopy images (b) and quantification (c) of LDs in WT and TAG-TKO HAP1 cells left untreated (b,c) or treated with ATGLi (50 pM NG-497) for 16 hrs or 200 pM oleic acid for 6 hrs (b). Cells were slained with BODIPY 493/503 and DAPI; the scale bar is 5 microns. Each data point represents the total number of LDs per cell in a single field of view, with 75 cells examined per field across 3 independent fields of view (c) d-g mNG-ALOD4 flow cytometry analysis of PM cholesterol in WT and TAG-TKO cells left untreated **(d)** or treated with ATGLi for 16 hrs (e-g) followed by ACC1i-treatment (30 pM Firsocostat) for the indicated times **(e)** or 1S hrs **(f).** Each circle is the median value for a biological replicate, with n=5,000 cells analyzed per replicate **(d,f** and g); in (e), each data point represents the average of three biological replicates with n=5000 cells analyzed per replicate; error bars denote the standard deviation, **h** MS analysis showing all detected TAG species in WT and TAG-TKO cells plotted with approximate increasing acyl chain length and unsaluration from left to right (x-axis). Selected TAG species are annotated; the ionized fatty acid (FA) is shown. Statistical significance was determined by a two-tailed Welch’s t-test **(c,d)** or two-way ANOVA followed by Sidak’s multiple comparisons test (e-g). Exact p-values are reported in the Methods. AU is an arbitrary unit.

Strikingly, TAG-TKO cells had indistinguishable PM cholesterol accessibility compared to WT cells (**Fig. 5d**). Furthermore, ACC1 inhibition still elicited a dramatic increase in PM accessible cholesterol in TAG-TKO cells, which remained sensitive to both ATGL inhibition (NG-497) and fatty acid rescue (**Fig. 5e**, **f**; **Supplementary Fig. 8c**). Acute ATGL inhibition in these LD-deficient cells caused a reduction in PM accessible cholesterol, mirroring the effects seen in WT cells (**Fig. 5g**). These data indicate that ATGL-mediated regulation of PM cholesterol is LD-independent.

While LDs were not observed in TAG-TKO cells, we sought to quantify TAG species using lipidomics. Surprisingly, while total TAG levels were dramatically reduced, these cells showed low-level but clearly detectable levels of TAG species that were either fully saturated or monounsaturated (**Fig. 5h**). Prior to lipidomics, these cells were cultured in lipoprotein-depleted media for 48 hours, and we do not believe that these TAGs originated from lipoproteins. This suggests there may exist additional pathways to make TAGs, though they are not responsive to exogenous fatty acid addition. Additionally, the TAG abundance does not exceed the threshold needed to nucleate LD formation, which was previously determined to be about 3 mole % of membrane lipids^55^. Interestingly, these residual TAG pools remained responsive to ATGL, as their levels were depleted upon ATGL activation and elevated upon ATGL inhibition (**Supplementary Fig. 8d,e**). Altogether, these data provide convincing evidence that PM accessible cholesterol is not sensitive to LD abundance.

### ATGL activation raises phospholipid acyl chain unsaturation and PM fluidity

We next investigated how ATGL activity modulates PM cholesterol in the absence of LDs. Lipidomics revealed that the total DAG levels were largely unaltered in TAG-TKO cells compared to WT cells, suggesting that TAG hydrolysis plays a minor role in maintenance of the bulk DAG pool (**Fig. 6a**). Interestingly, ATGL activation (via ACC1 inhibition) specifically enriched the pool of polyunsaturated DAGs, particularly those containing linolenic (18:2), alpha-linolenic (18:3), or eicosatrienoic (20:3) acid chains, while simultaneously depleting saturated and monounsaturated species (**Fig. 6b**). Conversely, ATGL inhibition yielded a mirror-image profile, reducing polyunsaturated DAGs and increasing saturated/monounsaturated species (**Fig. 6c**). The remarkable reciprocity between ACC1 and ATGL inhibition across identical DAG species (e.g., 16:1/18:3 and 18:1/18:3) further validates that ACC1 inhibition triggers ATGL activation.

**Figure 6.**
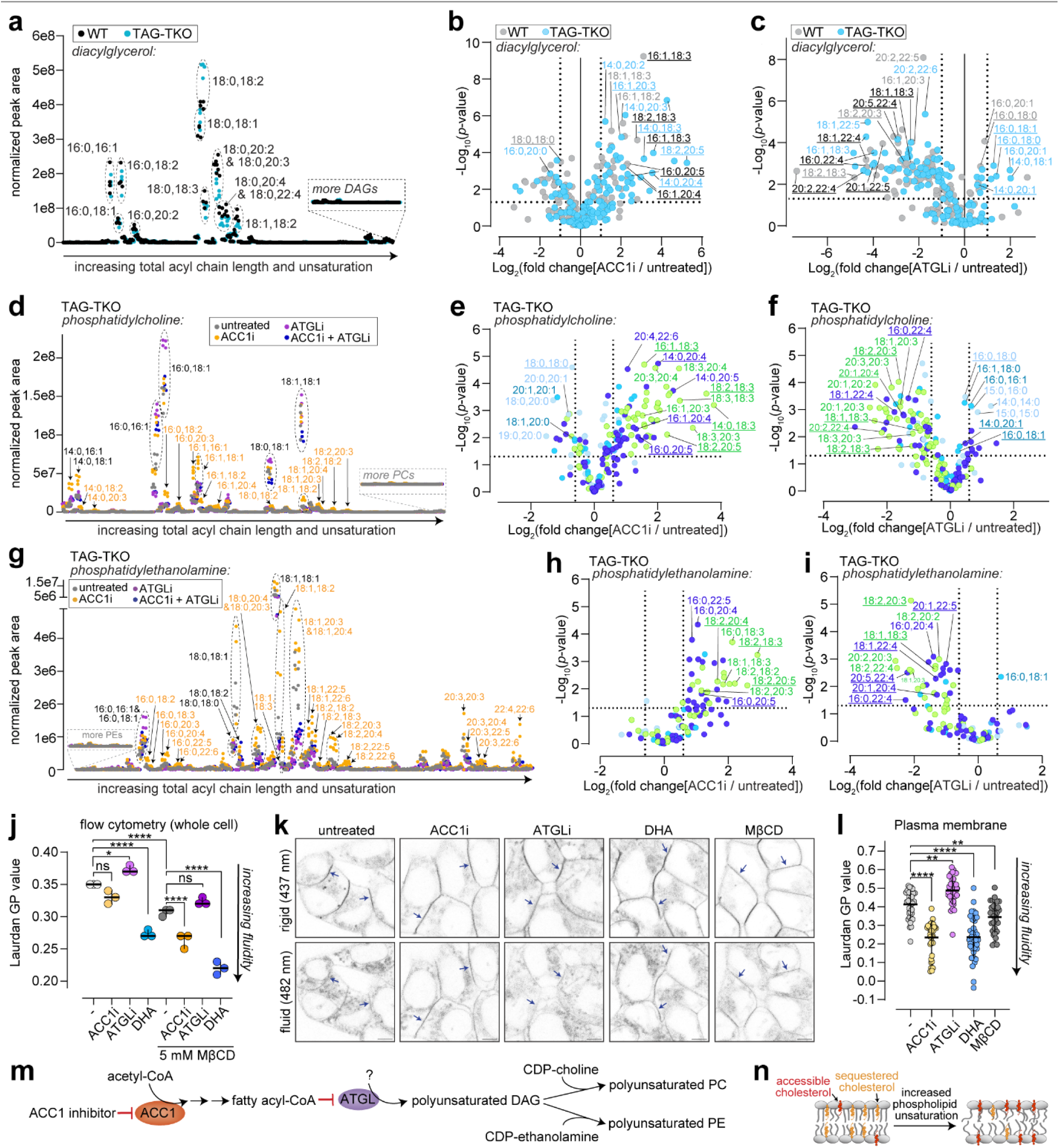
ATGL activation raises phospholipid acyl chain unsaturation and PM fluidity, **a** MS analysis showing DAG normalized peak areas plotted against increasing total acyl chain length and unsaturation for WT and TAG-TKO cells. High-abundance DAG species are annotated. **b,c** Volcano plots showing the log_2_(fold change) for DAG species in both WT and TAG-TKO cells; fold changes compare ACC 1 inhibitor (ACC1i; 30 pM Firsocostat) to untreated **(b)** or ATGL inhibitor (ATGLi; 50 pM NG-497) to untreated **(c).** Each point represents an individual lipid species annotated by acyl chain composition. Black annotations are lipid species found in both WT and TAG-TKO cells, and underlined lipid species are those whose acyl chain compositions match those found in phosphatidylcholine (PC) or phosphatidylethanolamine (PE) lipids in the subsequent figures, **d-i** MS analysis of PC **(d-f)** and PE (g-i) species. Plots show either the normalized peak area for the indicated treatments **(d** and **g)** or the log (fold change) versus the -log_10_(p-value) for indicated treatments **(e,f** and **h,i).** Vertical and horizontal dotted lines indicate arbitrary significance thresholds. Underlined lipid species are those whose acyl chains were identical to DAGs in **b** and **c. j** Whole cell membrane fluidity measured by Laurdan generalized polarization (GP) values for cells left untreated or treated with ACC1i, ATGLi or docosahexaenoic acid (DHA; 200 pM) for 16 hours, with or without a 20 minute treatment with 5 mM M(3CD to extract cholesterol. Each circle is the median value for a biological replicate, with n=5,000 cells analyzed per replicate; the error bar denotes the 95% confidence interval around the median, **k** Confocal images showing Laurdan fluorescence in both rigid (437 nm) and fluid (482 nm) emission wavelengths; arrows highlight visible PM regions; the scale bar is 5 microns. I Quantification of PM Laurdan GP values. Each data point represents the GP value measured from a PM segment of an individual cell. n=34 for untreated, n=37 for ACC1i-treated, n=34 for ATGLi-treated, n=37 for DHA-treated, and n=31 for M3CD-treated cells examined across 3 independent fields of view per condition. The error bar shows the standard deviation from the mean, m Proposed model showing that ATGL activation is triggered by ACC1 inhibition (red inhibitory arrows), leading to either direct or indirect generation of polyunsaturated DAGs, which are shunted into PC and PE biosynthesis, **n** Schematic showing that phospholipid acyl chain saturation alters the accessibility of cholesterol in membranes. Statistical significance was determined by two-way ANOVA followed by Sidak’s multiple comparisons test (j) or one-way ANOVA followed by Dunnett’s multiple comparisons test **(I).** Exact p-values are reported in the Methods.

The majority of cellular DAG is located in the ER where it acts as a precursor for phosphatidylcholine (PC) and phosphatidylethanolamine (PE) biosynthesis through the Kennedy pathway. We therefore analyzed PC and PE acyl-chain composition and found that similarly to DAGs, acyl chain unsaturation was increased in response to ATGL activation, and decreased in response to ATGL inhibition (**Fig. 6d**-**i**). Visual inspection supported this precursor-product relationship, revealing a striking conservation of acyl-chain signatures between DAGs and the PC and PE lipids (**Fig. 6b,c,e,f,h**, and **i**, see underlined lipid species). Importantly, ACC1 inhibition did not cause a global metabolic collapse; major species like 18:1/18:1 PC and 16:0/18:1 PC remained unchanged (**Fig. 6d**). WT cells showed the same changes in PC and PE unsaturation compared to the TAG-TKO cells, providing evidence that ATGL regulates these lipid pools through a mechanism that is TAG-independent (**Supplementary Fig. 8f**, **g**).

The lipid remodeling observed in our assays was highly specific. Only DAG, PC and PE saturation mirrored the bidirectional effects of ATGL on PM cholesterol; other major phospholipid classes, including phosphatidylinositol (PI), phosphatidylglycerol (PG), and phosphatidylserine (PS), were not inversely modulated by ATGL activation and inhibition (**Supplementary Fig. 8h**-**j**). Additionally, lysophosphatidylinositol (LPI), lysophosphatidylglycerol (LPG), lysophosphatidylserine (LPS), sphingomyelin (SM), ceramide (CER), hexosylceramide (hex-CER), lactosylceramide (lac-CER) and cholesterol esters (CE) were not inversely modulated by ATGL activation and inhibition (**Supplementary Fig. 9a**-**k**). Lipidomics analysis revealed that both ACC1 and ATGL inhibition independently lowered PG levels, with the combination treatment showing the most dramatic reduction in PG. Conversely, lysophosphatidylcholine (LPC) levels were elevated by both treatments, and even more so by the inhibitor combination. Plasmalogens were also elevated by combined ACC1 and ATGL inhibitor treatment. Finally, lysophosphatidylethanolamine (LPE) abundance was elevated by ACC1 inhibition, and unaltered by ATGL inhibition. Altogether, the lipidomics data is consistent with DAG, PC and PE causing the PM cholesterol changes observed.

Previous work demonstrated that phospholipid saturation directly modulates cholesterol accessibility^15,56,57^. Intuitively, because cholesterol is a planar molecule, it makes van der Waals interactions more robustly with saturated fatty acyl chains. We thus hypothesized that ATGL activation elevates PM cholesterol accessibility by elevating PC/PE unsaturation. To measure phospholipid saturation through an orthogonal method we utilized the membrane fluidity probe Laurdan, which detects polarity (water penetrability) in the membrane as a proxy for lipid packing. Flow cytometry measurements of Laurdan spectral shifts revealed that ATGL activation strongly elevates membrane fluidity, and conversely ATGL inhibition elevates membrane rigidity (**Fig. 6j**). The fluidity changes were maintained after cholesterol extraction from the membrane with methyl-beta cyclodextrin (MβCD), indicating that the fluidity changes are a primary result of phospholipid unsaturation, and not due to changes in cholesterol. Finally, confocal microscopy confirmed that fluidity changes occur at the PM (**Fig. 6k, l**). These data support a model in which ACC1 inhibition activates ATGL to generate polyunsaturated DAGs, which are channeled into PC and PE to remodel PM fluidity and elevate cholesterol accessibility (**Fig. 6m, n**).

### ATGL activation traps accessible cholesterol at the PM and triggers de novo cholesterol synthesis in a male mouse model

Elevated cholesterol accessibility is commonly correlated with cholesterol extractability from the membrane. For example, pioneering experiments demonstrated that the accessible pool of cholesterol is extracted by MβCD faster than the sequestered pool^58,59^. The proposed high chemical activity of accessible cholesterol is what makes it an excellent signaling molecule– it can readily bind proteins and be transported around the cell. In contrast, we find that ACC1 inhibition (leading to ATGL activation) traps cholesterol at the PM, blocking it from exiting in response to 25HC (**Fig. 1**). We hypothesize that in highly fluid membranes, cholesterol is energetically sequestered to shield non-polar acyl chains from the aqueous environment, making its extraction energetically unfavorable (**Fig. 7a**, **b**). To test this, we treated cells with a dose curve of MβCD. Consistent with previous reports, reducing sphingomyelin levels raised PM cholesterol accessibility, and also elevated its extractability by MβCD (**Fig. 7c**). Control cells showed 50% extraction (Ex_50_) at ∼0.7 mM M◻CD, and sphingomyelin-depleted cells showed an Ex_50_ of 0.45 mM. In contrast, ACC1 inhibition, which strongly raises PM cholesterol accessibility, shifted the Ex_50_ to ∼1.3 mM, suggesting that the accessible cholesterol observed in ACC1 inhibited cells is resistant to extraction (**Fig. 7c**). This suggests that high cholesterol accessibility does not inherently correlate with transport efficiency; rather, ATGL-mediated fluidization renders PM cholesterol resistant to mobilization.

**Figure 7.**
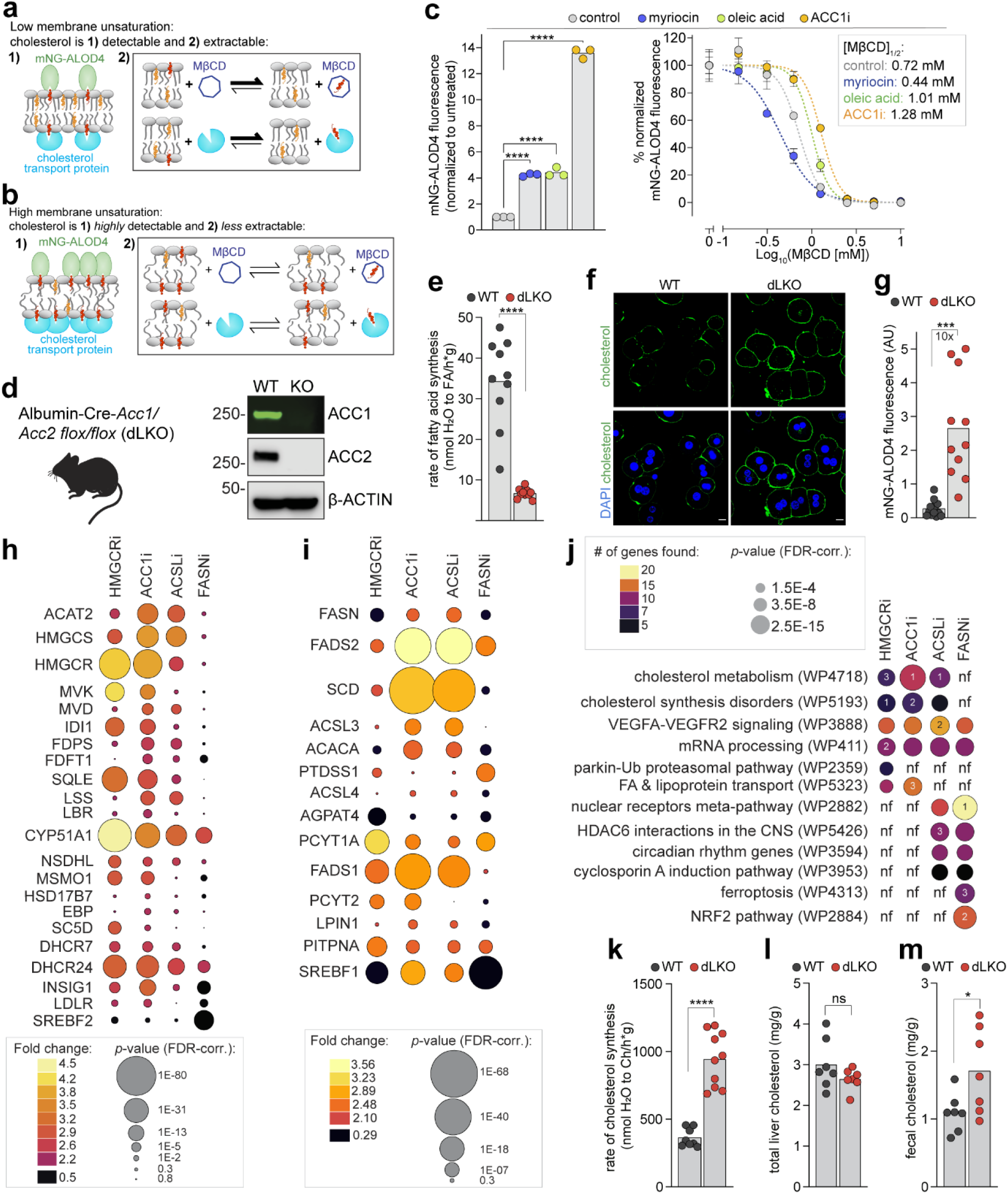
ATGL activation traps cholesterol at the PM and triggers de novo cholesterol synthesis in a male mouse model. **a,b** Model schematics showing that membranes with low phospholipid unsaturation have detectable and extractable cholesterol **(a).** In highly unsaturated membranes, cholesterol is highly detectable but is less efficiently extracted **(b). c** Flow cytometry analysis of PM cholesterol in WT HAP1 cells left untreated or treated with myriocin (30 pM), oleic acid (200 pM), or ACC1 i (30 pM Firsocostat) either alone (left) or followed by MpCD incubation for 20 min (right). Each circle is the median value for a biological replicate, with n=5,000 cells analyzed per replicate; the error bars show the standard deviation. The myriocin treatment was 72 hrs: oleic acid and ACCIi treatments were 16 hrs. The inset (right) shows the concentration at which MpCD extracted half of the accessible cholesterol for each treatment, **d** Western blot analysis of ACC1 and ACC2 proteins in primary hepatocytes isolated from male Acc1/Acc2 flox/flox control and Albumin-Cre-Acc1/Acc2 double liver knockout (dLKO) mice, e Hepatic fatly acid synthesis rates for male control and dLKO mice (n-10 mice per group). **f,g** Fluorescence microscopy images and quantification of primary hepatocytes stained with mNG-ALOD4 and DAPI: the scale bar is 10 microns, **h,i** Bubble plot showing differential gene expression from RNAseq in WT HAP1 cells pre-cultured in LDS media for 16 hours and then left untreated or treated for 6 hrs with HMGCRi (10 pM Lovastatin), ACCIi (30 pM Firsocostat), ACSLi (5 pM Triacsin-C) or FASNi (10 pM C75). Differential gene expression compares each treatment to the untreated control, **j** GSEA from RNAseq data using fhe top 200 genes sorted by FDR-corrected p-value. “nF is “not found” and the numbers inside of circles show the top three most enriched pathways for a given treatment, **k-m** Hepatic sterol synthesis rate **(k)** (n=10 mice per group), hepatic cholesterol content (I) (n=7 mice per group), or fecal cholesterol measurement **(m)** (n=7 mice per group) in chow-fed control and dLKO mice. Statistical significance was determined by one-way ANOVA followed by Dunnett s multiple-comparisons test **(c)** or a two-tailed Welch’s t-test **(e,g** and **k-m).** Exact p-values are reported in the Methods.

To strengthen the findings of this study, we extended our studies with an animal model. In hepatocytes, ACC2 contributes to the synthesis of ∼10% of cellular fatty acids^60^. Therefore, we induced recombination-mediated removal of both ACC1 and ACC2 using Albumin-CRE *Acc1/Acc2* flox/flox mice (with two flox sites between exons 21/22 and 26/27 for ACC1) to generate a double liver knockout (dLKO) of *Acc1/Acc2*^60^. Hepatocytes isolated from male *Acc1/Acc2* dLKO mice showed undetectable levels of ACC1 and ACC2 proteins, and fatty acid synthesis was markedly reduced (**Fig. 7d**, **e**). These hepatocytes also showed a reduction in TAG and CE levels compared to controls (**Supplementary Fig. 10a**, **b**). Strikingly, dLKO primary hepatocytes showed a 10-fold increase in PM accessible cholesterol, showing that the effects of ACC1 loss are maintained in vivo (**Fig. 7f**, **g** and **Supplementary Fig. 10c**,**d**).

Trapping cholesterol at the PM is expected to reduce ER-localized cholesterol and trigger de novo cholesterol biosynthesis. To assess the cellular changes that occur in response to ATGL activation, we performed RNA sequencing on cells treated with inhibitors against ACC1 and ACSL. As a comparator for the potent activation of cholesterol biosynthesis, we treated cells with Lovastatin, which blocks the rate limiting enzyme in cholesterol biosynthesis, HMG CoA reductase (HMGCR). We also included a FASN inhibitor treatment; this condition did not modulate PM cholesterol or ATGL activity in our assays (**Fig. 4e**), but was expected to reveal the cellular response to lowered fatty acid synthesis. Strikingly, we observed that ACC1 and ACSL inhibition closely resembled HMGCR inhibition and potently elevated the expression of cholesterol biosynthesis genes (**Fig. 7h**); in contrast, FASN inhibition had small effects, consistent with its apparent inability to activate ATGL (**Fig. 4e**, purple curve).

SREBP1 and SREBP2 are transcription factors that control the expression of genes involved in fatty acid and cholesterol biosynthesis, respectively. Both SREBP1 and SREBP2 are regulated by the ER-localized *cholesterol* sensor SREBP cleavage-activating protein (SCAP); consequently, low ER cholesterol levels trigger the simultaneous induction of both fatty acid and cholesterol biosynthetic pathways^22,61,62^. In contrast, we find that many SREBP1 target genes (FASN, FADS2, SCD, ACSL3, ACACA, and SREBP1 itself) were elevated more strongly through the inhibition of ACC1 or ACSL than by HMGCR (**Fig. 7i**). This was not due to the relative potency of different inhibitors since HMGCR inhibition had the strongest effects for multiple cholesterol biosynthesis genes (including HMGCR, MVK, SQLE and CYP51A1). Factors such as insulin and polyunsaturated fatty acids are known to regulate SREBP1 independently of SREBP2; future investigations are required to determine which of these are driving the discordance seen between SREBP2 and SREBP1 targets in our study^63–66^.

To determine which pathways were most significantly affected by the treatments that activate ATGL (ACC1 and ACSL inhibitors), we performed a Gene Set Enrichment Analysis (GSEA) using the top 200 most differentially regulated genes from each treatment condition. Among the pathways enriched, “Cholesterol Metabolism” was number one for both ACC1- and ACSL-inhibited cells (**Fig. 7j**). ACSL and FASN inhibition showed multiple overlapping enriched pathways unrelated to cholesterol metabolism. Additionally, all four inhibitors showed similar effects on “VEGFA-VEGFR2 Signaling” and “mRNA Processing” pathways, suggesting that blocking lipid synthesis (either cholesterol or fatty acid synthesis) induces comparable stress response pathways. Altogether, RNA sequencing provided strong evidence that ER cholesterol is depleted by ACC1 and ACSL inhibitor treatments, which supports the model that ATGL activation reduces cholesterol trafficking out of the PM.

Finally, we tested whether the rate of cholesterol synthesis was altered in the *Acc1/Acc2* dLKO primary hepatocytes. Consistent with our cellular findings, radiotracing of tritiated water showed dLKO hepatocytes had markedly elevated rates of cholesterol synthesis compared to controls (**Fig. 7k**). Despite having elevated PM cholesterol and elevated cholesterol synthesis, *total* cellular cholesterol levels were not significantly elevated in vivo or in cell culture (**Fig. 7l**, **Fig. 2i** and **Supplementary Fig. 10e**, **f**). Interestingly, dLKO mice had elevated fecal cholesterol levels, suggesting that in vivo homeostasis is maintained through cholesterol excretion (**Fig. 7m**). These findings suggest that there may be mechanisms to maintain total cellular cholesterol levels, which is at odds with the dogma that cells sense and regulate the accessible pool of cholesterol specifically. In conclusion, our study finds that ACC1 or ACSL inhibition activates ATGL. ATGL activation elevates PM phospholipid unsaturation, which raises membrane fluidity and cholesterol accessibility. Interestingly, this accessible cholesterol appears to be trapped at the PM, leading to the activation of de novo cholesterol biosynthesis.

## Discussion

In this study we set out to find regulators of cholesterol accessibility at the PM. We were surprised to discover that ACC1, a protein extensively studied for its role in fatty acid synthesis, also impacts cellular cholesterol homeostasis. ACC1 knockout raises PM accessible cholesterol levels 5- to 10-fold in cells and a mouse model; additionally, ACC1 loss blocks PM cholesterol depletion induced by 25HC through a mechanism that is independent of cholesterol esterification by ACAT1/2. Inhibition or knockout of the ACSL proteins, which charge fatty acids with CoA, phenocopies ACC1 loss, indicating that fatty acyl-CoA (the shared metabolic product of ACC1 and the ACSLs) is the relevant signal sensed by this pathway. Long-chain fatty acyl-CoAs were previously shown to allosterically inhibit ATGL in vitro^50^. In this study we used epistasis, pharmacology, and fatty-acid rescue experiments to demonstrate that ATGL senses fatty acyl-CoA abundance in cells; we propose that ATGL then acts as a rheostat, converting metabolic information into changes in the physical state of the plasma membrane.

Lipidomics data reveals that ATGL activation elevates polyunsaturated DAG abundance, as well as polyunsaturated PC and PE which we propose are made from DAG precursors. We hypothesize that ATGL generates polyunsaturated DAGs through a transacylation reaction for two reasons. First, ATGL was previously reported to have transacylase activity on DAG both in vitro and in vivo^67–70^. Secondly, we observed dramatic changes in PM cholesterol accessibility with *ACSL1/3*-DKO cells and ACSL inhibition. ACSL loss blocks the formation of fatty acyl-CoAs; therefore, the polyunsaturated DAG species must be made through a mechanism that does not require fatty acid charging with CoA. This rules out de novo lipid synthesis, and nominates either elevated desaturase activity or a transacylation reaction. Future work will be needed to determine how ATGL remodels the cellular DAG pool, and the consequences of this activity on cell and organismal biology.

Cholesterol accessibility is fundamentally a biophysical property governed by the strength of the interaction between cholesterol and the surrounding phospholipids and sphingolipids: saturated acyl chains sequester cholesterol efficiently, while polyunsaturated chains pack poorly and leave cholesterol more chemically exposed^71–73^. Consistent with this principle, we find that increased PC and PE unsaturation elevates PM fluidity and cholesterol accessibility. This relationship is well established in reconstituted systems such as liposomes and giant unilamellar vesicles, but has been difficult to test as a regulated, physiological process in cells. Our data suggests that a cell can dial PM cholesterol accessibility up or down by shifting the acyl-chain saturation of PC and PE, and that ATGL can drive this shift bidirectionally. To the best of our knowledge, this is the clearest demonstration that endogenous, physiologically triggered remodeling of bulk phospholipid saturation is sufficient to reset the chemical activity of PM cholesterol in intact cellular membranes.

The kinetics of PM accessible cholesterol changes occur on the timescales of minutes to a few hours after ATGL activation or inhibition (**Fig. 3b**, **j**), which is far faster than the transcriptional programs classically invoked for membrane adaptation. This pathway is thus mechanistically distinct from the transcriptional feedback loops that implement homeoviscous adaptation in response to temperature^74–76^. Here, membrane fluidity is instead tuned post-translationally on the timescale of metabolic fluctuation rather than gene expression.

Why would a cell couple fatty acyl-CoA supply to PM cholesterol trafficking? First, this pathway may coordinate the two major membrane lipid classes: our RNA-seq data shows that fatty acyl-CoA depletion drives SREBP2-dependent cholesterol biosynthesis. This suggests that when a cell is unable to build membranes from fatty acids, it may compensate by increasing sterol supply. Secondly, microbial infection in humans is well documented to reduce food intake, which has been shown to have both positive and negative consequences on the disease severity. Because PM accessible cholesterol governs susceptibility to pathogen entry, it is plausible that fatty acid starvation-induced changes in PM cholesterol accessibility have consequences for immune defense during nutrient stress. Further elucidation of these possibilities will require in vivo studies of fasting/refeeding and immune challenges.

While pioneering studies of accessible cholesterol defined it as the pool of cholesterol readily extractable by MβCD, we find that ALOD4 binding and cholesterol extractability are not equatable in polyunsaturated membranes. Our data show that these two readouts report distinct physical properties of cholesterol that are not required to track together. ALOD4 binding reflects cholesterol’s chemical activity at the membrane surface (its ability to be recognized by a cholesterol-binding protein), while MβCD or protein-mediated extraction instead reflects the kinetics of cholesterol dissociation into a small-molecule or protein acceptor. In support of the latter, our data show that polyunsaturated membranes trap cholesterol at the PM to a degree sufficient to trigger SREBP2 activation, indicating that extraction by intracellular cholesterol-binding proteins is impaired. This distinction has practical consequences for how “accessible cholesterol” is defined and measured across the field: an operational definition (cholesterol that binds ALOD4 or related probes) and a functional one (cholesterol that can be readily transported out of a membrane). Future work relating probe-based accessibility to actual transport and extraction rates across a range of defined membrane compositions will be needed to resolve which property governs cholesterol’s biological functions in signaling, transport, and disease.

## Methods

### Cell culture

HAP1 cells were grown in HyClone Isocove’s Modified Dulbecco’s Medium (IMDM) (SH30228FS, Fisher/Cytiva), supplemented with 5% Fetal Bovine Serum (FBS) (F0926, Millipore Sigma), 2 mM L-glutamine (MT25005CI, Fisher/corning), Penicillin (50 U/mL) and Streptomycin (50 μg/mL) (15140122, Fisher/Gibco). 3T3, 293T and MEF cells were cultured in HyClone™ Dulbecco’s Modified Eagles Medium (SH30081FS, Fisher/Cytiva), enriched with 10% FBS (F0926, Millipore Sigma), 2 mM L-glutamine (MT25005CI, Fisher/Corning), Penicillin (50 U/mL) and Streptomycin (50 μg/mL) (15140122, Fisher/Gibco), 1 mM sodium pyruvate (11360070, Fisher/Gibco), 1× MEM Non-Essential Amino Acids Solution (11140050, Fisher/Gibco). After the supplemented media was prepared, it was filter-sterilized through a 0.2 μm filter and then stored at 4°C for future use.

Cells were maintained at 30-80% confluency by regular passaging. First, media was aspirated and the cells were washed with 1x phosphate buffered saline (1x PBS; P38135, Millipore Sigma). Cells were then trypsinized (0.05%) (25-300-120, Gibco) for 5 minutes and finally the trypsin was quenched and cells were resuspended into a single-cell population with fresh media before plating on a new tissue culture dish. All cell lines were confirmed to be negative for mycoplasma infection.

Lipoprotein Depleted Serum (LDS) media was prepared by supplementing media with 5% LDS instead of 5 or 10% FBS (for IMDM or DMEM medias, respectively). To prepare LDS, 100% FBS was incubated with fumed silica (S5130, Millipore Sigma) (1 gram per 50 mL FBS) overnight at 4°C with constant shaking. After 24 hours, the suspension was centrifuged at 4000 x*g* to pellet silica complexes. The supernatant was carefully decanted without disturbing the pellet, followed by addition of fresh fumed silica at the original 1 g/50 mL ratio. After a second overnight incubation at 4°C with shaking, the suspension was centrifuged again and then the supernatant was filtered through 0.2 μm filter, aliquoted and snap-frozen in liquid nitrogen for future use.

### Drug treatments

Small molecules used in this study include: 25-hydroxycholesterol (25HC) (11097, Cayman Chemical), the LXR inhibitor GSK2033 (1 μM) (25443, Cayman Chemical), the NPC1 inhibitor U18666A (1 μM) (10009085, Cayman Chemical) the ACC1 inhibitor Firsocostat (also named GS-0976 and ND-630) (30 μM) (HY-16901, MedChemExpress), the ACC1 inhibitor CP-640186 (20 μM) (HY15259, MedChemExpress), the HMGCR inhibitor Lovastatin (10 μM) (10010338, Cayman Chemical), the ACAT1/2 inhibitor SZ58-035 (50 μM) (S9318, Millipore Sigma), the FASN inhibitor C75 (10 μM) (10005270, Cayman Chemical), the long-chain fatty acyl CoA synthetase (ACSL) inhibitor Triacsin-C (5 μM) (BMLEI218, Enzo), ACSL4 inhibitor AS-252424 (10 μM) (10009052,Cayman Chemical), human adipose triglyceride lipase (ATGL) inhibitor NG-497 (50 μM) (HY-148756, MedChemExpress), oleic acid (200 μM) (O1383, Millipore Sigma), docosahexaenoic acid (DHA; 200 μM) (D2534, Sigma), palmitic acid (200 μM) (P5585, Millipore Sigma), M◻CD (5 mM) (AAJ6684714, Thermo Scientific), the DGAT1 inhibitor T863 (20 μM) (SML0539, Sigma-Aldrich), the DGAT2 inhibitor PF-06424439 (10 μM) (PZ0233, Sigma-Aldrich) and myriocin (10 μM) (63150, Cayman Chemical). Drug treatments were carried out in various media compositions indicated in the Figure legends, for 16 hours at 37°C unless otherwise noted in the Figure legends.

### Generation of HAP1 mutant cell library for random insertion mutagenesis screen

To randomly mutate the genome we used a retrovirus bearing a Gene Trap (GT) cassette that introduces a splice acceptor site and a poly A tail, which terminates transcription regardless of whether it inserts into introns or exons^77^. In contrast to CRISPR knockout screens, GT mutagenesis can identify regulatory mechanisms that originate in non-coding regions like promoters, and it can identify hypomorphic and dominant negative alleles by truncating proteins.

To produce GT retrovirus, 15 million HEK 293FT cells were seeded into six T-175 flasks in 10% FBS DMEM media lacking antibiotics. When cells reached 80% confluency, they were transfected with two retroviral transfer plasmids containing the GT in two reading frames (3.3 μg pGT-mCherry and 3.3 μg pG-+1-mCherry), a plasmid with viral packaging genes (4 μg of pCMV-Gag-Pol), a plasmid with viral envelope genes (2.6 μg of pCMV-VSV-G), and a plasmid that enhances translation (1.7 μg of pAdVAntage). First, 45 μL of X-tremeGENE HP was added to 450 μL of OptiMEM for 5 minutes at room temperature, and then plasmids were added to the mixture, which was briefly vortexed and then incubated for 30 minutes at room temperature. The transfection reaction was added dropwise to the 293FT cells. After 16 hours, virus-containing media was harvested, filtered through a 0.45 μm low-protein binding filter and then concentrated by ultracentrifugation at 65,000 x*g* and 4°C for 1.5 hours in a swinging bucket rotor. Eight hours later, the virus was harvested and concentrated for a second time. After centrifugation, the supernatant was removed, leaving ∼ 1 mL of media to avoid disrupting the viral pellet. Tubes were then sealed with parafilm and incubated overnight at 4°C^78^.

To generate the HAP1 cell library WT HAP1 cells were pre-sorted for haploidy using a Hoechst stain. Then, 20 million haploid HAP1 cells were seeded into each of three T-175 flasks in antibiotic free 10% FBS IMDM. Immediately before infecting with GT-bearing retrovirus, the viral pellets were combined and resuspended in 76 mL of antibiotic free 10% FBS IMDM containing 4 μg/mL polybrene. One-third of this mixture was added to each of the three flasks containing WT HAP1 cells. Cells were expanded and maintained in multiple flasks, such that 50-100 million cells were kept in passage at any given time to ensure the genotypic richness of the library was maintained. Flow cytometry analysis measuring the GT-based mCherry fluorescence was used to assess transduction efficiency, which was determined to be >90%.

### Random insertion mutagenesis screens

For each screen, 20 million HAP1-GT cells were seeded in 7x T-175 flasks in 10% FBS IMDM media. To find genes that increase PM cholesterol after NPC1 loss of function, all 7 flasks were treated 8 hours after seeding with 1 μM U18666A. 20 hours later, 1 flask of cells was harvested by trypsinization, pelleted at 1000 x*g*, and then snap frozen as an unsorted control. The remaining flasks were prepared for Fluorescence Activated Cell Sorting (FACS). Plates were processed one at a time, one per hour. After washing with 1x PBS, cells were trypsinized and then spun at 1000 x*g* to isolate a cell pellet. Cells were then resuspended in 4 μM mNG-ALOD4 diluted into ice cold Probe Block Buffer (1x PBS, 1% BSA, filter sterilized). After 20 minutes of staining on ice, cells with the top 10% of mNG-ALOD4 fluorescence were isolated with FACS. Approximately 1-2 million cells were harvested per T-175 flask. After FACS, cells were spun down and then seeded into the same 15 cm plate. Cells were then expanded until 6x T-175 flasks could be reseeded. FACS was carried out 2 consecutive times to enrich the cells with high mNG-ALOD4 fluorescence. After the second sort, 30 million cells were pelleted and snap frozen in liquid nitrogen.

To identify genes that regulate cholesterol trafficking in response to 25HC, 20 million HAP1-GT cells were seeded in 7x T-175 flasks in 10% FBS IMDM, as described above. On the day of FACS sorting, one plate was isolated as the unsorted control. The remaining 6 plates were treated in a staggered manner, with 1 flask treated every hour with 4 μM 25HC. Each total treatment time was 6 hours. After 6 hours of treatment, cells were harvested by trypsinization, pelleted, and then isolated using the same protocol described above. For this screen, cells were sorted a total of three times, which may give rise to the increased FDR-corrected p-values observed for this screen compared to the U18666A screen (**Fig. 1g** and**Supplementary Fig. 2e**).

To prepare the GeneTrap insertion libraries, genomic DNA was first extracted with a QIAamp DNA Mini Kit (Qiagen Cat. # 51304). Next, 20 μg of DNA was digested with 100 units of MseI (NEB Cat. # R0525L), and a second 20 μg was digested with 100 units of SpeI-HF (Cat. # R3133L). Next, the DNA was purified with a QIAquick PCR Purification Kit (Qiagen Cat. # 28104) and then pooled; 14 ug of pooled DNA was then used to perform seven replicate Linear Amplification (LAM) PCRs initiated from a 5’ biotinylated primer specific to a unique sequence in the GeneTrap cassette. The resulting PCR products were purified on 8 μL of magnetic Dynabeads M280 (Thermo Fisher Scientific Cat. # 60101). Next, a 5’-phospho,3’-dideoxy DNA linker complementary to a Solexa adaptor sequence was appended to the 3’ end of each amplicon with a CircLigase II ssDNA ligase (Epicentre, Cat. # CL9025K). Finally, the LAM PCR products were captured by a second round of PCRs, and the PCR product was analyzed on a 2% agarose DNA gel, along with a control reaction prepared from unmutagenized cells– a smear between 100 and 500 bp was clearly visible for the GeneTrap-mutagenized DNA. Successful reactions were pooled and a 10 nM solution of the product was sequenced^79^.

To map the sequence reads, the sequences in the FASTQ data files were aligned to the human genome (hg18) using the Bowtie alignment software. For each screen, the sorted cell populations were compared to an unsorted control population. GeneTrap insertions in annotated genes were classified as being in the sense or antisense orientation, and as falling into introns or exons. Inactivating insertions were those that fell in exons (regardless of orientation), or those that fell into introns in the sense orientation; GeneTraps that were antisense within introns are considered not inactivating. Enrichment of GeneTrap insertions for a given gene was calculated by comparing the relative frequency that a gene carried an inactivating insertion compared to the frequency by which a gene carried *any* GeneTrap insertion in the control dataset^79^.

### Crispr Cas9 mediated knockout of genes

To knockout a gene of interest we designed two single guide RNA (sgRNA) sequences using the Benchling software to direct Cas9 binding and cleavage at two separate genomic sites that are 200-300 base pairs (bp) apart^80^. A ∼250 bp spacing was previously found to be an ideal distance- it results in a deletion of that segment of DNA rather than two independent repair steps involving nucleotide insertion/deletions (indels). We used the Ensembl Genome Browser to ensure that our sgRNAs targeted early exons that were present in all common splice isoforms for that gene. Often, one sgRNA was targeted to the middle of an exon and the second was targeted to the up or downstream intron- this arrangement was considered to be favorable because it would disrupt both the coding sequence of that exon and splicing. Forward and reverse strand guide RNA oligos were designed following previous protocols^80^, and then these oligos were annealed to generate duplexed DNA. “Guide 1” duplex DNA was cloned into a px458-GFP plasmid and “Guide 2” duplex DNA was cloned into a px458-mCherry plasmid. The sequence verified plasmids were next transfected into HAP1 cells and after 24-48 hours cells that were double-positive for mCherry and GFP were single-cell sorted into a 96 well plate using fluorescence activated cell sorting (FACS).

These single-cell clones were allowed to grow until they formed a visible colony of cells in the 96-well plate. Cells were then washed with 1x PBS, trypsinized with 20 μL of trypsin, and then quenched with 100 μL of supplemented media. Ten uL of the resuspended cells were immediately transferred into a 200 μL PCR tube, and the remainder of the cells were transferred to a 24-well plate for further growth. The single cell clones were then genotyped. First, cells were lysed with Quick Extract (NC9904870, Fisher Scientific) to obtain the DNA (40 μL of Quick Extract was added to the 10 μL of cells). Next, the DNA lesion was analyzed by Polymerase Chain Reaction (PCR) using a forward and reverse primer annealing to genomic DNA ∼100 bp up and downstream of the expected CRISPR edit. Finally, PCR products were analyzed with DNA gel electrophoresis.

### Western blotting

Cells were plated on a 6 cm dish and treated with small molecules when indicated. In **Fig. 7d**, primary hepatocytes were isolated from 8-week old mice and then plated on collagen-coated coverslips prior to analysis. Cells (including HAP1 cells, MEFs, and hepatocytes) were then washed and scraped with chilled 1x PBS, and then pelleted by spinning at 1000 x*g* for 5 minutes at 4°C. After removing the supernatant, the cells were lysed with Lysis Buffer (10% glycerol, 2x Protease Inhibitor Cocktail (S8830, Millipore sigma), 150 mM NaCl, 50 mM Tris-HCl (pH 8), 1% NP-40 and 1 mM MgCl_2_) at 4°C using a mechanical shaker for 1 hour. Lysates were spun for 30 minutes at 18,000 x*g* and 4°C, and then supernatants were transferred to a new tube. A BCA assay was performed on the supernatant to determine the total protein concentration, and then samples containing equal amounts of protein (∼40 μg per well) were incubated with 1X mPAGE LDS Sample Buffer (MPSB, Millipore Sigma) and Dithiothreitol (25 mM final concentration) (BP17225, Fisher) at 37 °C for 30 minutes before performing SDS-PAGE.

After completion of the SDS-PAGE, samples were transferred from the gel to a PVDF membrane (IPVH85R, millipore sigma). Membranes were blocked with 5% nonfat milk in TBST (1x Tris-Buffered Saline, 0.1% Tween-20). For analysis of knockout cell lines, the membranes were incubated with the appropriate dilutions of commercially available primary antibodies (ACAT1, ACC1, NPC1, FASN, ATGL) in 5% nonfat milk diluted in TBST for 2 hours at room temperature, washed 3 times with TBST for 5 minutes each, and then incubated with HRP-conjugated secondary antibodies diluted to 1:10,000 for 1 hour at room temperature. After washing the membrane three times with TBST for 5 minutes each the blot was developed using ECL solutions (PI32209, Fisher/ Pierce).

### Flow cytometry analysis of plasma membrane accessible cholesterol or sphingomyelin-cholesterol complexes

Purification of mNG-ALOD4 and OlyA-EGFP were carried out by transforming ALOD4 and OlyA plasmids into Rosetta TM2 2(DE3) pLysS E. coli competent cells (Millipore Sigma, Cat. # 71403-3). Next, a 1 L liquid culture was grown at 37 °C until an OD_600_ of ∼0.5, at which point the temperature was reduced to 18 °C before protein expression was induced with 1 mM IPTG for 24 hours. Bacteria were harvested by centrifugation and then lysed with 50 mM Tris-HCl pH7.4, 150 mM NaCl, and 1 mM DTT with protease inhibitors. After removing aggregated material by centrifugation, mNG-ALOD4 or OlyA-EGFP were first purified with complete His-Tag Purification resin (Millipore Sigma, Cat.# 5893682001) before further purification with size exclusion chromatography^81^. To measure cell surface lipids, the cells of interest were seeded in a 24-well cell culture plate (3 replicate wells for each condition) aiming for ∼75% confluency on the day of flow cytometry analysis. To prepare cells for flow cytometry, they were first rinsed with 1x PBS, harvested with trypsin, and quenched with 4°C 5% FBS containing IMDM media (or 10% FBS containing DMEM for 3T3, 293T or MEF cells) while gently pipetting up and down to generate a single cell suspension. Cells were then transferred to a 1.5 mL eppendorf tube on ice, and then spun for 5 minutes at 4°C (HAP1 cells at 1000 x*g* and 3T3/293T cells at 750 x*g*). The supernatant was removed and cell pellets were resuspended in 25 μL of 2 μM mNeonGreen-ALOD4 (mNG-ALOD4) or 4 μM OlyA-EGFP diluted in ice cold 1x Probe Block Buffer (PBB- 1x PBS, 1% BSA, filter sterilized through a 0.2 μm filter). The cells were incubated on ice, 12 minutes for mNG-ALOD4 or 30 minutes for OlyA-EGFP, before being diluted to 150 μL in PBB, and then transferred to a 96-well plate.

Flow cytometry was performed using the BD Accuri C6 Plus Flow Cytometer or the Sony Cell Sorter. During flow cytometry, cells were first gated based on forward scatter (FSC) area versus height, such that only single-cells and living cells were analyzed. Secondly, cells were gated based on FSC area versus side scatter (SSC) area to gate for cells with similar granularity and remove cells that were either sick or stressed. The final gated population made up at least 60% of the total cells. The fluorescence of mNG-ALOD4 or OlyA-EFGP was measured by exciting using a 488 nm laser and detecting the emission through a 585/40 nm bandpass filter. For each measurement, at least 5000 cells were analyzed. The FCS files generated from BD CSampler software or the Sony Cell Sorter Software were loaded to FCS Express software and then exported directly to GraphPad Prism for analysis.

To assess how various small molecules impacted PM accessible cholesterol in kinetic assays (e.g. 25HC-mediated PM accessible cholesterol depletion) small molecules were added at the indicated times prior to flow cytometry analysis both with and without 16 hour prior incubation with the noted inhibitors. Every well was treated independently– each time point represents three independent wells that are not experimentally coupled to the wells for other time points.

### Measuring plasma membrane accessible cholesterol using microscopy

Cells were seeded on acid-washed coverslips and grown to ∼60% confluency. The ACC1 inhibitor Firsocostat (30 μM) or M◻CD:cholesterol (0.3 mM) were added 16 hours before analysis. MβCD:cholesterol complexes were prepared by dissolving cholesterol in chloroform:methanol (2:1, v/v) to generate a 10 mg/mL (25.86 mM) stock solution. An aliquot containing 8.7 μmol cholesterol was transferred to a glass vial and the solvent was evaporated under a stream of nitrogen to form a thin sterol film. MβCD was dissolved in Opti-MEM at 50 mg/mL (38 mM) and 2 mL was added to the dried sterol film. The mixture was sonicated in a room-temperature water bath for 2–5 min followed by micro-tip sonication on ice until the solution became clear. The resulting solution was passed through a 0.2-μm filter and stored at 4 °C or aliquoted and stored at −80 °C^81^. The coverslips containing cells were moved to a metal rack on ice (with cells facing up), and then washed once with ice cold 1x PBS. Cells were then stained with 2 μM mNG-ALOD4 diluted in ice cold 1x PBB for 30 minutes. Cells were then washed three times with ice cold 1x PBS and then fixed for 10 minutes on ice with 4% paraformaldehyde diluted in 1xPBS. Finally, cells were washed an additional three times with ice-cold 1x PBS before being mounted on glass slides using ProLong Diamond Antifade Mountant containing DAPI to visualize the nuclei (P36962, Invitrogen/Thermo Fisher) and left to cure overnight at room temperature before imaging. Images were acquired on an inverted Leica SP8 laser scanning confocal microscope with a 63× oil immersion objective lens (numerical aperture 1.4).

### Measuring sphingomyelin levels with sphingomyelinase

HAP1 WT cells and HAP1 *ACC1*-KO cells were seeded in 5% FBS IMDM in a 24-well plate and grown to ∼50% confluency. Cells were then left untreated or treated for 16 hours with 30 μM Firsocostat in 5% FBS IMDM. The day of the assay, cells were first washed with 1x PBS and then incubated at 37 °C with sphingomyelinase (SMase) (150 mU/mg protein,S8633, Millipore Sigma) for 30 minutes to hydrolyze sphingomyelin. After incubation, the cells were washed and trypsinized for flow cytometry analysis as described above^11^.

### Measurement of free cholesterol and cholesterol esters

Cells were plated in a 12-well tissue culture dish in 5% FBS IMDM media and grown to 80% confluency. On the day of the assay, cells were washed with 1x PBS and lysed for 10 minutes at 4°C with 150 μL of lysis buffer containing 10% glycerol, 2x Protease Inhibitor Cocktail (S8830, Millipore sigma), 150 mM NaCl, 50 mM tris-HCl (pH 8), 1% NP-40 and 1 mM MgCl_2_. The Amplex Red Cholesterol Assay Kit (A12216, Invitrogen/Fischer) was used to measure the free cholesterol and cholesterol esters in each sample. After lysis, lysates were diluted with an equal volume of 1x reaction buffer from the kit. Total cholesterol and free cholesterol were measured by preparing two working solutions (with and without cholesterol esterase) as described in the assay protocol and mixing the lysates with the mastermix in a 1:1 ratio in a 96-well plate. The plate was incubated at 37 °C for 30 minutes and the fluorescence emission was measured at 590 nm using a microplate reader.

### Microscopy analysis of lipid droplets (LDs)

For LD analysis, the cells of interest were seeded on acid-washed coverslips in a 24-well plate. Various treatments were performed as described in the figure legends. To stain for LDs, cells were washed with 1x PBS and fixed in a solution containing 4% paraformaldehyde (PFA) and 1x PBS for 10 minutes at room temperature. After fixing, cells were washed 3 times with 1x PBS, and then cells were permeabilized with immunofluorescence (IF) block buffer (1x PBS, 1% BSA and 0.1% Triton X-100, filter sterilized through a 0.2 μm filter) for 15 minutes at room temperature. Cells were then incubated with 10 μM BODIPY 493/503 (Cayman Chemical; Cat# 25892) in IF block buffer for 1 hour followed by 3 washes with 1x PBS. Coverslips were mounted on glass slides using ProLong Diamond Antifade Mountant containing DAPI to visualize the nuclei (P36962, Invitrogen/Thermo Fisher) and left to cure overnight at room temperature before imaging. Imaging was conducted using a Keyence BZ-X800L fluorescence microscope.

To co-stain for ACC1 and LDs, cells were treated the same as above. After the 15 minute permeabilization step, cells were stained with ACC1 primary antibody (RRID: AB_2882621) diluted 1:1000 in IF block buffer for 1 hour at room temperature. Coverslips were then washed 3x with IF block buffer before adding a mixture of 10 μM BODIPY 493/503 and secondary antibody (anti- mouse 647; RRID:AB_3073503) diluted 1:500 in IF block buffer for 1 hour. Alternatively (**Supplementary Fig. 6a**), CoraLite Plus 647-conjugated ACC1 mouse polyclonal antibody (RRID: AB_11042445) was diluted 1:500 along with 10 μM BODIPY 493/503 in IF block buffer. After 1 hour of incubation, cells were washed 3x with IF block buffer, and then mounted on glass slides and imaged as described above.

### Quantification of LDs

To quantify LDs, images were processed using ImageJ software. First, images were imported into ImageJ and then the images were converted to an 8-bit grayscale format. Thresholding was performed to eliminate background by manually adjusting the threshold intensity. Within a single experiment, all threshold values were kept constant for equal comparisons of the images.

LDs were then quantified using the analyze particles function in ImageJ and the parameters were configured to limit particle size to a minimum of 2 pixels² to eliminate very small thresholded noise; circularity was set to 0.00-1.00. The results window provided raw data, including the LD area (pixels² was converted into μm² using the conversion factor for our microscope = 0.12581 μm/ pixel), which was copied into excel for analysis. LDs per cell were calculated by dividing the total number of LDs for a field from the total number of nuclei in a given field. In **Supplementary Fig. 6b**, LDs were counted manually and assigned to a specific cell to get a measurement of per-cell LD numbers. This latter method was not a preferred method because when cells reach >70% confluency, it is often difficult to ascertain which LD belongs to which cell.

### Cellular fractionation to isolate LDs

HAP1 WT cells were seeded on two 15 cm plates, with one plate treated with 30 μM Firsocostat for 16 hours. Cells were harvested by gentle scraping in ice cold 1x PBS. The cell suspensions were then centrifuged at 1000 x*g* for 5 minutes at 4°C to obtain cell pellets. After removing the supernatant the pellets were resuspended in 8% sucrose buffer containing 0.25 M sucrose, 50 mM Tris-HCl (pH 7.4), 150 mM NaCl, 1x Protease Inhibitor Cocktail (S8830, Millipore sigma), and 1x Phosphatase Inhibitor (4906845001,Millipore Sigma). After incubating for 10 minutes on ice, the cells were lysed by passage through a chilled ball-bearing cell homogenizer (Isobiotec) with a 12 μm clearance 8 times. Successful lysis was confirmed using Trypan blue staining (15250061, Gibco). A fraction (∼50 μL) of the homogenate was saved for Western blotting (whole-cell lysate). The remainder of the homogenate was centrifuged twice at 1000 x*g* for 10 minutes each to pellet nuclei. The supernatant (∼600 μL) was pipetted into a separate chilled microtube and the pelleted nuclei was saved for Western blotting.

To isolate LDs, 600 μL of 0% sucrose buffer (150 mM NaCl and 50 mM Tris-HCl pH 7.4) was slowly added on top of the supernatant isolated above while trying to maintain the separation of different sucrose densities. The tubes were centrifuged at 130,000 x*g* for 1 hour at 4°C. LDs were collected in four, 50 μL fractions (total volume 200 μL) using a pre-wetted pipette tip starting from the top floating layer, which appeared cloudy. The remaining ∼400 μL of 0% sucrose buffer was removed. The bottom ∼600 uL cytoplasmic fraction was collected and saved for Western blotting. Finally, the pelleted membrane fraction was resuspended in Lysis Buffer (10% glycerol, 2x Protease Inhibitor Cocktail (S8830, Millipore sigma), 150 mM NaCl, 50 mM tris-HCl (pH 8), 1% NP-40 and 1 mM MgCl_2_) and saved for Western blotting.

The four LD fractions were washed with 500 μL 0% sucrose buffer by spinning at 20,000 x*g* for 30 minutes at 4°C. Next using a syringe needle, ∼600 μL from the bottom of the tube was removed and the top portion was saved for Western blotting of the LD fraction.

For Western blot analysis, the whole cell fraction was lysed by adding NP-40 to 1%. The nuclear fraction was also resuspended in Lysis Buffer. The whole cell, nuclear, and membrane fractions were mechanically shaken at 4°C for 45 minutes to aid in lysis. The lysates were then clarified by spinning at 18,000 x*g* for 30 minutes to remove aggregated material and DNA. Finally, protein concentrations for the whole cell, nucleus, cytoplasm and membrane fractions were measured with a BCA assay, and equal amounts of protein were saved for Western blots. LD fractions were also lysed by adding 1% NP-40, but BCA assays revealed very low protein concentrations. For Western blot, the ∼40 μL (a maximum possible volume) of lysed LDs were loaded.

### Mass spectrometry analysis of lipid species

WT HAP1 cells and TAG-TKO HAP1 cells were grown on four 15-cm dishes per treatment condition until ∼60% confluent in 5% FBS IMDM. 48 hours before analysis, all cells were moved into 5% LDS media. Cells receiving the ATGL inhibitor NG-497 (50 μM) were treated at this time for 48 hours. The ACC1 inhibitor Firsocostat (30 μM) was a 24-hour treatment (added after 24-hour preincubation in 5% LDS). For cells that received both NG-497 and Firsocostat, NG-497 was added first for 24 hours, and then both inhibitors were added together for a second 24 hours. To isolate cells, media was aspirated and the cells were washed gently with room temperature 1x PBS. Cells were then scraped in ice cold 1x PBS on an ice-cold metal rack in a tissue culture hood. Each plate of cells was moved to a single 15 mL conical tube and then spun at 1000 x*g* to pellet. The PBS supernatant was aspirated, and then cells were snap frozen in liquid nitrogen and stored at -80°C.

To extract lipids, cell pellets were resuspended in 50 μL of 1x PBS (enough to pipette up and down a homogenous mixture- the total volume was ∼150 μL), and then 100 μL of the concentrated cells were added to a sterile 2 mL Eppendorf tube. Next, 250 μL of LC/MS grade ice-cold methanol (A456-500, Fisher Scientific) was added to the cell pellet along with 10 μL of EquiSPLASH™ LIPIDOMIX® (330731, Avanti Polar Lipids). EquiSPLASH™ was also added to a tube that was later used as an internal standard blank. After vortexing to aid in cell-lysis, 750 μL of methyl tert-butyl ether (MTBE; AA40477K2, Fisher Scientific) was added to the sample; this mixture was again vortexed and then chilled for 30 minutes. Finally, 250 μL of ice-cold LC/MS-grade water (AA47146Y0, Fisher Scientific) was added to induce phase separation. Samples were then centrifuged at 15,000 *xg* and 4°C for 15 minutes. The upper organic phase was carefully transferred to a new tube and then dried under nitrogen gas. Dried lipids were reconstituted in 100 μL of Mobile Phase A (93:7 acetonitrile:dichloromethane; 047138M1 and D151SK-4, Fisher Scientific) for analysis. Quality control, extraction blanks, and internal standard-only controls were included throughout. For storage, lipid samples were dried down under nitrogen and reconstituted with methanol with 1% butylated hydroxytoluene (ICN10116290, Fisher Scientific) and stored at -80°C to prevent oxidation.

Targeted lipidomics was conducted using a Sciex 7500+ Triple Quadrupole mass spectrometer coupled with the Exion UHPLC system. Samples were run on a phenomenex Amide column (Luna 3 μm NH2 100 Å, LC column 100 x 2 mm (00D-4377-B0, Phenomenex). Mobile Phase A was 93:7 acetonitrile:dichloromethane and Mobile Phase B was 1:1 methanol:water with 1 mM ammonium acetate (pH 8.2) added. Sample injections were 5 μL, which were analyzed in both positive and negative ion modes using HILIC chromatography^82^. 2,200 lipid transitions were monitored over a 20 minute method^83^. Data analysis was completed in Sciex OS version 3.5.; chromatographic peaks were integrated using the MQ4 integration algorithm with a S/N cutoff of 4 and a Gaussian smooth of 1– additionally, peaks were manually reviewed to ensure they had the correct boundaries. Lipid species were annotated based on established precursor/product-ion transitions and retention times. Each lipid class was normalized to the EquiSPLASH internal standards. For lipid classes without an internal standard, chemically similar lipid standard was used. Statistical analyses were performed in excel with a two-tailed two-sample unequal variance (Welch’s) t-test.

To analyze DAGs and TAGs, samples were run on an ACQUITY UPLC BEH C18 Column (130 Å, 1.7 μm, 2.1 x 100 mm) (176000864, Waters). Mobile Phase A was 1:1 methanol:water with 1 mM ammonium acetate (pH 8.2) added and Mobile Phase B was 95:5 methanol:water with 1 mM ammonium acetate (pH8.2) added. A 14-minute method was used and 220 multiple reaction monitoring (MRM) transitions were monitored in negative mode to detect different species of TAGs and DAGs. EquiSPLASH internal standards were used for normalization, and analysis was performed as described above. The EquiSPLASH TAG internal standard was used for normalization of DAGs due to DAG internal standard degradation. This work was supported by the Vincent Coates Foundation Mass Spectrometry Laboratory, Stanford University Mass Spectrometry (RRID:SCR_017801) utilizing the Sciex 7500+ (RRID:SCR_026608).

For **Supplementary Fig. 3**, WT HAP1 cells were grown on four 15-cm dishes per treatment condition until ∼70% confluent in 5% FBS IMDM. Cells were then moved to 5% LDS media with or without 30 μM Firsocostat for 24 hours. After isolating cells via the same protocol discussed above, cell pellets were shipped on dry ice to Metabolon Inc. for analysis. Raw values obtained from Metabolon Inc. are areas under the curve for each detected lipid species, normalized to protein concentrations for each sample. Values are not exact quantitative measurements since internal standards were not used.

For analysis of lipids from *Acc1/Acc2* flox/flox mice and Albumin-CRE *Acc1/Acc2* flox/flox mice, mice were chow-fed, with n=4 per group. Lipids were extracted by liquid-liquid extraction and analyzed by LC-MS/MS using internal standards. Each value was normalized to tissue weight.

### Measurement of membrane fluidity with Laurdan

For flow cytometry measurements (Fig. 6j), cells were grown in triplicate for each condition in a 24-well dish in 5% FBS serum until ∼50% confluent. All cells were then moved to 5% LDS-containing media for 48 hours prior to analysis, and treatments were carried out in this media. The ATGL inhibitor (50 μM NG-497) was added for 48 hours; the ACC1 inhibitor (30 μM Firsocostat) and docosahexaenoic acid (DHA; 150 μM) were added for 24 hours. M◻CD (5 mM) treatments were carried out for 20 minutes immediately before analysis. Cells were harvested by washing briefly with 1x PBS, trypsinizing for 5 minutes, and then quenching the trypsin with room temperature 5% FBS DMEM. Cells were then spun at 1000 *xg* to pellet. Pelleted cells were resuspended in 5 μM Laurdan diluted into 1x PBS for 30 minutes at room temperature. Finally, fluorescence was measured at the Stanford Shared FACS Facility (RRID: SCR_017788) on a NovoCyte Penteon flow cytometer using laser configurations for ZombieAqua and DyeCycle Violet. Median fluorescence measurements were used to calculate the Laurdan GP value [GP = (Median_440nm_ - Median_490nm_)/(Median_440nm_ + Median_490nm_)].

Measurements of PM fluidity were carried out by plating cells on 8-well Ibidi live-cell imaging chambers (NC1930679, Fisher Scientific). All treatments were carried out exactly to the flow cytometry treatments. To image live cells, media was removed and replaced with warmed 5% LDS media containing 5 μM Laurdan. After 30 minutes, live cells were imaged on a Leica SP8 confocal at the Stanford University Cell Sciences Imaging Core Facility (RRID: SCR_017787). This microscope was supported, in part, by Award Number 1S10OD010580 from the National Center for Research Resources (NCRR). Its contents are solely the responsibility of the authors and do not necessarily represent the official views of the NCRR or the National Institutes of Health. Laurdan was excited with a 405nm laser, and emissions were collected in spectral scanning mode at a single focal plane for a given field of cells. To generate the images in **Fig. 6k**, the fluorescence signal from the denoted emission wavelength was inverted in ImageJ. To calculate the GP values (**Fig. 6l**), a segment of a single cell plasma membrane (∼½ the length of one edge of the cell) was manually circled. The fluorescence signal was then measured (by pressing the “m” key while in the ImageJ software) for both the 440nm and 490nm wavelengths. The average fluorescence intensities were then moved into an excel file, and the GP value was calculated as described above. Each circle in the plot represents the GP value for one segment of PM, and three-four independent fields of cells were analyzed to generate the final plot.

### Analysis of cholesterol extractability with M◻CD dose curves

All cells were grown in triplicate for each condition in 24-well plates and 5% FBS-containing media until ∼60% confluent. Media was then replaced with 5% LDS-containing media for 24 hours with or without small molecule treatments prior to analysis. ACC1 inhibitor (50 μM Firsocostat) and oleic acid (150 μM) were added for 24 hours. Myriocin (30 μM) was added for 72 hours in total- the first 48 hours were in 5% FBS and the final 24-hours were in 5% LDS. Immediately prior to analysis, media was removed and cells were treated with different doses of M◻CD (indicated in the figure) diluted into 5% LDS for 20 minutes at 37°C. MBCD was then removed and cells were washed with 1xPBS before trypsination. All downstream analysis steps were identical for that described in the section on “Flow cytometry analysis of plasma membrane accessible cholesterol or sphingomyelin-cholesterol complexes.”

### Measurements of the rate of fatty acid and cholesterol synthesis in male mouse livers

All male mice were used throughout this study. Male mice are more susceptible to developing fatty liver disease and they also do so more rapidly, allowing for reduced timescales for experiments. Secondly, and importantly, male mice also develop fatty liver disease more uniformly (at similar times and extents) compared to female mice, which may be partially protected by estrogen and are more variable due to their estrous cycle^84^. *Acc1* (*Acaca*) floxed mice were generated with loxP sites flanking exons 22–26, and *Acc2* (*Acacb*) floxed mice were obtained from The Jackson Laboratory. Hepatocyte-specific *Acc1/Acc2* double-knockout mice were generated by crossing *Acaca*^f/f^;*Acacb*^f/f^ mice with albumin-Cre transgenic mice^60^. For analysis of fatty acid and sterol synthesis rates, male mice were maintained on a chow diet and subjected to time-restricted feeding (feeding from 6 PM to 6 AM and fasting from 6 AM to 6 PM) for 3 consecutive days. Mice were injected intraperitoneally with [³H]H₂O (50 mCi), and samples were collected 1 hour later. Plasma [³H]H₂O specific activity was determined, and 200–300 mg of liver was saponified. Sterols were isolated by digitonin precipitation, whereas fatty acids were recovered by acidification and two petroleum ether extractions. Tracer incorporation into each fraction was normalized to plasma specific activity and liver mass^85^. Hepatic sterol synthesis rates were assessed by measuring [³H] incorporation into newly synthesized sterols^85^.

### Measurement of total liver cholesterol and fecal cholesterol content in male mice

All male mice were used throughout this study. For hepatic liver cholesterol measurements, approximately 100 mg of liver tissue was used for lipid extraction, and cholesterol quantification was measured using the Infinity™ Cholesterol Liquid Stable Reagent.

For fecal cholesterol measurements, feces were collected from mice (n = 7 per group), which were individually housed overnight. Around 100 mg of feces was homogenized in Folch reagent (chloroform:methanol, 2:1), and the organic phase was extracted by phase separation with saline and isolated by centrifugation. A portion of the organic phase was dried in the presence of Triton X-100, and cholesterol content was measured using the Infinity™ Cholesterol Liquid Stable Reagent.

### RNA sequencing analysis

WT HAP1 cells were grown in 6-well plates, with 4 wells per condition, until ∼75% confluent in 5% FBS IMDM. Cells were then moved into 5% LDS media for 16 hours. Cells were then treated with or without 10 μM Lovastatin, 30 μM Firsocostat, 5 μM Triacsin-C or 10 μM C75 for 6 hours. Cells were then washed twice with 1x PBS, and then all PBS was aspirated.

RNA was isolated using Trizol-chloroform extraction. Briefly, 800 μL Trizol was added directly to the wells, incubated for 5 minutes, and then transferred to DNase/RNase-free 1.5 mL eppendorf tubes. After adding 160 μL chloroform, samples were shaken vigorously for 15 seconds and incubated at room temperature for 3 minutes. Phase separation was achieved by centrifugation at 12,000 x*g* for 15 minutes at 4 °C, and then the aqueous phase was transferred to a new tube. Next, 400 μL isopropanol was added and then samples were incubated for 10 minutes at room temperature. RNA was then pelleted by centrifugation at 12,000 x*g* for 10 minutes at 4 °C. The RNA pellet was washed with 800 μL 75% ethanol, vortexed briefly, and centrifuged at 7,500 x*g* for 5 minutes at 4 °C. The supernatant was removed, and pellets were spun again for 5 minutes. Residual ethanol was removed and pellets were air-dried at room temperature for 5–10 minutes until the pellet turned translucent. RNA was resuspended in 40 μL DNase/RNase-free water, incubated at room temperature for 10 minutes, and gently mixed. RNA was stored at -80 °C and shipped to Novogene USA Inc. for standard RNA sequencing analysis. The returned FASTQ files were analyzed using the PartekFlow software using methods described by Xiaowen Wang (Field Application Specialist from Partek Incorporated). Data was processed using a differential gene set enrichment analysis (DeSeq2).

The bubble plots (**Fig. 7h**, **i**) were generated using both GraphPad Prism 10 and Adobe Illustrator. First, a list of genes of interest were curated for SREBP2 (**Fig. 7h**) and SREBP1 (**Fig. 7i**). SREBP1 targets were found using a publically available CHIP-Seq database. Next, FDR-corrected *p*-values were -Log_10_ transformed, and Log_2_ fold changes values were subjected to exponential transformation (2^(Log_2_(fold-change values)). The transformed *p*-values and fold change values for genes of interest were found for each treatment and given a unique identifier (e.g. HMGCRi-ACAT2, ACC1i-ACAT2, ACSLi-ACAT2, and FASNi-ACAT2). All genes were then plotted into one column using a Multiple Variable analysis plot type in Graph Pad Prism 10. Bubble size was set to be determined by the transformed *p*-value, and bubble color was set to be the fold change; this software also assigned x-axis and y-axis positioning for the bubbles, however these were not used in the final figure generation. GraphPad Prism 10 was simply used to generate bubbles of meaningful size and color for each gene in its respective treatment. Finally, the GraphPad Prism 10 file was exported as a .emf file, opened in Adobe Illustrator, and the bubbles were re-arranged into a column graph.

### Gene Set Enrichment Analysis

To analyze the top pathways affected by ACC1, ACSL, FASN, and HMGCR inhibition, we utilized the Gene Set Enrichment Analysis (GSEA) website. First, we navigated to the Explore the Molecular Signatures Database (MSigDB), and then to the “Investigate” tab in the Human Collections. In the “Input Gene Identifiers” box, we pasted the top 200 genes that were differentially regulated for each treatment– rankings were made by sorting based on FDR-corrected *p*-values. In the Compute Overlaps window, we selected the WikiPathways gene sets, and then computed the overlaps. This returned the top 10 WikiPathways for each set of genes. Enterocyte Cholesterol Metabolism (WP5333), Cholesterol Metabolism (WP5304), Cholesterol Biosynthesis Pathway (WP197), and Cholesterol Biosynthesis Pathway in Hepatocytes (WP5329) were excluded from the final plot due to their significant overlap with the genes found in Cholesterol Metabolism with Bloch and Kandutsch-Russell Pathways (WP4718) and Cholesterol Synthesis Disorders (WP5193), which were used in the final plot. If a pathway was not found in the top 10 pathways identified, it was denoted in the plot as “nf” for not found.

### Statistical analysis and schematic generation

Schematics (**Fig. 1e**, **1f**, **2f**, **3a**, **3k**, **4a**, **5a**, **6m**, **6n**, **7a**, **7b**, and **7d**; also **Supplementary Figure 1a**, **1h** (top), and **3a**) were generated in Adobe Illustrator. All other plots were generated in GraphPad Prism 10.6.1 and then moved to Adobe Illustrator for further labeling and to compile each figure. **Fig. 3k** shows the Alphafold 3 structure of ATGL– this was opened in ChimeraX where it was colored pink and then truncated to only show the globular domain. Finally, it was exported as a .png file for further editing in Adobe Illustrator.

Data analysis was completed in GraphPad Prism 10.6.1. To determine which statistical test to use we used the following decision tree: (**1**) are only two datasets being compared or (**2**) are more than two datasets compared? If two datasets were compared (**1**), we used a Welch’s t-test; this is the recommended approach to compare two independent groups and it is robust when the variance is unequal. If more than two datasets were compared, we assessed if a single variable was compared across multiple groups (**2**- e.g. one cell line with different treatments, comparing untreated to each treatment) or if multiple independent variables were compared (**3**- e.g., treatment and genotype or treatment and time). When a single variable was compared across multiple groups (**2**), a one-way ANOVA was used. For all one-way ANOVAs used, we ensured an equal spread of variance with a Brown-Forsythe test. When making an all-to-one comparison (**2a**) (e.g. all treatments to untreated), we used a Dunnett’s test post-hoc, otherwise when making comparisons without a clear comparison structure (**2b**), we used a Sidak correction post-hoc. Post-hoc Tukey tests (**2c**) are ideal when every possible comparison is made, which was not used in this manuscript. One experiment had unequal variance spread (**Fig. 2i**); for this experiment, we performed a Welch’s ANOVA with a Dunnett T3. Next, if multiple independent categorical factors were compared (**3**), we performed a two-way ANOVA with a Sidak correction ,which reduces false positives while maintaining statistical power. *P*-values from GraphPad Prism 10.6.1 are presented on the graphs using the following notation: not significant (ns) *p*>0.0332, \**p*≤0.0332, \*\**p*≤0.0021, \*\*\**p*≤0.0002, and \*\*\*\**p*<0.0001. Unless stated otherwise, all experiments were repeated three times. Exact *p*-values and information about statistical tests for the Supplementary data are listed in the Supplementary data figure legends. Main figure information is shown here:

**Figure 1** exact *p*-values are: (**a**) WT vs *GRAMD1*-TKO *p*<0.0001; (**b**) WT vs *GRAMD1*-TKO: 2 hr 25HC *p*=0.9916, 4 hr 25HC *p*=0.9879, and 6 hr 25HC *p*=0.9533; (**c**) control vs LXR inhibitor *p=*0.0468; (**d**) control vs LXR inhibitor: 2 hr 25HC *p*=0.1852, 4 hr 25HC *p*=0.0415, and 6 hr 25HC *p*=0.0011; (**h**) WT vs *ACC1*-KO: 3 hr 25HC *p*<0.0001, 4.5 hr 25HC *p*<0.0001, and 6 hr 25HC *p*<0.0001; and (**i**) untreated vs ACC1i: 3 hr 25HC *p*<0.0001, 4.5 hr 25HC *p*<0.0001, and 6 hr 25HC *p*<0.0001.

**Figure 2** exact *p*-values are: (**b**) WT vs *ACC1*-KO *p*<0.0001; (**c**) WT HAP1 untreated vs ACC1i *p*<0.0001, 293T untreated vs ACC1i *p*<0.0001 and 3T3 untreated vs ACC1i *p*<0.0001; (**d**) untreated vs CP-640186 *p*<0.0001 and untreated vs Firsocostat *p*<0.0001; (**e**) WT vs *ACC1*-KO: 0.2 μg/mL PFO* *p*=0.8607, 0.5 μg/mL PFO* *p*=0.0380, 1.3 μg/mL PFO* *p*<0.0001, 3.2 μg/mL PFO* *p*<0.0001, 8 μg/mL PFO* *p*=0.8388, 20 μg/mL PFO* *p*>0.9999, and Triton X-100 *p*>0.9999; (**f**) control vs control + SMase *p*<0.0001, ACC1i vs ACC1i + SMase *p*<0.0001, ACC1-KO vs ACC1-KO + SMase *p*<0.0001, and ACC1i alone vs control + SMase p= 0.1932. (**h**) WT untreated vs 25HC *p*=0.0003 and *ACAT1/2*-DKO untreated vs 25HC *p*=0.8854; (**i**) WT vs *ACAT1/2*-DKO untreated *p*=0.3811, *ACAT1/2*-DKO vs *ACAT1/2*-DKO + ACC1i *p*<0.0001, and WT vs *ACAT1/2*-DKO + ACC1i *p*<0.0001; (**j**) *ACAT1/2*-DKO vs *ACC1/ACAT1/2-*TKO: 3 hr 25HC *p*<0.0001, 4.5 hr 25HC *p*<0.0001, and 6 hr 25HC *p*<0.0001.

**Figure 3** exact *p*-values are: (**b**) ACC1i 0 hr vs 3hr *p*=0.0004, ACC1i 0 hr vs 6hr *p*<0.0001, ACC1i 0 hr vs 24hr *p*<0.0001, and ACC1i 0 hr vs 48hr *p*<0.0001; (**c**) control vs ACSLi *p*<0.0001, WT vs *ACSL1/3*-DKO *p*<0.0001, and *ACSL1/3*-DKO vs *ACSL1/3*-DKO + ACSL4i *p*=0.0001; (**d**) control vs ACSLi: 2 hr 25HC *p*<0.0001, 4 hr 25HC *p*<0.0001, and 6 hr 25HC *p*<0.0001; (**e**) WT vs *ATGL*-KO: 2 hr ACC1i *p*=0.4510, 4 hr ACC1i *p*= 0.0002 and 6 hr ACC1i *p*<0.0001; (**f**) control vs ATGLi: 2 hr ACC1i *p*=0.1119, 4 hr ACC1i *p*<0.0001, and 6 hr ACC1i *p*<0.0001; (**g**) untreated vs ACC1i *p*<0.0001, untreated vs ATGLi *p*<0.0001, untreated vs ACC1i + ATGLi *p*=0.5874, and ACC1i vs ACC1i + ATGLi *p*<0.0001; (**h**) untreated vs OA *p*<0.0001, ACC1i vs ACC1i + OA *p*<0.0001 and ACSLi vs ACSLi + OA *p*=0.8073; (**i**) WT ACC1i vs ACC1i + OA *p*<0.0001, and *ATGL*-KO ACC1i vs ACC1i+OA *p*=0.9939; and (**j**) ATGLi 0 hr vs 2 hr *p*<0.0001, ATGLi 0 hr vs 4 hr *p*<0.0001, ATGLi 0 hr vs 6 hr *p*<0.0001, and ATGLi 0 hr vs 24 hr *p*<0.0001.

**Figure 4** exact *p*-values are: (**d**) LD numbers (purple curve) *p*<0.0001 across all comparisons, PM changes (blue curve) *p*<0.0001 across all comparisons; (**e**) LD numbers (purple curve) *p*=0.7987 across all comparisons, PM changes (blue curve) *p*=0.0017; (**f**) LD numbers (purple curve) *p*<0.0001 across all comparisons, PM changes (blue curve) *p*<0.0001 across all comparisons; (**g**) WT untreated vs ACC1i *p*=0.0006, and *ATGL*-KO untreated vs ACC1i *p*=0.9331.

**Figure 5** exact *p*-values are: (**c**) WT vs TAG-TKO *p*=0.0005; (**d**) WT vs TAG-TKO *p*=0.8283; (**e**) control vs ATGLi: 2 hr ACC1i *p=*0.0012, 4 hr ACC1i *p*<0.0001, and 6 hr ACC1i *p*<0.0001; (**f**) WT control vs ACC1i *p*<0.0001, WT ACC1i vs ACC1i + ATGLi *p*<0.0001, TAG-TKO control vs ACC1i *p*<0.0001, and TAG-TKO ACC1i vs ACC1i + ATGLi *p*<0.0001; (**g**) WT untreated vs ATGLi *p*=0.0045 and TAG-TKO untreated vs ATGLi *p*=0.0062.

**Figure 6** exact *p*-values are: (**j**) untreated vs ACC1i *p*=0.0378, untreated vs ATGLi *p*=0.0124, untreated vs DHA *p*<0.0001, untreated vs MβCD *p*<0.0001, MβCD vs ACC1i + MβCD *p*<0.0001, MβCD vs ATGLi + MβCD *p*=0.1111 and MβCD vs DHA + MβCD *p*<0.0001; and (**l**) untreated vs ACC1i *p*<0.0001, untreated vs ATGLi *p*=0.0026, untreated vs DHA *p*<0.0001 and untreated vs MβCD *p*=0.0090.

**Figure 7** exact *p*-values are: (**c**) untreated vs myriocin *p*<0.0001, untreated vs oleic acid *p*<0.0001, and untreated vs ACC1i *p*<0.0001; (**e**) WT vs dLKO *p*<0.0001; (**g**) WT vs dLKO *p*=0.0004; (**k**) WT vs dLKO *p*<0.0001; (**l**) WT vs dLKO *p=*0.1915; (**m**) WT vs dLKO *p*=0.0434.

## Data Availability

All data generated in this manuscript is available in a public repository, and source data are also provided with this paper. Raw data from the random insertion mutagenesis screen can be found at the NCBI Sequence Read Archive (SRA) with the BioProject ID: PRJNA1266240 [<u>1</u>]. The results of the analyzed screens are available as a source data file. Raw data for the metabolomics can be found at Metabolomics Workbench with the study ID ST004029 [<u>2</u>]. The lipidomics data used to generate all figures is included in Source Data Files. Raw data for the RNA sequencing can be found at NCBI GEO with the accession number: GSE298034 [<u>3</u>].

## Acknowledgements

We thank Dr. Giovanni Luchetti, Dr. Yvonne Lange, Dr. Theodore Steck, and Dr. Pehr Harbury for helpful discussions.

## Core facilities

This work was supported by the Vincent Coates Foundation Mass Spectrometry Laboratory, Stanford University Mass Spectrometry (RRID:SCR_017801) utilizing the Sciex 7500+ LC/MS system (RRID:SCR_026608). This work was also supported by the Stanford University Cell Sciences Imaging Core Facility (RRID: SCR_017787) utilizing the Leica SP8 Confocal which was supported, in part, by Award Number 1S10OD010580 from the National Center for Research Resources (NCRR).

## Author Contributions

KMW performed experimental studies, carried out data analysis, generated figures and contributed to writing this manuscript. CK performed experimental studies. SC carried out experimental studies and contributed to writing this manuscript. EOS carried out experimental studies and generated figures. ET carried out experimental studies. SL contributed to the experimental design of this manuscript. CBK carried out experimental studies and data analysis. DC contributed to data interpretation and data analysis. MD carried out experimental studies. RL supervised mass spectrometry experimental studies. JGB contributed to data interpretation and data analysis. DT contributed to data analysis. AML contributed to experimental design and carried out data analysis. JAO contributed to experimental design, data interpretation, and writing this manuscript. RR supervised experimental studies and contributed to data interpretation and analysis. MK supervised experimental studies, performed experimental studies, carried out data analysis, generated figures, and contributed to the writing and planning of this manuscript.

