## Supplementary Figures and Tables for "Fatty acid metabolism controls plasma membrane cholesterol accessibility via ATGL-dependent phospholipid remodeling"

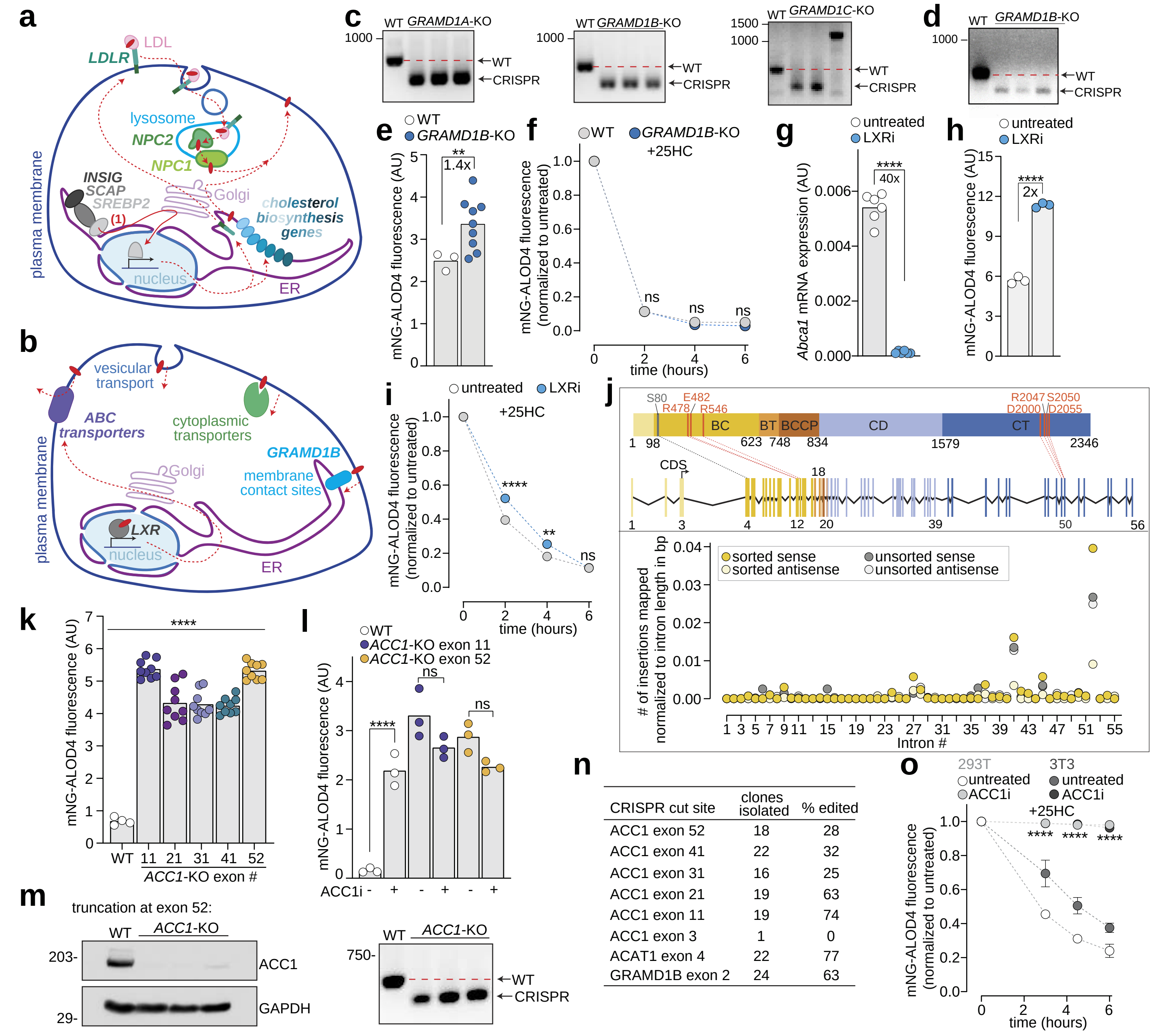


| CRISPR cut site | clones isolated | % edited |
|---|---|---|
| ACC1 exon 52 | 18 | 28 |
| ACC1 exon 41 | 22 | 32 |
| ACC1 exon 31 | 16 | 25 |
| ACC1 exon 21 | 19 | 63 |
| ACC1 exon 11 | 19 | 74 |
| ACC1 exon 3 | 1 | 0 |
| ACAT1 exon 4 | 22 | 77 |
| GRAMD1B exon 2 | 24 | 63 |

**Supplementary Figure 1. a** Schematic showing overview of cholesterol biosynthesis and uptake pathways. When ER cholesterol is low, SCAP is released from INSIG tethering at the ER. SCAP escorts SREBP2 to the Golgi (solid red line) where SREBP2 is proteolytically cleaved into a transcriptionally active protein fragment. SREBP2 target genes include the >20 genes involved in cholesterol synthesis and LDLR. LDL-derived cholesterol is transported out of the lysosome by NPC2 and NPC1. **b** Schematic showing that cholesterol can be internalized from the PM by membrane contact site proteins like GRAMD1B, by cytoplasmic transporters, or by vesicular transport. Cholesterol is exported from the PM by ABC transporters, whose expression is elevated in response to LXR activation by sterols. **c**,**d** PCR analysis of *GRAMD1A/B/C*-TKO clones (**c**) and *GRAMD1B*-KO clones (**d**). DNA gels were repeated three times with the same results. **e**,**f** Flow cytometry analysis of PM accessible cholesterol in WT and *GRAMD1B*-KO cells either left untreated (**e**) or treated with 25HC for the indicated times (**f**). **g** Quantitative real-time PCR analysis of *Abca1* mRNA levels normalized to *Gapdh* from 3T3 cells treated for 20 hrs with LXRi (1 μM GSK2033). **h**,**i** Flow cytometry analysis of PM accessible cholesterol in 3T3 cells treated for 20 hrs with or without 1 μM of LXRi alone (**h**) or followed by treatment with 25HC for the indicated times (**i**). **j** Domain architecture of ACC1 (top) including the biotin carboxylase domain (BC), biotin transferase domain (BT), biotin carboxyl-carrier protein domain (BCCP), central domain (CD), and the carboxyltransferase domain (CT). Residues implicated in catalysis are shown in dark orange. An exon map of ACC1 with exon colors matching the domain they encode is shown below. The number of insertions mapped per intron (bottom) are normalized to intron length in base pairs (bp). **k**,**l** Flow cytometry analysis of PM accessible cholesterol in WT and *ACC1*-KO cells made with CRISPR sgRNAs targeting the indicated exons either left untreated (**k**) or treated with ACC1i (30 μM Firsocostat) for 20 hrs (**l**). **m** Western blot (left) and PCR analysis (right) of WT HAP1 and *ACC1*-KO cells. Western blot and DNA gels were repeated at least three times with similar results. **n** Table showing the number of clones isolated from two 96-well dishes and the percentage of those clones that were found to be knockouts with PCR analysis. **o** Flow cytometry analysis of PM accessible cholesterol in cells either left untreated or treated for 16 hrs with ACC1i and then treated with 4 μM 25HC for the times indicated. In **f**, **i**, and **o**, each data point is the mean of three independent biological replicates. Each circle is the median value for a biological replicate, with n=5000 cells analyzed per replicate (**e**,**g**,**h**,**k**, and **l**). In **f**, **i** and **o**, data is normalized to cells not treated with 25HC for each condition, and each circle is the mean of three biological replicates, with n=5000 cells per replicate; error bars are the standard deviation. Cells in **k** and **l** were grown in 5% LDS for 16 hours prior to analysis. Statistical significance was determined by a two-tailed Welch's t-test (**e**,**g** and **h**), a two-way ANOVA followed by Sidak's multiple comparison test (**f**,**i**,**l** and **o**) or a Dunnett's one-way ANOVA (**k**). Exact *p*-values: (**e**) $p$=0.0043; (**f**) WT vs *GRAMD1B*-KO: 2 hr ACC1i $p$>0.9999, 4 hr ACC1i $p$=0.7491 and 6 hr ACC1i $p$=0.5126; (**g**) $p$<0.0001; (**h**) $p$<0.0001; (**i**) control vs LXRi: 2 hr 25HC $p$<0.0001, 4 hr 25HC $p$=0.0004 and 6 hr 25HC $p$>0.9999; (**k**) WT vs *ACC1*-KO: exon 11 $p$<0.0001, exon 21 $p$<0.0001, exon 31 $p$<0.0001, exon 41 $p$<0.0001 and exon 52 $p$<0.0001; (**l**) WT untreated vs ACC1i $p$<0.0001, *ACC1*-KO exon 11 untreated vs ACC1i $p$=0.0508 and *ACC1*-KO exon 52 untreated vs ACC1i $p$=0.0701 and (**o**) 3T3 control vs ACC1i: 3 hr 25HC $p$<0.0001, 4.5 hr 25HC $p$<0.0001 and 6 hr 25HC $p$<0.0001; 293T control vs ACC1i: 3 hr 25HC $p$<0.0001, 4.5 hr 25HC $p$<0.0001 and 6 hr 25HC $p$<0.0001. AU is arbitrary unit.

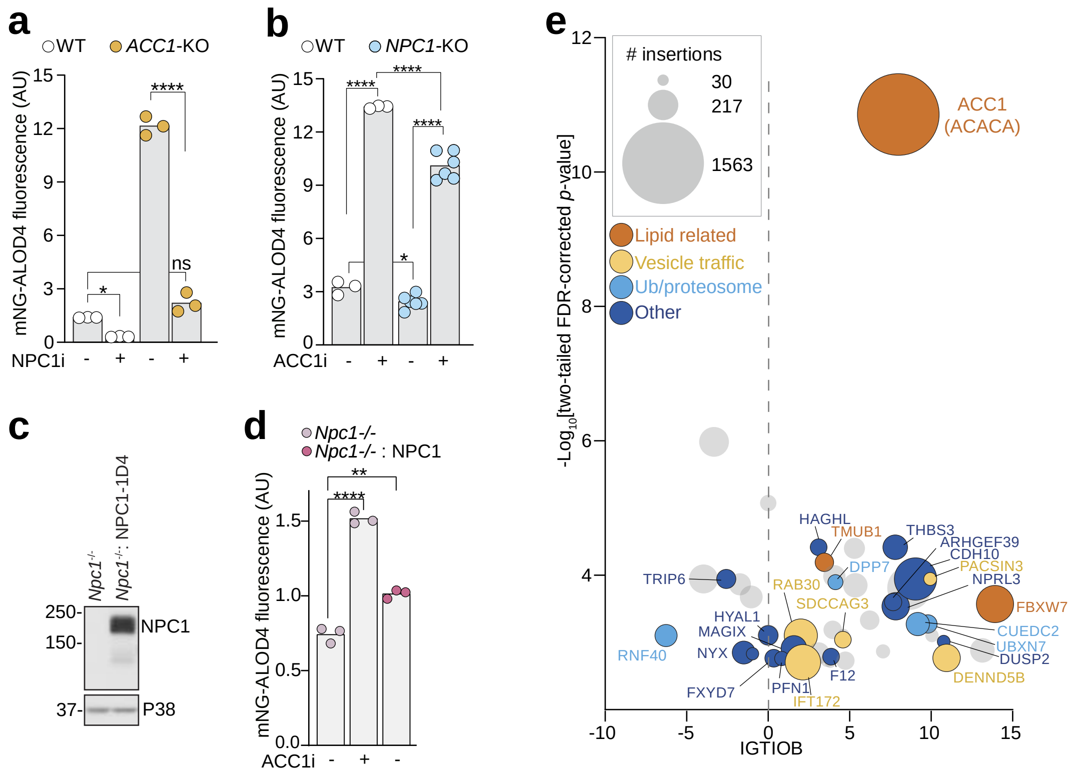


**Supplementary Figure 2. a**,**b** Flow cytometry analysis of PM accessible cholesterol in WT or *ACC1*-KO HAP1 cells treated with or without NPC1i (1 μM U18666A) for 20 hrs (**a**) or WT and *NPC1*-KO HAP1 cells treated with or without ACC1i (30 μM Firsocostat) for 16 hrs (**b**). **c**,**d** Western blot analysis (**c**) or mNG-ALOD4 flow cytometry analysis of PM accessible cholesterol (**d**) of *Npc1*-/- mouse embryonic fibroblasts (MEFs) or *Npc1*-/- MEFs with NPC1 stably re-introduced (*Npc1*-/-: NPC1) using MSCV retrovirus, and then treated with or without ACC1i for 20 hrs. The experiment was repeated independently three times with simlar results. Each circle is the median value for a biological replicate, with n=5,000 cells analyzed per replicate (**a**,**b** and **d**). **e** Screen results showing enriched genes with a two-tailed FDR-corrected *p*-value of less than 2E-4. IGTIOB is a measure of the inactivating potential of the mapped insertions, and circle size shows the number of insertions for each gene; genes are colored based on their proposed function. Statistical significance was determined by a two-way ANOVA followed by Sidak's multiple comparison test (**a**,**b** and **d**). Exact *p*-values are: (**a**) WT untreated vs NPC1i *p*=0.0459, *ACC1*-KO untreated vs NPC1i *p*<0.0001, WT untreated vs *ACC1*-KO + NPC1i p=0.1483; (**b**) WT untreated vs *NPC1*-KO untreated *p*=0.0211, and all other comparisons are *p*<0.0001; and (**d**) *Npc1*-/- untreated vs ACC1i *p*<0.0001 and *Npc1*-/- vs *Npc1*-/-: NPC1 *p*=0.0004. AU is arbitrary unit.

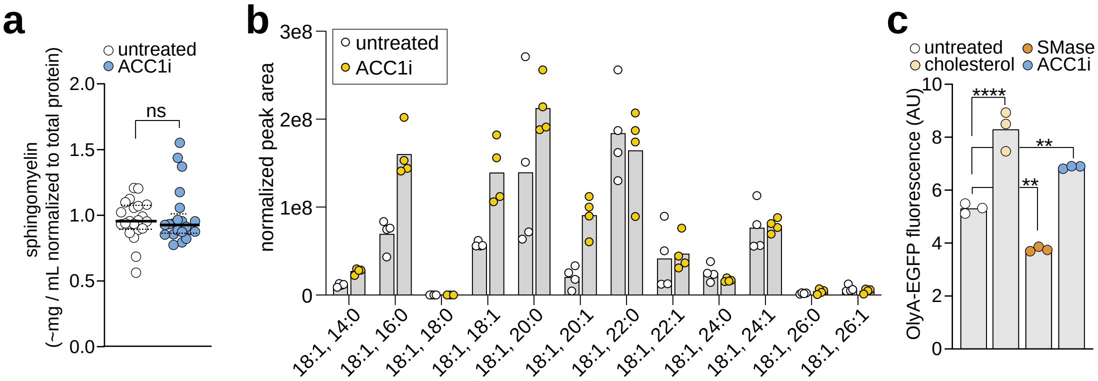


**Supplementary Figure 3. a**, MS analysis of sphingomyelin levels in WT HAP1 cells either left untreated or treated with ACC1i (30 µM Firsocostat) for 16 hrs. This MS dataset was gathered by Metabolon Inc. **b** MS normalized peak areas of sphingomyelin species in untreated and ACC1i treated cells. This MS dataset was performed in the Stanford University Mass Spectrometry facility. **c** OlyA-EGFP flow cytometry analysis of WT HAP1 cells left untreated or treated with 0.3 mM MβCD:cholesterol for 30 minutes, 150 mU/mg SMase for 30 minutes, or ACC1i for 16 hrs. Statistical significance was determined by a two-tailed Welch's t-test (**a**) or a Dunnett's one-way ANOVA (**c**). Each circle is the median value for a biological replicate, with n=5,000 cells analyzed per replicate (**c**).Exact *p*-values are: (**a**) untreated vs ACC1i $p$=0.6634; and (**c**) untreated vs cholesterol $p$<0.0001, untreated vs SMase $p$=0.0033 and untreated vs ACC1i $p$=0.0031. AU is arbitrary unit.

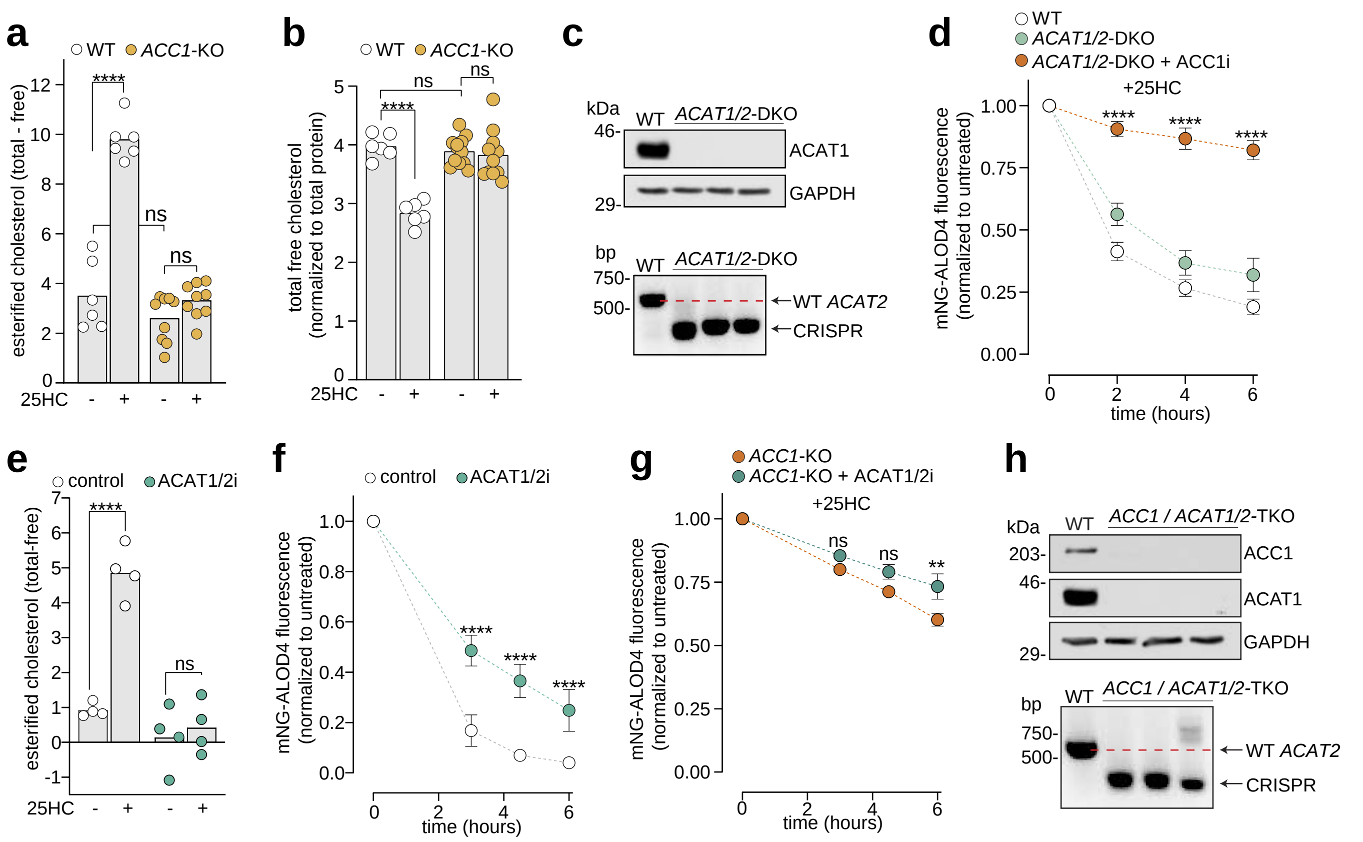


**Supplementary Figure 4. a**, **b** Analysis of the esterified (**a**) and total cholesterol (**b**) content of WT and *ACC1*-KO HAP1 cells either left untreated or treated with 4 μM 25HC for 8 hours. For (**a**), n= 6, 6, 9 and 9 independent biological replicates for WT untreated, WT +25HC, ACC1-KO untreated and ACC1-KO +25HC respectively. For (**b**), n= 6, 6, 12 and 12 independent biological replicates respectively and each sample was measured as a technical triplicate. **c** Western blot (ACAT1) and PCR (*ACAT2*) analysis of *ACAT1/2*-DKO clones. The experiment was repeated independently three times with similar results. **d** Cells were left untreated or treated for 16 hrs with ACC1i (30 μM Firsocostat, orange curve) and then subsequently treated with 4 μM 25HC for the times indicated. PM accessible cholesterol was then analyzed by flow cytometry analysis of mNG-ALOD4 staining. **e** Analysis of esterified cholesterol in WT HAP1 cells treated with or without ACAT1/2 inhibitor (ACAT1/2i; 60 μM SZ58-035) for 16 hrs, followed by a 6 hr treatment with or without 4 μM 25HC. For (**e**), n= 4 independent biological replicates per condition, with each sample measured in technical triplicate. **f**,**g** Flow cytometry analysis of PM accessible cholesterol in cells left untreated or treated for 16 hours with ACAT1/2i followed by 4 μM 25HC treatment for the indicated times. **h** Western blot (ACAT1 and ACC1) or PCR (*ACAT2*) analysis of *ACC1/ACAT1/ACAT2*-TKO clones. The experiment was repeated independently three times with similar results. For each condition (**d**,**f** and **g**), data is normalized to cells not treated with 25HC, and each data point represents the average of three biological replicates with n=5000 cells per replicate; error bars denote the standard deviation. Statistical significance was determined by a two-way ANOVA followed by Sidak's multiple comparison test (**a**,**b**, **d**-**g**). Exact *p*-values are: (**a**) WT untreated vs *ACC1*-KO untreated $p$=0.2143, WT untreated vs 25HC $p$<0.0001, and *ACC1*-KO untreated vs 25HC $p$=0.1296; (**b**) WT untreated vs *ACC1*-KO untreated $p$=0.8909, WT untreated vs 25HC $p$<0.0001, and *ACC1*-KO untreated vs 25HC $p$=0.9227; (**d**) all conditions: $p$<0.0001; (**e**) control vs 25HC $p$<0.0001 and ACAT1/2i without vs with 25HC $p$=0.8244; (**f**) all conditions: $p$<0.0001; (**g**) *ACC1*-KO vs *ACC1*-KO + ACAT1/2i: 3 hr 25HC $p$=0.3127, 4.5 hr 25HC $p$=0.0818 and 6 hr 25HC $p$=0.0020.

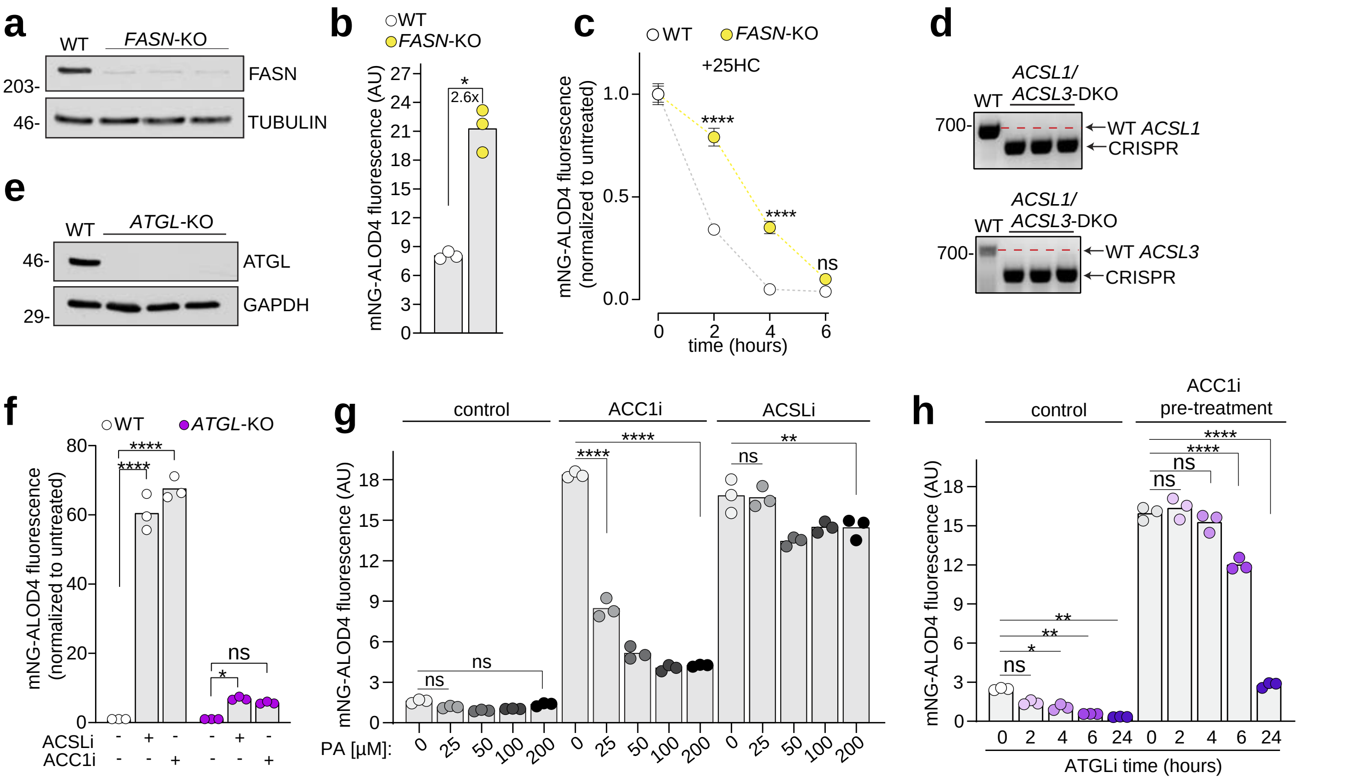


**Supplementary Figure 5. a** Western blot analysis of WT HAP1 cells compared to *FASN*-KO HAP1 cells. The experiment was repeated independently three times with similar results. **b**,**c** Flow cytometry analysis of mNG-ALOD4 PM staining in WT and *FASN*-KO cells grown in 5% LDS serum media for 24 hrs and then left untreated (**b**) or treated with 4 μM 25HC for the indicated times (**c**). **d** PCR analysis of *ACSL1/ACSL3* double knockout (*ACSL1/ACSL3*-DKO) cells compared to WT cells. The experiment was repeated independently three times with similar results. **e** Western blot analysis of *ATGL*-KO HAP1 cells compared to WT cells. The experiment was repeated independently three times with similar results. **f**-**h** Flow cytometry analysis of PM cholesterol in cells treated with or without ACC1i (30 μM Firsocostat) (**f**-**h**) or ACSLi (5 μM Triacsin-C) for 16 hrs (**f**,**g**), and then treated with different palmitic acid (PA) concentrations for 6 hrs (**g**), or ATGLi (50 μM NG-497) for the indicated times (**h**). Cells were grown in 5% LDS serum media for 16 hours prior to drug treatments (**f**-**h**). Each circle is the median value for a biological replicate, with n=5,000 cells analyzed per replicate (**b**, **f**-**h**). In **c**, each data point represents the average of three biological replicates with n=5000 cells per replicate; error bars denote the standard deviation. Statistical significance was determined by a two-tailed Welch's t-test (**b**), or a two-way ANOVA followed by Sidak's multiple comparison test (**c** and **f**-**h**). Exact *p*-values are: (**b**) WT vs *FASN*-KO $p=0.0080$; (**c**) WT vs *FASN*-KO: 2 hr 25HC $p<0.0001$, 4 hr 25HC $p<0.0001$ and 6 hr 25HC $p=0.1922$; (**f**) WT untreated vs ACSLi $p<0.0001$, WT untreated vs ACC1i $p<0.0001$, *ATGL*-KO untreated vs ACSLi $p=0.0279$, *ATGL*-KO untreated vs ACC1i $p=0.0827$; (**g**) control 0 μM PA vs 25 μM PA $p=0.9614$, control 0 μM PA vs 200 μM PA $p=0.9982$, ACC1i 0 μM PA vs 25 μM PA $p<0.0001$, ACC1i 0 μM PA vs 200 μM PA $p<0.0001$, ACSLi 0 μM PA vs 25 μM PA $p>0.9999$, and ACSLi 0 μM PA vs 200 μM PA $p=0.0018$; (**h**) control 0 vs 2 hr ATGLi $p=0.2657$, control 0 vs 4 hrs ATGLi $p=0.0307$, control 0 vs 6 hrs ATGLi $p=0.0012$, control 0 vs 24 hrs ATGLi $p=0.0003$, ACC1i 0 vs 2 hr ATGLi $p>0.9999$, ACC1i 0 vs 4 hr ATGLi $p=0.9533$, ACC1i 0 vs 6 hr ATGLi $p<0.0001$, and ACC1i 0 vs 24 hrs ATGLi p<0.0001. AU is arbitrary unit.

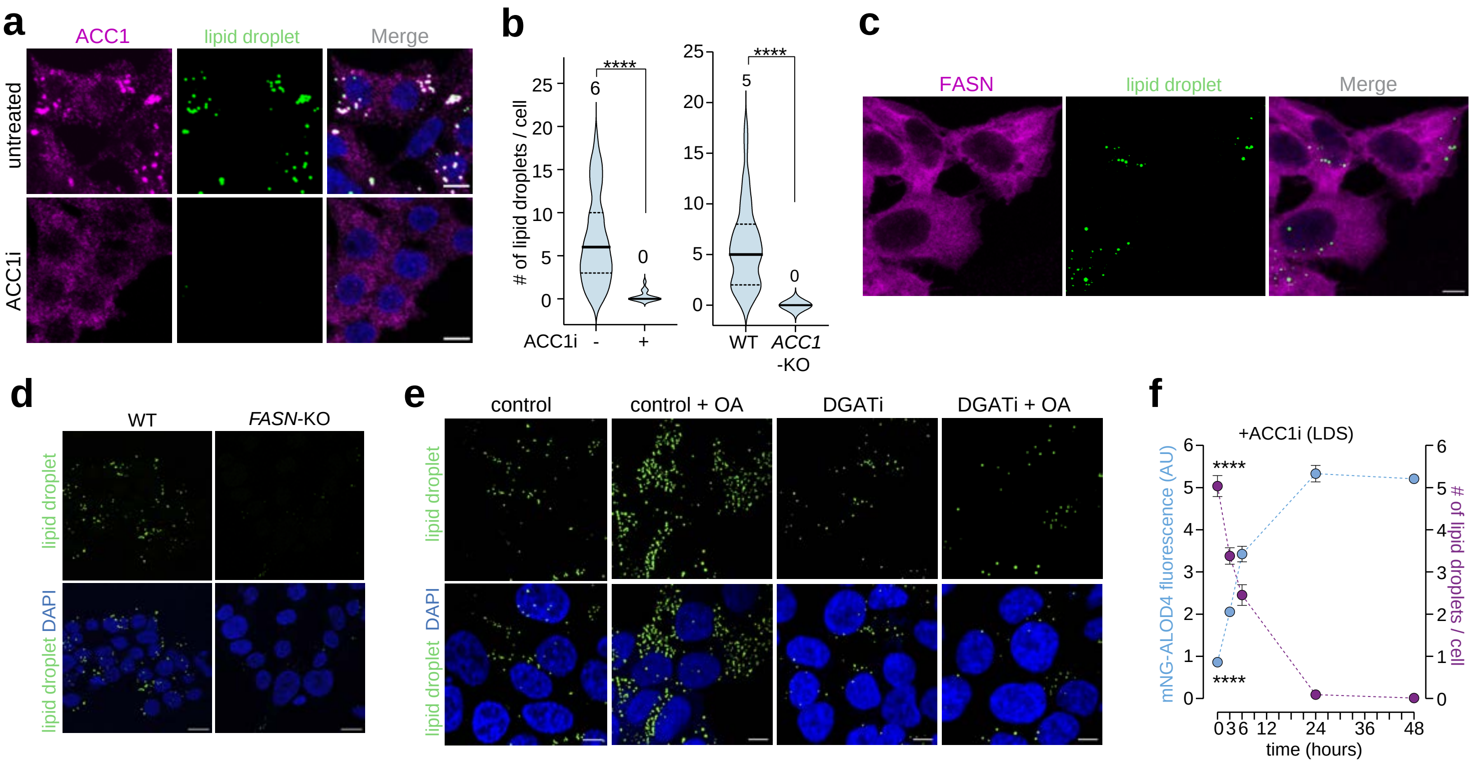


**Supplementary Figure 6. a** Microscopy images of HAP1 cells left untreated or treated with ACC1i (30 µM Firsocostat, 16 hrs), and then stained with 647-conjugated ACC1 polyclonal antibody, BODIPY 493/503 to measure LDs and DAPI (blue); the scale bar is 5 microns. **b** Quantification of LD numbers per cell in WT HAP1 cells treated with or without ACC1i (n= 92 and 97 cells, respectively; left) or WT HAP1 cells compared to *ACC1*-KO HAP1 cells (n= 77 and 57 cells, respectively; right). The solid middle line shows the median and dashed lines show the interquartile range. **c** Immunofluorescence microscopy image of untreated WT HAP1 cells stained with a FASN antibody, BODIPY 493/503 and DAPI (blue); the scale bar is 5 microns. **d**,**e** Microscopy images of LDs in WT HAP1 and *FASN*-KO cells (**d**) and WT HAP1 cells left untreated or treated with DGAT1 and DGAT2 inhibitors (DGATi) for 16 hrs followed by oleic acid (150 µM OA) treatment for 6 hrs (**e**); the scale bar is 10 microns. **f** Flow cytometry analysis of PM cholesterol (blue curve), with each circle representing the median value of a biological replicate (n=5,000 cells analyzed per replicate), and the corresponding fluorescence microscopy quantification of the number of LDs per cell (purple curve; n=75 cells examined per field of view across 3 fields of view and each data point represents the total number of LDs per cell in a single field of view) in HAP1 cells grown in 5% LDS and then treated with ACC1i for the indicated times. Statistical significance was determined by a two-tailed Welch's t-test (**b**) or a Dunnett's one-way ANOVA (**f**). Exact *p*-values are: (**b**) WT vs ACC1i $p<0.0001$ and WT vs *ACC1*-KO $p<0.0001$; (**f**) PM cholesterol $p<0.0001$ and LD number $p<0.0001$ across all treatment times. AU is arbitrary unit.

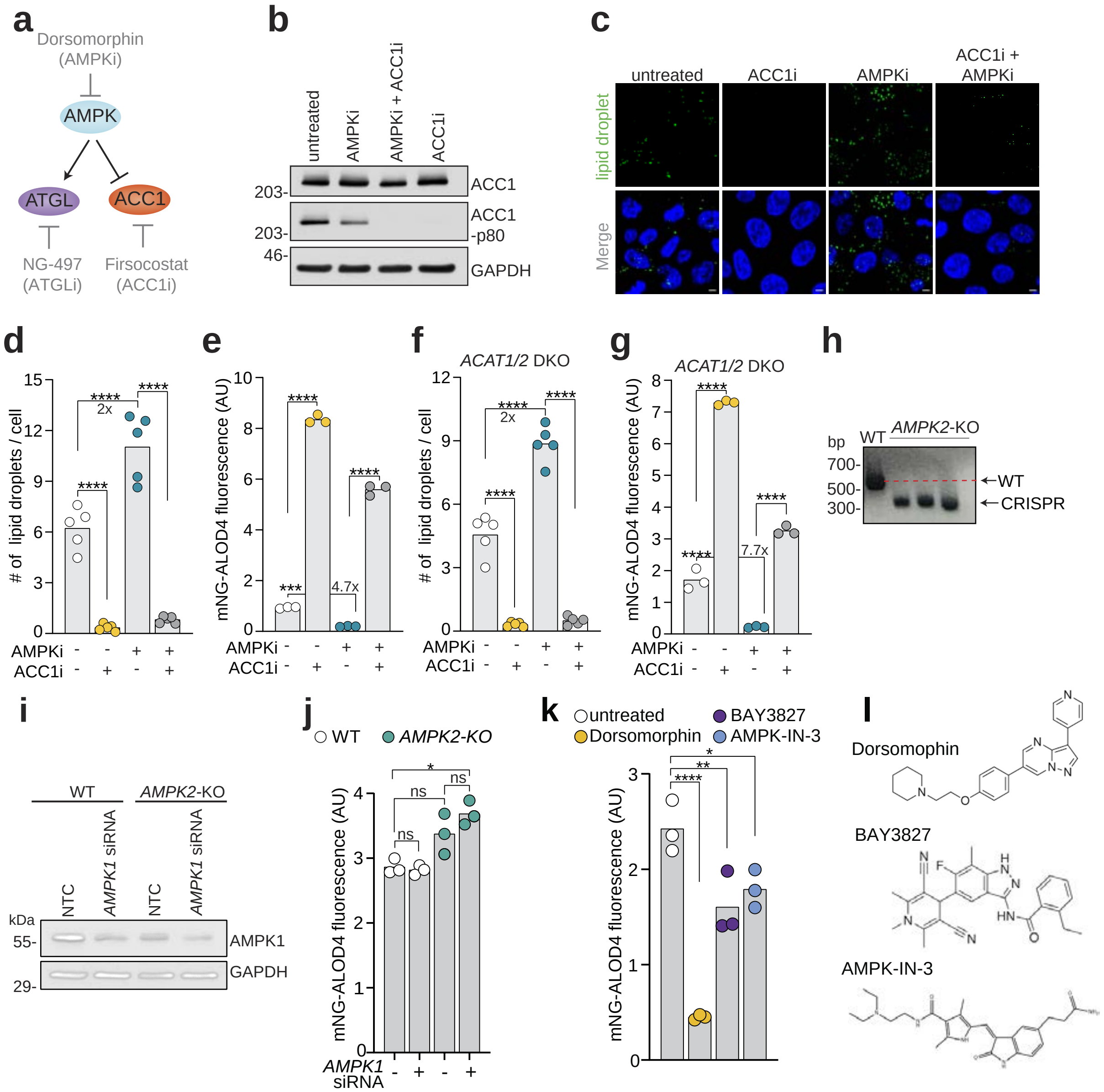


**Supplementary Figure 7. a** Schematic showing that AMPK activates ATGL and inactivates ACC1. Inhibitors for each enzyme are shown in gray. **b** Western blot analysis of ACC1 phosphorylation in WT HAP1 cells left untreated or treated with AMPKi (2.5 µM Dorsomorphin), ACC1i (30 µM Firsocostat) or both. The experiment was repeated independently three times with similar results. **c**-**e** WT HAP1 cells were left untreated or treated with ACC1i, AMPKi or both and then analyzed by microscopy to quantify LD numbers per cell (**c**, **d**) or by mNG-ALOD4 flow cytometry analysis to determine PM cholesterol levels (**e**); the scale bar is 5 microns. **f**,**g** LD quantification (**f**) and flow cytometry analysis of PM cholesterol using mNG-ALOD4 staining (**g**) in *ACAT1/2*-DKO cells left untreated or treated with AMPKi, ACC1i or both. **h** PCR analysis of *AMPK2* knockout (KO) clones. **i** Western blot analysis showing siRNA knockdown of AMPK1 or a non-targeting control (NTC) in WT and *AMPK2*-KO cells. The experiment was repeated independently three times with similar results. **j** Flow cytometry analysis of PM cholesterol in WT or *AMPK2*-KO cells with or without AMPK1 siRNA or a NTC. **k** Flow cytometry of PM accessible cholesterol levels after cells were treated for 20 hours with three distinct AMPK inhibitors (2.5 µM Dorsomorphin, 5 µM BAY3827 and 10 µM AMPK-IN-3). **l** The structure of each AMPK inhibitor. All cells were grown in lipoprotein depleted serum media for 16 hours prior to drug treatments. n=75 cells examined per field of view across 3 fields of view and each data point represents the total number of LDs per cell in a single field of view (**d**,**f**). Each circle is the median value for a biological replicate, with n=5,000 cells analyzed per replicate (**e**,**g**, **j** and **k**). Statistical significance was determined by one-way ANOVA followed by Sidak's multiple-comparisons test (**d**-**g**), a two-way ANOVA with Sidak's multiple comparison test (**j**), or by a Dunnett's one-way ANOVA (**k**). Exact *p*-values in are: (**d**) untreated vs ACC1i $p<0.0001$, untreated vs AMPKi $p<0.0001$, AMPKi vs ACC1i + AMPKi $p<0.0001$; (**e**) untreated vs ACC1i $p<0.0001$, untreated vs AMPKi $p=0.0005$, and AMPKi vs ACC1i + AMPKi $p<0.0001$; (**f**) *ACAT1/2*-DKO untreated vs ACC1i $p<0.0001$, untreated vs AMPKi $p<0.0001$, and AMPKi vs ACC1i+ AMPKi $p<0.0001$; (**g**), *ACAT1/2*-DKO untreated vs ACC1i $p<0.0001$, untreated vs AMPKi $p<0.0001$, and AMPKi vs ACC1i + AMPKi $p<0.0001$; (**j**) WT NTC siRNA vs AMPK1 siRNA $p=0.9999$, WT vs *AMPK2*-KO (NTC siRNA) $p=0.0967$, WT (NTC siRNA) vs *AMPK2*-KO with AMPK1 siRNA $p=0.0110$, and *AMPK2*-KO cells NTC vs AMPK1 siRNA $p=0.4357$; (**k**) untreated vs Dorsomorpin $p<0.0001$, untreated vs BAY3827 $p=0.0081$, and untreated vs AMPK-IN-3 $p=0.0319$. AU is arbitrary units.

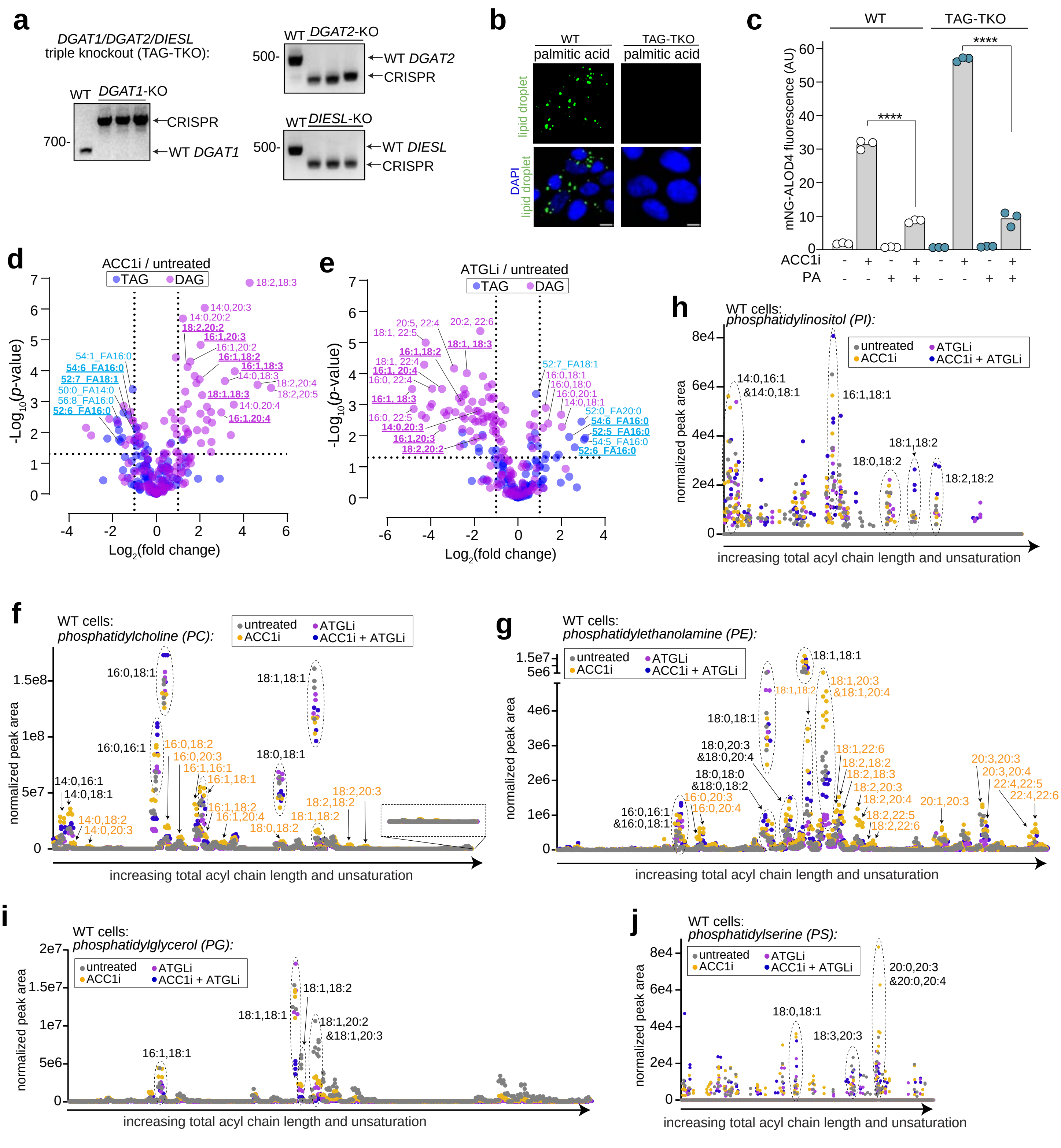

**Supplementary Figure 8. a** PCR analysis of WT HAP1 cells compared to TAG-TKO clones. **b** Microscopy images of LDs in WT and TAG-TKO cells left untreated or treated with 200 μM palmitic acid for 6 hrs. The scale bar is 5 microns. **c** Flow cytometry analysis of PM accessible cholesterol in WT and TAG-TKO cells left untreated or treated with ACC1i (30 μM Firsocostat) for 16 hrs followed by treatment with palmitic acid (200 μM PA) for 6 hrs. Each circle is the median value for a biological replicate, with n=5,000 cells analyzed per replicate. **d**-**j** MS analysis of TAGs and DAGs in TAG-TKO cells (**d**,**e**), or MS analysis of WT cells showing: phosphatidylcholine (**f**), phosphatidylethanolamine (**g**), phosphatidylinositol (**h**), phosphatidylglycerol (**i**), and phosphatidylserine (**j**) with the indicated treatments. The x-axis lists all lipid species in order of increasing acyl chain length and unsaturation (**f**-**j**). Select lipid species are denoted by a dotted oval and specification of the acyl chain composition. Statistical significance was determined by a two-way ANOVA followed by Sidak's multiple comparison test (**c**). Exact *p*-values are: (**c**) WT: ACC1i vs ACC1i+PA $p<0.0001$, TAG-TKO: ACC1i vs ACC1i+PA $p<0.0001$. AU is arbitrary unit.

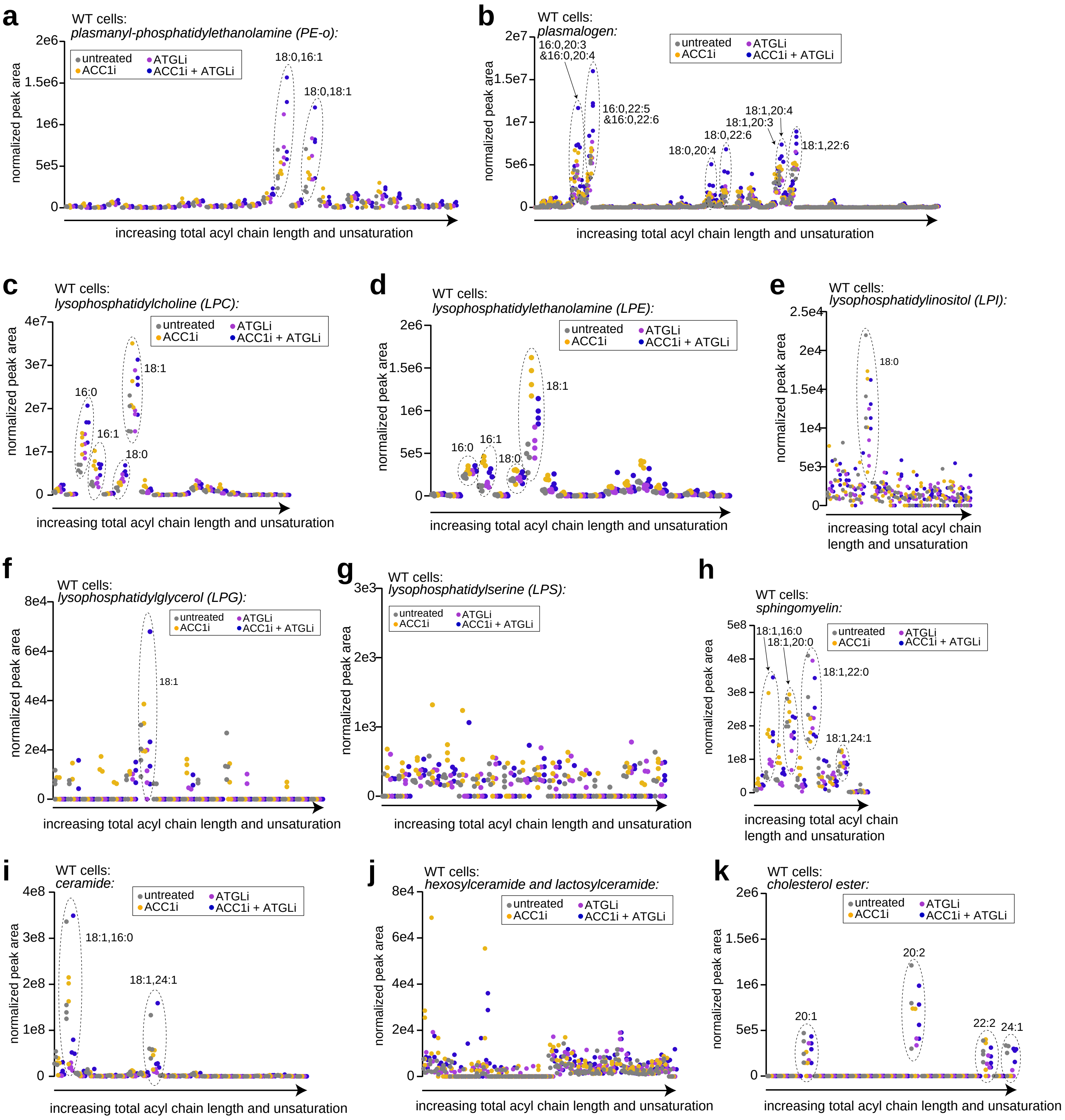


**Supplementary Figure 9. a-i** MS analysis of lipid species in WT HAP1 cells showing: plasmanyl-phosphatidylethanolamine (**a**), plasmalogen (**b**), lysophosphatidylcholine (**c**), lysophosphatidylethanolamine (**d**), lysophosphatidylinositol (**e**), lysophosphatidylglycerol (**f**), lysophosphatidylserine (**g**), sphingomyelin (**h**), ceramide (**i**), hexosylceramide and lactosylceramide (**j**), and cholesterol ester (**k**) with the indicated treatments. The x-axis lists all lipid species in order of increasing acyl chain length and unsaturation. Select lipid species are denoted by a dotted oval and specification of the acyl chain composition.

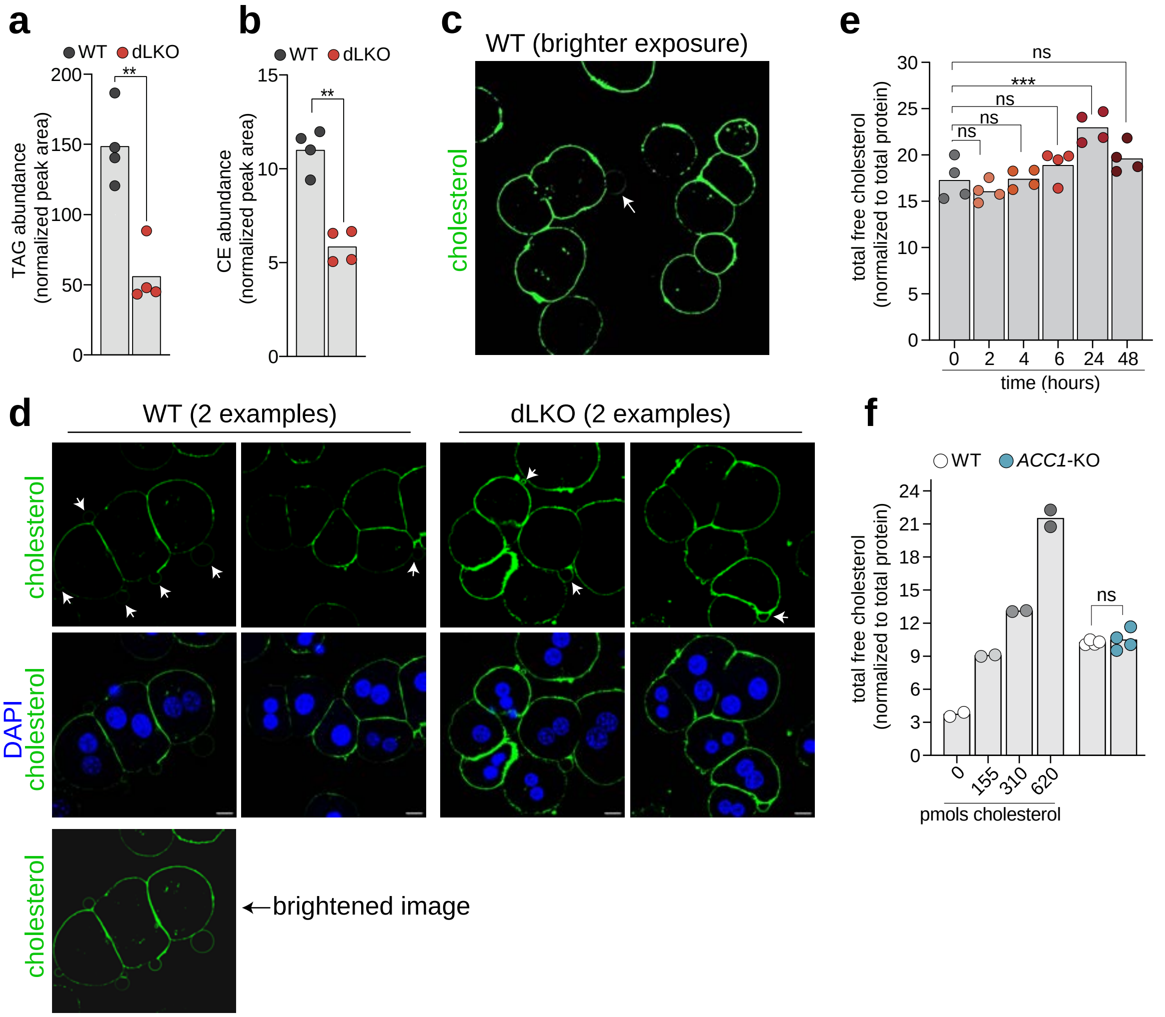


**Supplementary Figure 10. a**,**b** Hepatic TAG levels (**a**), and hepatic CE levels (**b**) in male *Acc1/Acc2 flox/flox* control and male *Albumin-Cre-Acc1/Acc2* double liver knockout (dLKO) mice. **c** A brightened image of **Fig. 7f**, showing that control primary hepatocytes have membrane blebbing similar to dLKO hepatocytes. **d** Additional examples of fluorescence microscopy images of primary hepatocytes stained with mNG-ALOD4 and DAPI. Membrane blebs are seen in both control and dLKO hepatocytes. The scale bar is 10 microns. **e,f** Amplex Red Assay quantification of free (unesterified) cholesterol normalized to total protein in WT HAP1 cells treated with ACC1i (30 μM Firsocostat) for the indicated times or for 16 hours (**f**). n=4 independent biological replicates per time point (**e**) or per condition (**f**), with each data point representing the median of triplicate technical measurements for each sample. Statistical significance was determined by a two-tailed Welch's t-test (**a**,**b** and **f**) or by a Dunnett's one-way ANOVA (**e**). Exact *p*-values are: (**a**) WT vs dLKO *p*=0.0022, (**b**) WT vs dLKO *p*=0.0005, (**e**) 0 vs 2 hrs *p*=0.7222, 0 vs 4 hrs *p*>0.9999, 0 v 6 hrs *p*=0.4782, 0 vs 24 hrs *p*=0.0004, and 0 vs 48 hrs *p*=0.1813; (**f**) WT vs *ACC1*-KO *p*>0.9999. AU is arbitrary unit.

Supplementary Table 1. Oligonucleotides used in this study

| Oligonucleotide | Application | Sequence (5′–3′) |
|---|---|---|
| ACC1 Ex3, Guide 1 Forward | CRISPR guide | caccgtttggggatctctagcctac |
| ACC1 Ex3, Guide 1 Reverse | CRISPR guide | aaacgtaggctagagatccccaaac |
| ACC1 Ex3, Guide 2 Forward | CRISPR guide | caccgcaagctggagtgcagtggtg |
| ACC1 Ex3, Guide 2 Reverse | CRISPR guide | aaaccaccactgcactccagcttgc |
| ACC1 Ex3, Forward | Genotyping | ggagctgaaccagcactctc |
| ACC1 Ex3, Reverse | Genotyping | gtggcaggcgcctgtaatc |
| ACC1 Ex11, Guide 1 Forward | CRISPR guide | caccggtgattgctctgtacaacgc |
| ACC1 Ex11, Guide 1 Reverse | CRISPR guide | aaacgcgttgtacagagcaatcacc |
| ACC1 Ex11, Guide 2 Forward | CRISPR guide | caccggcgtaccatttcaggataca |
| ACC1 Ex11, Guide 2 Reverse | CRISPR guide | aaactgtatcctgaaatggtacgcc |
| ACC1 Ex11, Forward | Genotyping | gctgaagttcctgga tct ccc |
| ACC1 Ex11, Reverse | Genotyping | ccgaaggcatcgctcatgac |
| ACC1Ex21, Guide1 Forward | CRISPR guide | caccgttcacacaggtagtctgcca |
| ACC1Ex21, Guide1 Reverse | CRISPR guide | aaactggcagactacctgtgtgaac |
| ACC1Ex21, Guide2 Forward | CRISPR guide | caccgcttttttttttcctcgaga |
| ACC1Ex21, Guide2 Reverse | CRISPR guide | aaactctcgaggaaaaaaaaaagc |
| ACC1Ex21, Forward | Genotyping | gtgcagtggtgtgtttgtacct |
| ACC1Ex21, Reverse | Genotyping | gcacgtgcctacaatcccc |
| ACC1Ex31, Guide1 Forward | CRISPR guide | caccgtagtgtcagcgatgtactgt |
| ACC1Ex31, Guide1 Reverse | CRISPR guide | aaacacagtacatcgctgacactac |
| ACC1Ex31, Guide2 Forward | CRISPR guide | caccgtgctgtatatgccgttggag |
| ACC1Ex31, Guide2 Reverse | CRISPR guide | aaacctccaacggcatatacagcac |
| ACC1Ex31, Forward | Genotyping | cccagtccttgctcctaccc |
| ACC1Ex31, Reverse | Genotyping | ctcatcacagagcagacagcttc |
| ACC1Ex41, Guide1 Forward | CRISPR guide | caccggaactggtactggatgatca |
| ACC1Ex41, Guide1 Reverse | CRISPR guide | aaactgatcatccagtaccagttcc |
| ACC1Ex41, Guide2 Forward | CRISPR guide | caccgatagtattgttttaacacaa |
| ACC1Ex41, Guide2 Reverse | CRISPR guide | aaacttgtgttaaaacaatactatc |
| ACC1Ex41, Forward | Genotyping | tgggagtctatgtccactcaagc |
| ACC1Ex41, Reverse | Genotyping | acacccaccctcacacctc |
| ACC1Ex52, Guide1 Forward | CRISPR guide | caccgggtgaaaaccatgcgtcggg |
| ACC1Ex52, Guide1 Reverse | CRISPR guide | aaaccccgacgcatggttttcaccc |
| ACC1Ex52, Guide2 Forward | CRISPR guide | caccgcacactatttcattctacca |
| ACC1Ex52, Guide2 Reverse | CRISPR guide | aaactggtagaatgaaatagtgtgc |
| ACC1Ex52, Forward | Genotyping | ctggagccagaagggacagtag |
| ACC1Ex52, Reverse | Genotyping | ccccagcagcaactctagc |
| hGRAMD1A, Guide1 Forward | CRISPR guide | caccgtagctcctcagagaagggtg |
| hGRAMD1A, Guide1 Reverse | CRISPR guide | aaaccacccttctctgaggagctac |
| hGRAMD1A, Guide2 Forward | CRISPR guide | caccgagtgcgtgctgagatggagt |
| hGRAMD1A, Guide2 Reverse | CRISPR guide | aaacactccatctcagcacgcactc |
| hGRAMD1A, Forward | Genotyping | cacttctcttcctcttcctgccc |
| hGRAMD1A, Reverse | Genotyping | ccagtagccacaatgacagctg |
| hGRAMD1B, Guide1 Forward | CRISPR guide | caccgggacaagtccccgtccacac |
| hGRAMD1B, Guide1 Reverse | CRISPR guide | aaacgtgtggacggggacttgtccc |
| hGRAMD1B, Guide2 Forward | CRISPR guide | caccggagacagtcattatccagtg |
| hGRAMD1B, Guide2 Reverse | CRISPR guide | aaaccactggataat gactgtctcc |
| hGRAMD1B, Forward | Genotyping | tgccagtaactccaaccgcag |
| hGRAMD1B, Reverse | Genotyping | cagcccaccaagaatgcttagc |
| hGRAMD1C, Guide1 Forward | CRISPR guide | caccggggcgctccgactgtccgtc |
| hGRAMD1C, Guide1 Reverse | CRISPR guide | aaacgacggacagtcggagcgcccc |

| | | |
|---|---|---|
| hGRAMD1C, Guide2 Forward | CRISPR guide | caccggcgagccctttccctacggg |
| hGRAMD1C, Guide2 Reverse | CRISPR guide | aaaccccgtagggaaagggctcgcc |
| hGRAMD1C, Forward | Genotyping | tggaagtactcggagggccg |
| hGRAMD1C, Reverse | Genotyping | gaaaggaagggggcaaggc |
| ACAT1, Guide1 Forward | CRISPR guide | caccggttacttaaccatttgaagg |
| ACAT1, Guide1 Reverse | CRISPR guide | aaacccttcaaatggttaagtaacc |
| ACAT1, Guide2 Forward | CRISPR guide | caccgcagcatcattagataatggt |
| ACAT1, Guide2 Reverse | CRISPR guide | aaacaccattatctaatgatgctgc |
| ACAT1, Forward | Genotyping | ccctagagttcagcttgggcaac |
| ACAT1, Reverse | Genotyping | gatcctcatgcctcagcctcc |
| ACAT2, Guide1 Forward | CRISPR guide | caccggggcccgtctgcgtctgcag |
| ACAT2, Guide1 Reverse | CRISPR guide | aaacctgcagacgcagacgggcccc |
| ACAT2, Guide2 Forward | CRISPR guide | caccggctactccattcactatctg |
| ACAT2, Guide2 Reverse | CRISPR guide | aaaccagatagtgaatggagtagcc |
| ACAT2, Forward | Genotyping | gggacaagagctctacagggc |
| ACAT2, Reverse | Genotyping | gacccagaggcatcctgtgg |
| NPC1, Guide1 Forward | CRISPR guide | caccggcgctggacacagtagcagc |
| NPC1, Guide1 Reverse | CRISPR guide | aaacgctgctactgtgtccagcgcc |
| NPC1, Guide2 Forward | CRISPR guide | caccgttctgagcttgttccatctg |
| NPC1, Guide2 Reverse | CRISPR guide | aaaccagatggaacaagctcagaac |
| NPC1, Forward | Genotyping | ctgcgcggggtgctgaaacag |
| NPC1, Reverse | Genotyping | ggagttcgtgcgcatgtagcag |
| ATGL, Guide1 Forward | CRISPR guide | caccgttcctcggcgtctactacgt |
| ATGL, Guide1 Reverse | CRISPR guide | aaacacgtagtagacgccgaggaac |
| ATGL, Guide2 Forward | CRISPR guide | caccggcgagtcggtgggtaccgtg |
| ATGL, Guide2 Reverse | CRISPR guide | aaaccacggtacccaccgactcgcc |
| ATGL, Forward | Genotyping | gccgtgagtcccacacttaa |
| ATGL, Reverse | Genotyping | ccctcggagcaaaacaaacc |
| FASN, Guide1 Forward | CRISPR guide | caccgttctgggacaacctcatcgg |
| FASN, Guide1 Reverse | CRISPR guide | aaacccgatgaggttgtcccagaac |
| FASN, Guide2 Forward | CRISPR guide | caccgtcctggacattgcaccagag |
| FASN, Guide2 Reverse | CRISPR guide | aaacctctggtgcaatgtccaggac |
| FASN, Forward | Genotyping | ggatgtgtggggcactcacacc |
| FASN, Reverse | Genotyping | cgtaacaagcagatgggccctg |
| hDGAT1, Guide1 Forward | CRISPR guide | caccgcaaggacggagacgccggcg |
| hDGAT1, Guide1 Reverse | CRISPR guide | aaaccgccggcgtctccgtccttgc |
| hDGAT1, Guide2 Foward | CRISPR guide | caccgagtcacctgacaaggtcact |
| hDGAT1, Guide2 Reverse | CRISPR guide | aaacagtgaccttgtcaggtgactc |
| hDGAT1, Forward | Genotyping | ctgaggccatgggcgacc |
| hDGAT1, Reverse | Genotyping | aaggggacactgtccaggg |
| hDGAT2, Guide1 Foward | CRISPR guide | caccgcggagccagcgctctcacgg |
| hDGAT2, Guide1 Reverse | CRISPR guide | aaacccgtgagagcgctggctccgc |
| hDGAT2, Guide2 Foward | CRISPR guide | caccggcttgtttccctcatccgtg |
| hDGAT2, Guide2 Reverse | CRISPR guide | aaaccacggatgagggaaacaagcc |
| hDGAT2, Foward | Genotyping | ggcttcagccatgaagaccct |
| hDGAT2, Reverse | Genotyping | ccatcctgcccatccagtcta |
| hDIESL, Guide1 Foward | CRISPR guide | caccggctaaaatatttatacacaa |
| hDIESL, Guide1 Reverse | CRISPR guide | aaacttgtgtataaatattttagcc |
| hDIESL, Guide2 Foward | CRISPR guide | caccgtagggtgtgtataacactct |
| hDIESL, Guide2 Reverse | CRISPR guide | aaacagagtgttatacacaccctac |
| hDIESL, Foward | Genotyping | agactgagctaccaaccatcc |
| hDIESL, Reverse | Genotyping | cactgtgcccagccaagac |
| ACSL1, Guide1 Forward | CRISPR guide | caccgaaccagaccaaccctatgaa |

| | | |
|---|---|---|
| ACSL1, Guide1 Reverse | CRISPR guide | aaacttcatagggttggtctggttc |
| ACSL1, Guide2 Foward | CRISPR guide | caccgaagttaaggaatttaacagg |
| ACSL1, Guide2 Reverse | CRISPR guide | aaaccctgttaaattccttaacttc |
| ACSL3, Guide1 Forward | CRISPR guide | caccgttgatggaaagccaccgacc |
| ACSL3, Guide1 Reverse | CRISPR guide | aaacggtcggtggctttccatcaac |
| ACSL3, Guide2 Foward | CRISPR guide | caccgtttatggtgatagttttggg |
| ACSL3, Guide2 Reverse | CRISPR guide | aaaccccaaaactatcaccataaac |
| ACSL1, Foward | Genotyping | ccctctcctgccccacttaaac |
| ACSL1, Reverse | Genotyping | aataatgctatgccgccaggg |
| ACSL3, Foward | Genotyping | ggccaagggtacacacagtg |
| ACSL3, Reverse | Genotyping | aggcctccaggtttagccac |

### Supplementary Table 2. Antibodies used in this study

| Antibody | Application | Source | Identifier | Dilution |
|---|---|---|---|---|
| Anti-NPC1 (Rabbit polyclonal) | Western Blot | Proteintech | Cat# 13926-1-AP RRID:AB_2152050 | 1:1000 |
| Anti-P38 (Rabbit polyclonal) | Western Blot | Abcam | RRID: AB_306166 | 1:2000 |
| Anti-ACAT1 (Rabbit monoclonal, clone EPR26952-40) | Western Blot | Abcam | Cat# ab307597 RRID:AB_3750386 | 1:1000 |
| Anti-GAPDH (Mouse monoclonal, clone 1E6D9) | Western Blot | Proteintech | Cat# 60004-1-Ig RRID:AB_2107436 | 1:50,000 |
| Anti-ACC1 (Mouse monoclonal, clone 1A11G10) | Western Blot | Proteintech | Cat# 67373-1-Ig RRID:AB_2882621 | 1:10,000 |
| CoraLite Plus 647-conjugated ACC1 (Mouse Polyclonal) | Immunofluorescence | Proteintech | Cat# CL647-21923 RRID:AB_2934949 | 1:500 |
| Anti-ACC1 (phospho S79, Rabbit, monoclonal, clone EP1885Y) | Western Blot | Abcam | Cat#ab68191 RRID: AB_11156104 | 1:5000 |
| Anti-ATGL (Rabbit monoclonal, clone EPR19650) | Western Blot | Abcam | Cat#ab207799 RRID:AB_2888664 | 1:1000 |
| Anti-LDLR (Rabbit polyclonal) | Western Blot | Proteintech | Cat#10785-1-AP RRID:AB_2281164 | 1:2000 |
| Anti-Lamin A/C (Rabbit polyclonal) | Western Blot | Proteintech | Cat# 10298-1-AP RRID:AB_2296961 | 1:10,000 |

| | | | | |
|---|---|---|---|---|
| Anti-GFP (Mouse, monoclonal, clone 1E10H7) | Western Blot | Proteintech | Cat# 66002-1-Ig RRID:AB_11182611 | 1:20,000 |
| Anti-FASN (Rabbit polyclonal) | Western Blot | Proteintech | Cat# 10624-2-AP RRID:AB_2100801 | 1:5000 |
| Anti-Alpha Tubulin (Rabbit polyclonal) | Western Blot | Proteintech | Cat# 11224-1-AP RRID:AB_2210206 | 1:2000 |
| Anti-Rabbit IgG (H+L) (Goat polyclonal, HRP-conjugated) | Western Blot | Proteintech | Cat# SA00001-2 RRID:AB_2722564 | 1:10,000 |
| Anti-Mouse IgG (H+L) (Goat polyclonal, HRP-conjugated) | Western Blot | Proteintech | Cat# SA00001-1 RRID:AB_2722565 | 1:10,000 |
| Anti-AMPK alpha 1 (Mouse monoclonal) | Western Blot | Proteintech | Cat# 66536-1-Ig RRID: AB_2881899 | 1:1000 |
| Anti-ACC1 | Western Blot | Generated in Ref. 56 | N/A | 1:1000 |
| Anti-ACC2 | Western Blot | Generated in Ref. 56 | N/A | 1:1000 |
| Anti-β-Actin (Mouse monoclonal, clone AC-15) | Western Blot | Sigma | Cat# A5441 RRID:AB_476744 | 1:1000 |
